# Residue-level structural information from isotope-edited FTIR spectroscopy reveals distinct molecular structures of different amyloid-β42 oligomers

**DOI:** 10.1101/2025.03.31.646305

**Authors:** Faraz Vosough, Simon M. Carstensen, Cesare M. Baronio, Andreas Barth

## Abstract

Oligomers of the amyloid-β peptide 1-42 (Aβ42) are one of the underlying causes of pathogenicity in Alzheimer’s disease. However, investigating their molecular structure is challenging with common structural biology techniques. In this work we employed isotope-edited Fourier transform infrared spectroscopy with site-specific labeling, an advanced method enabling collection of residue-level structural information on oligomers of all sizes in aqueous solution. The Aβ42 peptides studied comprised those that were ^13^C, ^15^N-labeled at either residue V18, F20, A30, or I32, as well as those doubly-labeled at V18 and A30, at F20 and A30, and at F20 and I32. We studied three types of oligomers. For Aβ42 oligomers prepared in the absence of detergents, all labeled residues are positioned within β-strands. Moreover, in these oligomers intramolecular vibrational coupling was detected between residues V18 and A30, as well as F20 and A30, in addition to intermolecular coupling between V18 amides, and between F20 and I32 residues. This indicates that these residues are close in the 3-dimensional structure (contact). By contrast, detergent-stabilized oligomers formed in the presence of low concentrations of sodium dodecyl sulfate have different structures: V18 and F20 are not incorporated in a β-sheet, whereas A30 and I32 are found to reside in a β-sheet. No interactions (contacts) were detected between any pair of labeled residues in detergent-stabilized Aβ42 oligomers. We propose that in our oligomer preparations, V18 and F20 are stably incorporated into a β-sheet only for oligomers that are larger than dodecamers.

## Introduction

Amyloid-beta (Aβ) peptides are considered one of the major culprits in the pathophysiology of Alzheimer’s disease (AD).^1–3^ In particular, oligomers of Aβ peptides have become the subject of intense research in the last three decades, since growing evidence suggests that they are the most toxic Aβ aggregates involved in AD pathology.^4–12^

Despite the pivotal role of Aβ oligomers (AβOs) in AD and the increasing number of published reports on these neurotoxic species, experimental information on their atomic structure is yet largely scarce, particularly compared to the available structural details for Aβ fibrils.^13–22^ The structural data for AβOs are difficult to obtain with conventional structural biology techniques;^23^ major obstacles for structural studies of AβOs include their metastable and transient nature,^24,25^ as well as diversity in terms of size, shape and three-dimensional structure.^26,27^

Whereas the formation of a parallel, in-register β-sheet structure is well-documented in Aβ fibrils,^13,16,18,22,28^ experimental and computational studies have pointed to an (at least partially) antiparallel, intra- and/or intermolecular β-sheet architecture for AβOs.^29,30–39,40^ This is supported by the β-hairpin structure of a stabilized Aβ monomer, where the β-strands are formed by stretches of hydrophobic residues in the central hydrophobic cluster (L_17_VFFA_21_) and the C-terminal segment (mostly A_30_IIGLMV_36_).^29,41^ Also in the current models for AβOs, these residues constitute the stem of intramolecular hairpin-like structure(s) in the Aβ units within aggregates.^31,32,37,39^

So far, several more-detailed structural studies on AβOs have been reported, in most cases after stabilizing the oligomers by different biochemical methods such as preparation in the presence of detergents^31,39^, introduction of stabilizing disulfide bonds,^37,42^ inclusion in chimeric structures,^43,44^ and binding to antibodies.^45,46^ Ahmed *et al*^32^ reported a structural model of disc-shaped pentameric Aβ42Os in which three strands constitute an intramolecular antiparallel β-sheet but which does not contain intermolecular β-sheets. Lendel and colleagues^37^ proposed a hexameric β-barrel structure for protofibrillar assemblies of disulfide-stabilized Aβ42 peptide. The structure contains intra- and intermolecular antiparallel strands forming a barrel-like architecture. Solution NMR studies on small SDS-stabilized Aβ42Os (pre-globulomers) revealed a mixed β-structure, comprising intramolecular antiparallel and intermolecular parallel β-sheets.^31^ Also the first experimentally determined atomic structure of AβOs was obtained by solution NMR spectroscopy in 2020^39^ showing a pore-forming Aβ42 tetramer in a membrane-mimicking environment with a six-stranded, intra- and intermolecular antiparallel β-sheet.

The disparities between different molecular models of AβOs highlight the necessity for further structural studies. However, the present experimental methods for structural studies of amyloid proteins (mainly solid-state NMR) require relatively large amounts of sample^23,47^ and therefore are expensive to apply. In consequence, a comparison of different Aβ oligomer preparations is often not feasible. Fourier transform infrared (FTIR) spectroscopy provides a powerful alternative technique for structural analysis of protein aggregates.^36,48–51^ FTIR measurements are low-cost and quick and therefore compatible with the short-lived and metastable nature of oligomeric assemblies. Introduction of ^13^C and ^15^N-isotopes into the backbone at specific residues of the polypeptide sequence splits the amide I band in IR spectra for β-sheet structures, so that a new band is resolved at lower wavenumbers compared to the main ^12^C-band. This occurs due to the different vibrational frequencies of amide groups bearing ^12^C and ^13^C isotopes. The precise spectral position of the ^13^C-band depends on the electrostatic environment of the labeled residue(s) and the extent of the ^13^C-^13^C and/or ^12^C-^13^C coupling.^48^ A number of studies using isotope-edited FTIR spectroscopy on Aβ peptides have been reported, mostly on fibrillar structures of modified versions or fragments of the peptides.^52–57^ There are also a few reports of isotope-edited FTIR studies on AβOs^58,59^ using uniformly or segmentally labeled peptides. Our contribution was to establish that Aβ40 and Aβ42 mix (close to) randomly in the β-sheets of heterooligomers^60^ and that each peptide contributes at least two adjacent β-strands in homooligomers of either Aβ40 or Aβ42.^59,60^

In this work, we take advantage of combining site-specific labeling with FTIR spectroscopy to shed light on the local secondary structure as well as on intra/inter-molecular contacts.^48,50,52,61,62^ This information is similar to that obtained by solid-state NMR, but will help develop novel molecular models for different types of AβOs in aqueous solution. We applied this approach to three different detergent-stabilized and detergent-free Aβ42Os characterized previously^63^ and find that oligomers with and without SDS have different structures.

## Methods

### Peptides and chemicals

Unlabeled Aβ42, as well as Aβ42 peptides with site-specifically ^13^C, ^15^N-labeled residues at either V18, F20, A30, or I32, or doubly-labeled at either F20 & A30, V18 & A30, or F20 & I32 were synthesized, purified and provided by JPT Peptide Technologies (Berlin, Germany). D_2_O was obtained from Cambridge Isotope Laboratories (Tewskbury, MA, USA). The peptides are named V18L, A30L-, V18A30L-Aβ42 and so on. Other chemicals were purchased from Merck (Germany), unless noted otherwise.

### Preparation of Aβ42 monomers

Solutions of Aβ42 monomers and homogeneous SDS-stabilized oligomers were prepared as reported earlier.^63^ In summary, Aβ42 was dissolved in DMSO at 4 mg/ml concentration and applied to a pre-equilibrated HiTrap Desalting column (GE Healthcare, USA) at room temperature. An alkaline solution of 5mM NaOH in D2O (pD>12.3) was used to equilibrate the column and to elute the peptide fractions. 100-120 µl fractions of the Aβ42 solution were collected on ice and the peptide concentration was determined by measuring the tyrosine absorbance at 280 nm^64^ using a NanoDrop instrument (Eppendorf, Germany).

### Preparation of SDS-stabilized oligomers from Aβ42 monomers

SDS-stabilized oligomers of Aβ42 were prepared by the method originally reported by Barghorn and colleagues,^65^ with some modifications.^63^ Monomeric solutions of Aβ42 (pure labeled, pure unlabeled or mixed) were incubated for 24 hours at 37°C with either 0.2% or 0.05% SDS in D_2_O-based phosphate-buffered saline (PBS) to produce small (∼20 kDa) and medium (∼60 kDa) AβOs,^65^ respectively. Solutions of monomeric and oligomeric Aβ42 were flash-frozen in liquid nitrogen after production and stored at - 80°C (monomers) or -20°C (oligomers) until being used.

### Time-resolved FTIR spectroscopy of detergent-free Aβ42Os formation

18-20 µl of 50 mM sodium phosphate buffer in D_2_O, pD 7.4 was laid at the center of a flat CaF_2_ window covered on the periphery with a 90-100 µm plastic spacer, which was smeared with vacuum grease on both sides. The buffer solvent was evaporated under vacuum. The same volume of ice-cold alkaline monomeric solutions of either unlabeled Aβ42, differently labeled Aβ42 peptides, or a mix of labeled and unlabeled Aβ42 at the molar ratio of either 1:1 or 1:3 was applied to the dried buffer on the CaF_2_ window and mixed thoroughly by repeated pipetting. A second flat window was added and pressed gently onto the spacer. The IR cuvette was assembled and mounted in the sample holder of a Tensor 37 FTIR spectrophotometer (Bruker Optics, Germany) already cooled down to 0°C. The instrument was equipped with a sample shutter which enabled alternate measurements of reference and sample spectra without opening the chamber’s lid. An assembled IR cuvette containing a solution of the experiment’s buffer was mounted in the reference position. Both holders were connected to an external water bath with a thermostat to control the temperature. The spectrophotometer was supplied with an HgCdTe detector cooled with liquid nitrogen. Throughout the measurements, the instrument was purged with CO_2_-depleted dry air. Spectra were recorded earliest 20-30 minutes after closing the chamber’s lid to minimize the water vapor content in the wavenumber range 1900-800 cm^-1^, at a resolution of 2 cm^-1^, with a zero-filling factor 2, and with a 6 mm aperture. 300 scans were collected for the reference and the sample spectra. The light intensity above 2200 cm^-1^ was blocked with a germanium filter, while a 25 µm cellulose filter did the same for light intensity below 1500 cm^-1^ in order to be able to increase the light intensity in the spectral range of interest.^66^ The instrument was programmed to record absorption spectra in 30 min intervals. The temperature was kept at 0°C for the initial 2-4 hours of the experiments, after which it was raised to 37°C. A typical aggregation experiment lasted for 16-18 hours. The oligomers at the end of such experiments are the largest oligomers in this study with an average molecular mass of ∼100 kDa.

The OPUS 5.5 software was used for programming the instrument, analysis of the recorded spectra and plotting the Figures for IR spectra. Second derivatives of the absorbance spectra were calculated with a smoothing range of 9 or 17 data points, corresponding to spectral ranges of 9 and 17 cm^-1^.

### Preparation of SDS-stabilized oligomers from detergent-free oligomers monitored by FTIR spectroscopy

Detergent-free F20L-Aβ42Os were prepared in low binding plastic reaction tubes by mixing equal volumes of monomeric peptide and 50 mM sodium phosphate buffer, pD 7.4, followed by about 16 hours of incubation at 37°C. After the incubation and to induce formation of SDS-stabilized oligomers, SDS was introduced into the prepared oligomer solution at a final concentration of 0.2% w/v. The solution was loaded between two CaF_2_ windows, the IR cuvette assembled and the sample studied with time-resolved FTIR spectroscopy for about 16 hours using the experimental settings detailed above.

### FTIR spectroscopy of the SDS-stabilized oligomers

Transmission IR spectra for solutions of SDS-stabilized Aβ42Os were acquired on the Tensor 37 instrument with the experimental settings detailed above. The measurements were done with fresh buffer samples for each experiment at room temperature.

Attenuated total reflection (ATR)-FTIR spectra for SDS-stabilized Aβ42 oligomers were recorded on a Vertex 70 FTIR spectrophotometer equipped with an HgCdTe detector cooled with liquid nitrogen. For each sample, 2-2.5 µL of the oligomer solution was spread on the surface of the diamond crystal of a SensIR 9 reflection accessory and the solvent was allowed to evaporate at room temperature. Spectra were obtained for the dry film with a resolution of 2 cm^-1^ and a zero-filling factor 2 in the spectral range 1900-800 cm^-1^. 200 scans were collected for the reference and the sample spectra at room temperature. During the measurements, the instrument was constantly purged with CO_2_-depleted dry air. Second derivatives of absorbance were calculated with OPUS 5.5 using a smoothing range of 13 data points.

### Calculation of the amide I spectrum of model structures

Model structures of antiparallel and parallel β-sheets were created from building blocks consisting of two adjacent strands with two Ala residues each using published coordinates.^67,68^ They are shown in the Supplementary Information (SI). A number of the generated structures were manually edited in order to delete some of the generated residues. In addition, we used the experimentally determined β-barrel structure of α-hemolysin (residues 120-138, pdb code 7AHL),^69^ which contains structural features that resemble those of AβOs.^70^ Each amide group in these structures is assigned a number which is equal to the residue number of its oxygen atom.

A further model combined the structure of an α-helix obtained from a previous density functional theory calculation^71^ with that of an antiparallel β-sheet. The sheet had 8 strands and 6 residues per strand (5 amide groups) and the helix had 22 residues (21 amide groups). The helix was manually positioned in ChimeraX in a grove formed by the Ala side chains of the β-sheet at a distance that was slightly larger than van der Waals contact. The shortest distance between the backbone atoms in the helix and the sheet was 5.9 Å, while the distance between the closest C=O groups was ∼7.0.

The amide I spectra of these model structures were calculated with our Matlab program,^71,72^ which is based on the floating oscillator mechanism.^73–77^ The program diagonalizes a matrix of mass-normalized force constants. The diagonal elements of this *F*-matrix represent the intrinsic or local wavenumbers of the individual amide I oscillators. They depend on the molecular environment and were calculated using the electrostatic model 3F_ONH_ that relates the electric field at the O, N, and H atom of the considered amide group to its local wavenumber.^78^ The electric field was generated by charges on amide groups other than the considered one using charges of 0.40, -0.55, 0, and 0.15 elementary charges on the C, O, N, and H atoms, respectively. It caused a shift of the local wavenumber of the considered group, which was added to our reference wavenumber of 1707 cm^−1^.

The non-diagonal elements describe the vibrational coupling between the amide I oscillators of different amide groups. For nearest neighbors, these are taken from quantum chemical calculations^79^, whereas the other non-diagonal elements were calculated from transition dipole coupling^80–82^ using our most recent parameters that consider the dependence of the dipole derivative magnitude on the electrostatic environment (model DD(F_CO_)).^78^

The effect of ^13^C, ^15^N-labeling of specific residues was simulated by multiplying the diagonal elements of the *F*-matrix by a mass factor of 0.94725. The same factor was used for non-diagonal elements that represented an interaction between two labeled groups. Its square root was used for those non-diagonal elements that described an interaction between a labeled and an unlabeled group.^59^

Diagonalization of the *F*-matrix returns its eigenvalues and eigenvectors from which wavenumbers and dipole derivatives of the amide I normal mode can be calculated. The IR spectra were generated from Gaussian lines with 15 cm^−1^ full width at half maximum at the normal mode wavenumbers. They were normalized such that the integral of each component band was equal to the squared dipole derivative (in units of Debye^2^ u^-1^Å^-2^) of the respective normal mode, which is proportional to the dipole strength and thus to the integrated absorbance of a normal mode. The relative contribution of particular amide groups to the vibrational energy of a normal mode was calculating by squaring the normalized vibrational amplitude of the respective group.

## Results

### Terminology

This work used Aβ42 peptides in which one or two residues were ^13^C, ^15^N-labeled. The labeled atoms reside in different amide groups. For example for labeled residue F20, the amide group between F19 and F20 is ^15^N-labeled and the amide group between F20 and A21 is ^13^C-labeled. Only ^13^C-labeling has a considerable influence on the amide I vibration^83,84^ and ^15^N-labeling is thus ignored in the following. For simplicity, we will denote an amide group according to the residue that contains its carbonyl group. For example, the amide group between F20 and A21 will be denoted as the amide group of F20.

^13^C-labeling influences the amide I vibrations and thus the IR spectrum. In the text we will often associate a spectral property like “band” or a vibrational mode with an isotope, for example in the term “^13^C-band”. This is meant to indicate that the specified isotope has a strong influence on the particular property, but we note that the other isotope may also influence these properties. For example, ^12^C-amide groups generally participate in the ^13^C-modes that give rise to the ^13^C-band at low wavenumbers.

### Simulation of isotope effects on the amide I spectrum of β-sheets

We calculated amide I spectra of model β-sheet structures and of the β-barrel structure of α-hemolysin in order to get insight into the effects of placing a labeled amide group at different positions into β-strands and β-sheets. Before discussing the isotope-effects, we present the spectra calculated for unlabeled β-sheet model structures. They reflect well-known features of antiparallel β-sheets: a main absorption band at low wavenumbers between 1630 and 1620 cm^−1^ and a distinct band at high-wavenumbers near 1680 cm^−1^ characteristic of the antiparallel orientation of adjacent strands.^36^ Spectra of 8-stranded sheets with 6 and 10 residues (5 and 9 amide groups, respectively) are shown as gray lines in Figure 1A-F. In our spectra of parallel β-sheets, a distinct high-wavenumber band is missing although an absorption tail is present in this region (see the Supplementary Data at https://doi.org/10.17045/sthlmuni.24321667) (private link until publication: https://figshare.com/s/56a3304bf1f89ef4a6dd).

**Figure 1.**
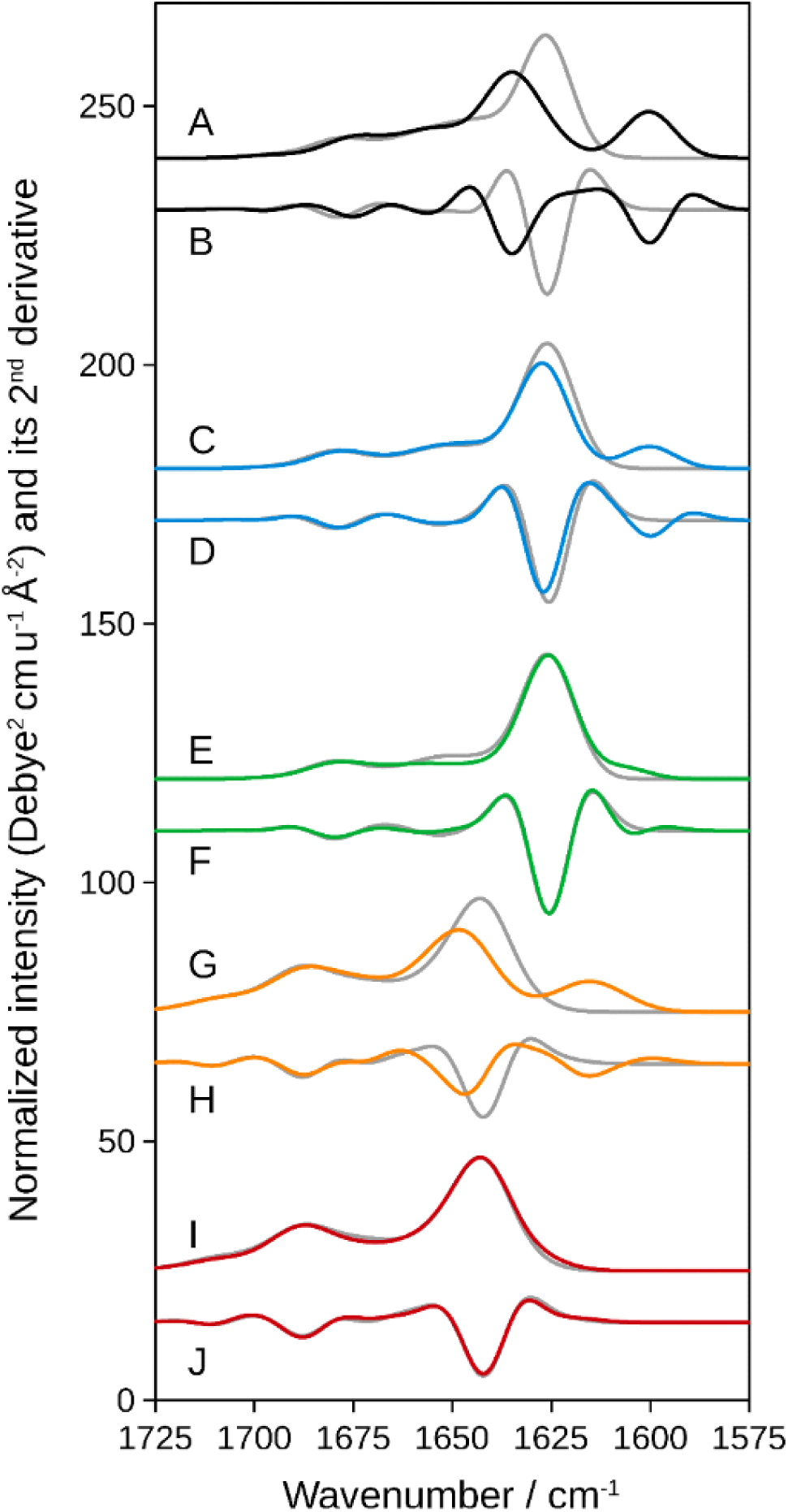
Calculated amide I spectra of β-sheet structures with and without ^13^C-isotopes. A, C, E, G, and I show normalized intensity spectra and B, D, F, H, and J their second derivatives. The colored curves are for the labeled structures and the gray curves for the unlabeled structures. A and B: 8-stranded antiparallel β-sheet with 6 residues per strand. Residues 3, 15, 27, and 39 in the middle of every second strand were labeled. C-F: 8-stranded antiparallel β-sheet with 10 residues per strand. Residues 5, 25, 45, and 65 in the middle of every second strand were labeled (C and D) or residues 1, 21, 41, and 61 at the C-terminus of every second strand (E and F). G-J: hemolysin β-barrel with a labels at position 122 (G and H) or 129 (I and J) in each β-hairpin.

The isotope-edited spectra were calculated for different positions of a single isotopically-labeled amide group per peptide unit. For most antiparallel β-sheet models, the peptide unit consisted of two adjacent strands because of the suggested β-hairpin structure of individual Aβ molecules. Such a labeling scheme has not been considered in previous spectrum simulations. Upon ^13^C-labeling of one amide group per peptide unit, a distinct ^13^C-band at lower wavenumber than the main ^12^C-band is observed for many label positions. The main ^12^C-band is often upshifted and its intensity reduced by labeling. Examples are shown in Figure 1A-D for the 8-stranded antiparallel β-sheet models and in Figure 1G-H for the α-hemolysin β-barrel. Labels at the edge or outside of β-sheets produced only a shoulder or hardly any change at all. For example, labeling the first residue in every second strand in an 8-stranded β-sheet with 10 residues generates a clear shoulder in Figure 1E, which gives a weak minimum in the second derivative spectrum in Figure 1F. An example for very minor changes is illustrated in Figure 1I-J, which shows the effects of labeling residue 129 in the loop of the hemolysin β-hairpins, the structure of which is shown in Figure S5 of the Supplementary Information (SI).

The results are discussed in detail in the SI and the spectra and normal mode information are contained in the Supplementary Data (https://doi.org/10.17045/sthlmuni.24321667) (private link until publication: https://figshare.com/s/56a3304bf1f89ef4a6dd). Here instead, we focus on those spectral features that provide insight into the probed antiparallel β-sheet structure: the position of the ^13^C-band, the shift of the main ^12^C-band caused by the presence of the ^13^C-groups and the relative intensities of the ^13^C- and of the main ^12^C-band. The dependence of these properties on the label positions can be summarized in the following guidelines for the interpretation of isotope-edited spectra of antiparallel β-sheets with a label in every second strand:

(i) A ^13^C-shoulder or ^13^C-band on the low-wavenumber side of the main ^12^C-band of β-sheets is observed when the labeled group is hydrogen bonded and resides in a β-sheet. Such features are not observed when the label is outside a β-sheet, for example in a loop region of a β-hairpin or in a close-by α-helix.
(ii) The spectral position of the ^13^C-band reflects the local wavenumber of the labeled amide group for sheets of 8 and more strands. For such sheets, it is less affected by the number of strands and very little by the number of residues per strand. The band is downshifted with respect to the local wavenumber of the labeled groups by ∼8 cm^-1^ due to coupling with other ^13^C-and^12^C-groups.
(iii) The main ^12^C-band shifts up upon labeling except for label positions at the strand ends. A strong upshift indicates central label positions in a narrow sheet (4-5 amide groups per β-strand, 8 strands). Up to ∼10 cm^−1^ upshift was observed for labels in every second strand in our simulations of such sheets and up to 6 cm^−1^ even for a single label. A smaller upshift indicates a peripheral label position or a wider sheet (more residues per strand).
(iv) The intensity of the ^13^C-band is rather constant for different label positions in regular β-sheets, where the local wavenumbers of the labeled groups are similar. However, when they are different due to structural heterogeneity or mobility, the ^13^C-band intensity decreases with increasing local wavenumber variation (see the hemolysin calculations in SI). In all cases, the ^13^C-band intensity is larger than expected from the number of labeled groups.^52,61,62^
(v) The intensity of the main ^12^C-band drops when labels are introduced in antiparallel β-sheets. The intensity drop is particularly large for label positions at or close to the strand end which cause only small or no ^12^C-band shifts. As a consequence, groups at the periphery of the β-strands can be diagnosed when a ^13^C-band is observed and the ^12^C-band is reduced but not shifted relative to the completely unlabeled case. The average ^12^C-band intensity decreases more for β-sheets with fewer residues per strand.
(vi) The above mentioned intensity effects affect the relative ^13^C-band intensity, i.e., the intensity of the ^13^C-band relative to that of the main ^12^C-band. This is of practical value, since the comparison of band intensities in the same spectrum is more reliable than such comparisons between the spectrum of a labeled and that of an unlabeled sample. The relative ^13^C-band intensity is enhanced compared to the relative abundance of the labeled amide groups. For the average relative ^13^C-band intensity, the enhancement is a factor of 3 to 4 (relative intensity / relative abundance) in our simulations of regular β-sheets. Accordingly, a smaller number of amide groups per strand gives a larger relative intensity, which may be used to estimate the number of amide groups per strand. The enhancement is less when the labeled groups experience different hydrogen bonding strengths but it is larger when only a single amide group is labeled in the entire sheet.
(vii) The intensity of the high-wavenumber band of β-sheets is only little affected by labeling. It can therefore be used to normalize approximately spectra of unlabeled and labeled structures. Note that a normalization using second derivative values needs to take into account the bandwidth because it strongly influences the second derivative value at the band position.

### Time-resolved Aβ42 aggregation experiments with detergent-freeAβ42Os

Aβ42 peptide aggregation and formation of oligomers from site-specifically labeled peptides was induced by a sudden drop of pD from over 12 to 7.4 in D_2_O-based 50 mM phosphate buffer. As in the similar experiments previously conducted on the unlabeled Aβ42 peptide,^63^ the aggregation process was monitored by following the position of the β-sheet main ^12^C-band in the amide I’ region of the IR spectra recorded in 30-minute intervals. The temperature was 0°C during the initial 2-4 hours to observe the early oligomers and 37°C in the following 14-17 hours. The average molecular mass of the oligomers at the end of such experiments is ∼100 kDa.

Figure 2 shows the second derivatives of the IR absorption in the amide I’ spectral range, where absorption peaks show up as negative bands, for unlabeled, singly- and doubly-labeled Aβ42. The band positions after 16 hours of aggregation at 37°C are reported in Table 1.

**Figure 2.**
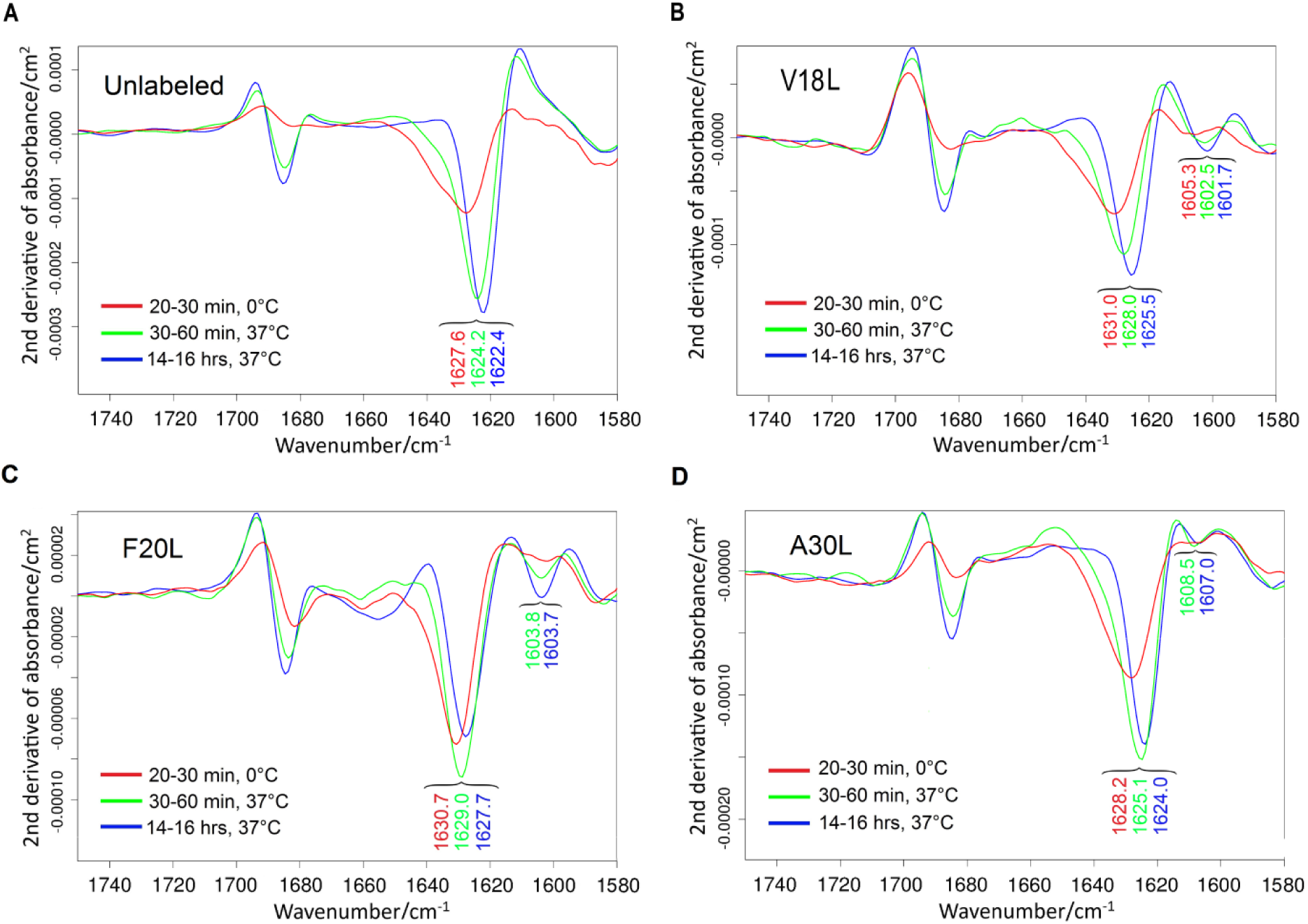
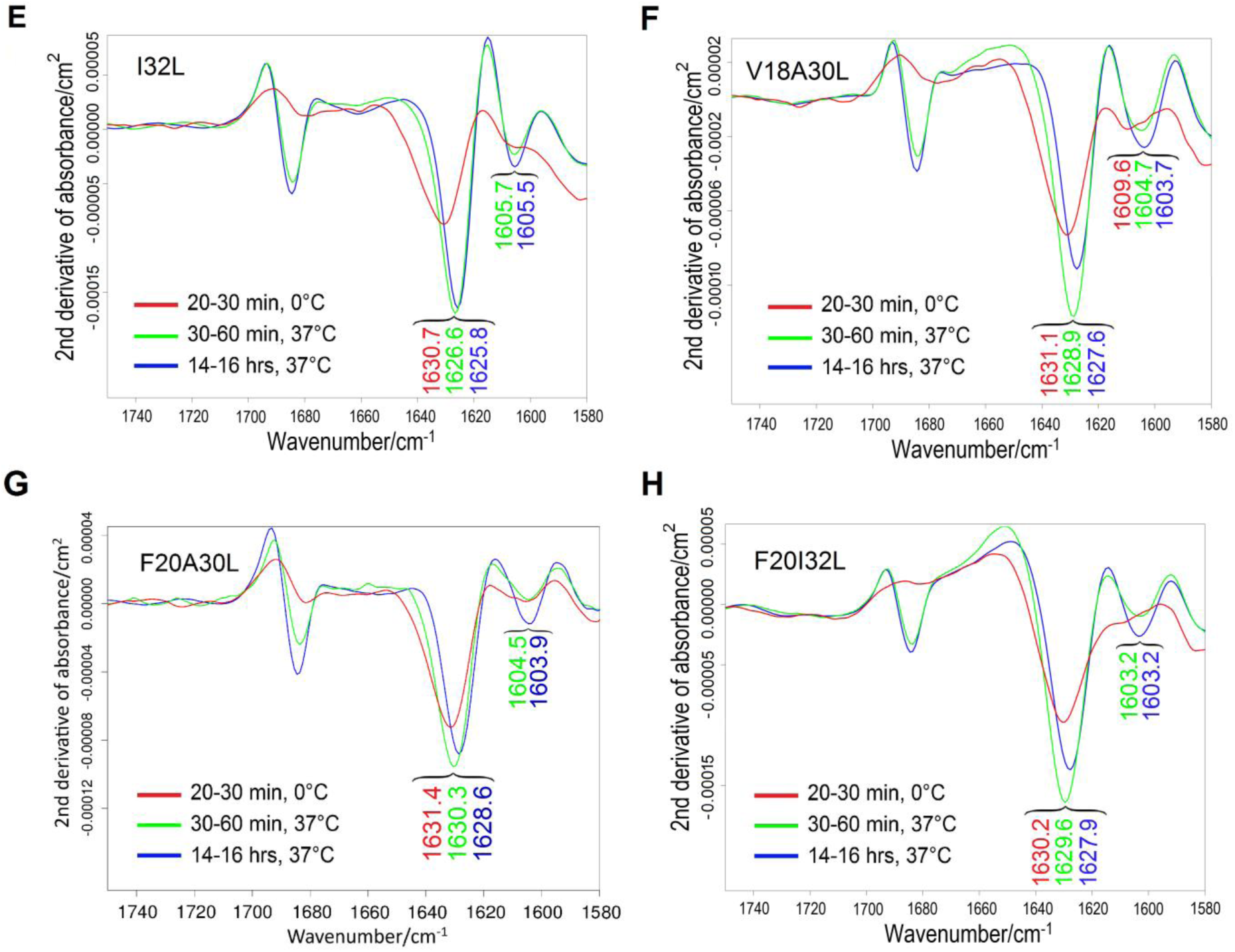
Second derivatives of IR absorbance spectra (calculated with smoothing range 17) recorded at three different time-points during the time-resolved aggregation of Aβ42 peptide variants, comprising unlabeled (panel A), V18L- (panel B), F20L- (panel C), A30L- (panel D), I32L- (panel E), V18A30L- (panel F), F20A30L- (panel G) and F20I32L- (Panel H) Aβ42. Red, green and blue spectra were recorded in the following time-points, respectively: 20-30 min after the pD drop and immediate incubation at 0°C, 30-60 min after temperature rise to 37°C, and after 14-16 hours of aggregation at 37°C. Spectra in panels A and H are averages from two independent experiments, whereas the other spectra were averaged from three experiments. None of the spectra was normalized.

**Table 1.** Spectral features in the amide I’ region of the IR second derivative spectra (calculated with soothing range 17) for the detergent-free and SDS-stabilized Aβ42Os of the unlabeled and site-specifically labeled peptides. The values for the detergent-free Aβ42Os were obtained after 16-18 hours of aggregation at 37°C.

| Aβ42 peptide/oligomer | $^{13}\text{C}$-band<sup>a</sup> | Main $^{12}\text{C}$-band<sup>a</sup> | High-wavenumber $^{12}\text{C}$-band<sup>a</sup> | $^{12}\text{C}$-band intensity ratio<sup>b</sup> |
| --- | --- | --- | --- | --- |
| **Detergent-free Aβ42O** (transmission spectra in D<sub>2</sub>O) |  |  |  |  |
| Unlabeled (n<sup>c</sup>=2) | ----- | 1622.4 ± 0.1<sup>b</sup> | 1685.3 ± 0.1 | 3.6 ± 0.1 |
| V18L (n=3) | 1601.9 ± 0.4 | 1625.4 ± 0.2 | 1684.8 ± 0.2 | 1.9 ± 0.1 |
| F20L (n=3) | 1603.7 ± 0.6 | 1627.7 ± 0.3 | 1684.5 ± 0.1 | 2.4 ± 0.5 |
| A30L (n=3) | 1607.0 ± 0.3 | 1624.0 ± 0.1 | 1685.0 ± 0.2 | 2.6 ± 0.1 |
| I32L (n=3) | 1605.5 ± 0.6 | 1625.8 ± 0.1 | 1684.7 ± 0.1 | 3.3 ± 0.6 |
| V18A30L (n=3) | 1603.7 ± 0.6 | 1627.6 ± 0.3 | 1684.5 ± 0.2 | 2.3 ± 0.1 |
| F20A30L (n=3) | 1603.9 ± 0.1 | 1628.6 ± 0.3 | 1684.2 ± 0.1 | 2.2 ± 0.3 |
| F20I32L (n=2) | 1603.2 ± 0.5 | 1627.9 ± 0.1 | 1684.2 ± 0.1 | 3.4 ± 0.1 |
| **SDS-stabilized Aβ42O** (transmission spectra in D<sub>2</sub>O) |  |  |  |  |
| Unlabeled (Aβ42O<sub>0.2%SDS</sub>) (n=2) | ----- | 1630.4 ± 0.5 | 1685.7 ± 0.3 | 10.7 ± 3.4 |
| V18L (Aβ42O<sub>0.2%SDS</sub>) (n=2) | N.A. | 1631.4 ± 0.0 | 1685.3 ± 0.1 | 3.7 ± 0.3 |
| F20L (Aβ42O<sub>0.2%SDS</sub>) (n=3) | N.A. | 1630.5 ± 0.2 | 1686.0 ± 0.6 | 6.0 ± 2.0 |
| A30L (Aβ42O<sub>0.2%SDS</sub>) (n=3) | 1609.5 ± 0.1 | 1631.3 ± 0.2 | 1685.3 ± 0.2 | 4.2 ± 1.4 |
| I32L (Aβ42O<sub>0.2%SDS</sub>) (n=3) | 1609.3 ± 0.4 | 1631.2 ± 0.2 | 1684.6 ± 0.2 | 2.6 ± 0.4 |
| V18A30L (Aβ42O<sub>0.2%SDS</sub>) (n=1) | 1610.4 | 1633.0 | 1684.7 | 3.2 |
| F20A30L (Aβ42O<sub>0.2%SDS</sub>) (n=2) | 1609.7 ± 0.4 | 1631.4 ± 0.4 | 1685.1 ± 0.1 | 3.4 ± 0.5 |
| F20I32L (Aβ42O<sub>0.2%SDS</sub>) (n=2) | 1609.1 ± 0.6 | 1631.4 ± 0.4 | 1684.3 ± 0.2 | 3.3 ± 0.4 |
| Unlabeled (Aβ42O<sub>0.05%SDS</sub>) (n=3) | ----- | 1629.6 ± 0.4 | 1686.1 ± 0.2 | 7.8 ± 1.5 |
| V18L (Aβ42O<sub>0.05%SDS</sub>) (n=2) | N.A. | 1630.5 ± 0.0 | 1685.9 ± 0.2 | 3.4 ± 0.1 |
| F20L (Aβ42O<sub>0.05%SDS</sub>) (n=2) | N.A. | 1629.2 ± 0.1 | 1686.7 ± 0.1 | 5.9 ± 2.7 |
| A30L (Aβ42O<sub>0.05%SDS</sub>) (n=3) | 1609.4 ± 0.2 | 1631.2 ± 0.4 | 1685.4 ± 0.4 | 3.9 ± 1.6 |
| I32L (Aβ42O<sub>0.05%SDS</sub>) (n=3) | 1606.9 ± 0.8 | 1631.0 ± 0.6 | 1684.7 ± 0.2 | 2.3 ± 0.1 |
| V18A30L (Aβ42O<sub>0.05%SDS</sub>) (n=1) | 1609.7 | 1632.4 | 1685.0 | 3.2 |
| F20A30L (Aβ42O<sub>0.05%SDS</sub>) (n=2) | 1609.6 ± 0.2 | 1631.2 ± 0.6 | 1685.3 ± 0.1 | 2.8 ± 0.1 |
| F20I32L (Aβ42O<sub>0.05%SDS</sub>) (n=2) | 1608.1 ± 0.3 | 1630.8 ± 0.1 | 1684.6 ± 0.1 | 2.4 ± 0.3 |
| **SDS-stabilized Aβ42O (ATR-FTIR spectra)** |  |  |  |  |
| Unlabeled (Aβ42O<sub>0.2%SDS</sub>) (n=2) | ----- | 1630.3 ± 0.1 | 1686.8 ± 0.3 | 2.5 ± 0.1 |
| F20L (Aβ42O<sub>0.2%SDS</sub>) (n=2) | 1607.7 ± 1.2 | 1631.1 ± 0.6 | 1686.9 ± 0.3 | 2.4 ± 0.1 |
| A30L (Aβ42O<sub>0.2%SDS</sub>) (n=1) | 1609.9 | 1631.6 | 1686.8 | 2.1 |
| F20A30L (Aβ42O<sub>0.2%SDS</sub>) (n=1) | 1609.2 | 1632.4 | 1686.2 | 1.8 |
| Unlabeled (Aβ42O<sub>0.05%SDS</sub>) (n=2) | ----- | 1628.2 ± 0.7 | 1687.1 ± 0.2 | 2.3 ± 0.1 |
| F20L (Aβ42O<sub>0.05%SDS</sub>) (n=2) | 1605.3 ± 0.4 | 1627.8 ± 0.0 | 1686.9 ± 0.2 | 2.2 ± 0.3 |
| A30L (Aβ42O<sub>0.05%SDS</sub>) (n=1) | 1609.7 | 1631.1 | 1686.7 | 1.6 |
| F20A30L (Aβ42O<sub>0.05%SDS</sub>) (n=1) | 1609.2 | 1631.7 | 1686.3 | 1.3 |

a) Average values and standard deviations were calculated for two or three independent experiments.
b) The <sup>12</sup>C-band intensity ratio was calculated by dividing the value of the second derivative at the position of the main <sup>12</sup>C-band by that of the high-wavenumber <sup>12</sup>C-band. This normalizes the signal of the main band, which is affected by labeling, to that of the high-wavenumber band, which is largely inert. c) n: number of experiments.

The β-sheet main band appears within 20-30 minutes from the induction of peptide aggregation at 0°C. In case of unlabeled Aβ42, its initial position is just below 1628 cm^-1^ (1627.6 cm^-1^ in Figure 2A). The raise in temperature from 0°C to 37°C accelerated the downshift of the main band to a final value of about 1622.4 cm^-1^. Peptide aggregation also led to the appearance and subsequent upshift of a high-wavenumber band, which finally stabilized at 1685.3 cm^-1^, in accordance with the antiparallel β-sheet structure of _Aβ42Os.30,36,85,86_

For the labeled peptides, the β-sheet main band resolves in the range 1631.4-1628.2 cm^-1^ at 0°C, depending on the labeled Aβ42 variant (Figures 2B-H). It shifts down during aggregation to wavenumbers between 1629 and 1624 cm^-1^, but the final band positions are higher than for the unlabeled peptide (about 1622.4 cm^-1^, Figure 2A), in line with the expected isotope effect on the main band of the β-sheet.^52,61^ The intensity of this β-sheet main band relative to the high-wavenumber band reduces upon labeling. In general, the position and intensity are most affected for the doubly-labeled peptides, which can be expected to experience the strongest isotope effects because the number of labeled groups is twice as large as for the singly-labeled peptides (Table 1). 30-60 minutes after raising the temperature to 37°C, the labeled peptides exhibit a relatively small ^13^C-band in the 1607-1602 cm^-1^ range, which grows over time for most labeled peptides but not for A30L-Aβ42 although the β-sheet content is still increasing as judged from the increase in the high-wavenumber band (Figure 2D). We note that the bandwidth of the high wavenumber band remains constant between the early and late spectra at 37°C (green and blue spectra of Figure 2D), which is a requirement for concluding band amplitudes in absorption spectra from second derivative spectra.

The positions of the β-sheet bands (^13^C-, main ^12^C- and high-wavenumber ^12^C-bands) at different stages of the time-resolved aggregation experiments are shown in Figure 3 for labeled and unlabeled Aβ42 peptides. Panel A shows band shifts already discussed above: the downshift of the β-sheet main band and the upshift of the high-wavenumber band for the unlabeled peptide. Similar shifts are also observed for the labeled peptides. In addition, panels B and D respectively show a downshift of the ^13^C-band for V18L- and A30L-Aβ42 between time-point “2” (30-60 minutes after the temperature raise to 37°C) and time- point “3” (the aggregation end-point). The downshift is also evident in the spectra shown in Figures 2B and 2D. This downshift indicates a conformational change of the oligomers as aggregation progresses and the oligomers grow in size. In contrast, there is no significant downshift of the ^13^C-band for F20L- and I32L-Aβ42 during aggregation (Figures 2C, 2E, 3C and 3E).

**Figure 3.**
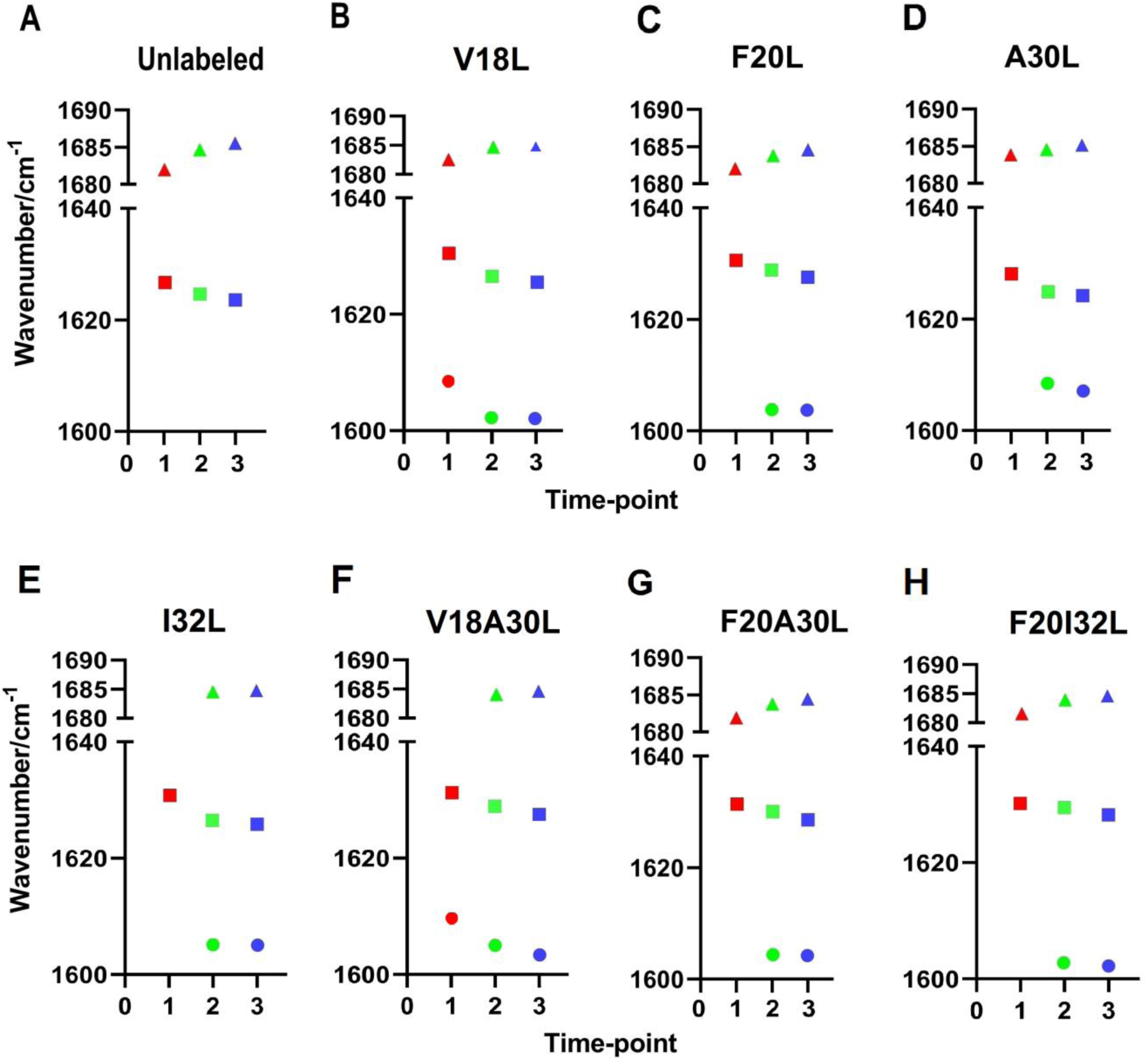
β-sheet band positions during the time-resolved aggregation of Aβ42 peptide variants, comprising unlabeled (panel A), V18L- (panel B), F20L- (panel C), A30L- (panel D), I32L- (panel E), V18A30L- (panel F), F20A30L- (panel G) and F20I32L- (Panel H) Aβ42. Time-points ‘’1’’, ‘’2’’ and ‘’3’’ indicate the following respective times: 20-30 min after the pD drop and immediate incubation at 0°C, 30-60 min after temperature rise to 37°C, and after 14-16 hours of aggregation at 37°C. The positions of the β-sheet main band are shown as squares, those of the ^13^C-mode band as circles and the high-wavenumber band positions as triangles.

### Backbone structure and environment of V18, F20, A30 and I32 in the late detergent-free Aβ42Os

Several conclusions can be drawn from the above-mentioned spectral features of singly-labeled peptides regarding the local conformation and the local environment of the labeled amide groups in the late Aβ42Os.

(i) The appearance of the ^13^C-band between 1607 and 1602cm^-1^, the upshift of the β-sheet main band and the relatively low intensity of this band (in relation to the high-wavenumber band at about 1685 cm^-1^, see Table 1) all indicate label positions within β-sheet structures.^52,61,62^ The labeled groups disrupt the coupling of ^12^C-carbonyl groups within and across β-strands, which leads to the observed effects according to our present calculations (see Supplementary Information) and previous ones by other groups.^62,87–89^ More specifically, the ^13^C-band positions of the singly-labeled peptides between 1607 and 1602 cm^-1^ are in agreement with a label location within out-of-register parallel β-sheets or a non-central position within antiparallel β-sheets. The second alternative – antiparallel β-sheets – is in line with the observation of the high-wavenumber band as discussed above. By contrast to the mentioned label locations, in-register label arrangements in parallel β-sheets or in the middle of antiparallel β-sheets give rise to considerably lower ^13^C-band positions around 1590 cm^-1^ ^88,90,91^ (see also the SI) because such a structure enables maximal vibrational coupling between the labeled groups.
(ii) The ^13^C-band wavenumber varies for labels on different positions; it is lowest for V18-Aβ42, and highest for A30L-Aβ42. Lower ^13^C-band positions indicate more favorable electrostatic interactions and/or more efficient coupling between the V18 ^13^C-amide groups or between V18 and ^12^C-groups (please refer to Methods for the naming of amide groups).
(iii) The relative ^13^C-band amplitude (relative to the main ^12^C-band) of V18L-Aβ42 is largest among the four residues studied, followed by that for the F20L-Aβ42. As for the upshift of the main ^12^C-band, F20L- Aβ42 shows the strongest effect. These observations are in line with fewer residues in the V18- and F20- containing β-strand than in the A30- and I32-containing β-strand. The above-mentioned results also indicate a more central position of F20 in its β-strand, or a more uniform hydrogen bonding strength of F20 in different peptide molecules.

### Conformational change between early and late detergent-free Aβ42Os

Further inspection of the singly-labeled peptide results indicates a different conformation for the early oligomers observed at 0°C and the late oligomers at the end of the aggregation experiment at 37°C:

(i) For V18L- and A30L-Aβ42, ^13^C-band appears to be present already from the beginning at 0°C. It is very clear in the first spectrum recorded at 37°C (green spectra in Figures 2B and 2D, respectively) indicating that V18 and A30 are part of the early β-sheets. While the band intensity of V18L-Aβ42 increases during further aggregation, that of A30L-Aβ42 remains rather constant. This is in contrast to the β-sheet content monitored by the high-wavenumber band. Thus, the ^13^C-band of A30L-Aβ42 decreases relative to the high-wavenumber band in the final phase of the experiment. This indicates a conformational change resulting in a larger number of residues in the A30-containing β-strand or an altered coupling to other amide groups due to a conformational change.
(ii) For F20L- and I32L-Aβ42, the ^13^C-band is not obvious at 0°C and is relatively small at the beginning of the aggregation at 37°C (green spectra in Figures 2C and 2E, respectively), whereas the high- and low-wavenumber ^12^C-bands of the β-sheet have considerable intensity. The ^13^C-band intensifies significantly as the late oligomers form. There are two explanations for the absence of a clear ^13^C-band in the early oligomers: (i) either F20 and I32 do not reside in a β-sheet, in which case our model calculations (see SI) indicate that there is no distinct ^13^C-band, or (ii) the F20 and I32 amide groups experience interaction strengths that vary in time or between different molecules, both of which will make the ^13^C-band broad and weak in the second derivative. Our observations indicate that the local environments of F20 and I32 are different in the early and late oligomers. Once the later oligomers are formed, the ^13^C-band position of F20L- and I32L-Aβ42 do not change significantly during further aggregation, indicating that the molecular environment of both amide groups is not altered after their incorporation into the β-sheets.
(iii) A change in the V18 and A30 local environment during aggregation can also be deduced from the downshift of the ^13^C-band of V18L- (Figures 2B and 3B) and A30L-Aβ42 (Figures 2D and 3D). The downshift can be explained by stronger favorable electrostatic interactions and/or a re-arrangement of strands which enables more efficient vibrational coupling of the V18 and A30 amide groups.
(iv) For F20L- and A30L-Aβ42, the intensity of the main band decreases during aggregation at 37°C while the β-sheet content increases as monitored by the increasing intensity of the high-wavenumber band (compare the green and the blue spectra in Figure 2). This is in contrast to the unlabeled peptide, where both bands increase throughout the experiment. The intensity decrease for the labeled peptides indicates a more pronounced isotope effect on the main band at the end of the aggregation experiment. Our calculations indicate that the intensity of the main band depends on the number of residues per strand and on the label position within the sheet. The intensity decrease can be explained by an elongation of β-strands during aggregation, or by a position shift of residues F20 and A30 in their respective β-strands.

### 3D contacts in detergent-free oligomers

The following section discusses whether the labeled amide groups are close to each other in the three-dimensional structure of the Aβ42Os. In order to investigate the nature (intra- or intermolecular) of such molecular contacts, isotope dilution experiments were conducted. Co-aggregation of a mixture of labeled and unlabeled Aβ42 peptides under the same conditions as for the pure peptides produces identical oligomeric assemblies in terms of molecular structure in which molecules of the unlabeled and labeled peptide are randomly positioned within the oligomers. Meanwhile, the ^13^C-band shifts due to isotope dilution can provide insights on the origin of the molecular contact(s).^50,52,88,92–94^ Dilution diminishes intermolecular coupling between ^13^C-groups but maintains the intramolecular couplings.

The results of isotope dilution experiments with different doubly- and singly-labeled peptides under various dilution ratios are listed in Table 2 and the interpretation summarized in Table 3. While a 1:3 dilution (labeled:unlabeled) would have been preferable for all peptide variants, it was not practical for some due to the small amplitude of their ^13^C-bands. The dilution experiments are discussed in the following, starting with the singly-labeled peptides. If not stated otherwise, the discussed band positions refer to those after 16-18 hours of aggregation at 37°C.

**Table 2.** The ^13^C-band positions (cm^-1^) in the amide I’ region of the second derivative IR spectra (calculated with smoothing ranges of either 9 or 17) for isotope dilution experiments on doubly- and singly-labeled peptides under various dilution ratios after 16-18 hours of co-aggregation at 37°C.

| Labeled variant | Pure labeled | D.R.<sup>a</sup> 1:1 | D.R. 1:2 | D.R. 1:3 | Average for respective singly-labeled variants |
| --- | --- | --- | --- | --- | --- |
| S.R.<sup>b</sup>9 |  |  |  |  |  |
| V18A30L (n=3<sup>c</sup>) | 1602.6 ± 1.4 / 1608.7 ± 0.6<sup>d</sup> | ---- | ---- | 1606.0 ± 1.0 | 1601.1/1607.4 |
| F20A30L (n=3) | 1604.1 ± 0.2 | 1603.9 ± 0.2 | ---- | 1603.1 ± 0.8 | 1604.9 |
| F20I32L (n=2) | 1602.8 ± 0.2 | 1605.5<sup>d</sup> | 1605.9 ± 1.7<sup>d</sup> | 1606.0 ± 0.1<sup>d</sup> | 1606<sup>d</sup> |
| V18L (n=2) | 1601.4 ± 0.2 | ---- | 1603.4 | 1605.8 | ---- |
| F20L (n=3) | 1603.6 ± 0.2 | 1604.0 ± 0.5 | ---- | ---- | ---- |
| I32L (n=3) | 1606.1 ± 0.5 | ---- | 1606.8 ± 0.3 | 1607.5 | ---- |
| S.R.17 |  |  |  |  |  |
| V18A30L (n=3) | 1603.7 ± 0.6 | ---- | ---- | 1604.9 ± 0.1 | 1603.1 |
| F20A30L (n=3) | 1603.9 ± 0.1 | n. d.<sup>e</sup> | ---- | n. d. | 1605.3 |
| F20I32L (n=2) | 1603.2 ± 0.5 | 1604.7 | 1604.1 ± 1.7 | 1603.9 ± 1.3 | 1604.6 |
| V18L (n=2) | 1601.9 ± 0.4 | ---- | n. d. | n. d. | ---- |
| F20L (n=3) | 1603.7 ± 0.6 | 1603.6 ± 0.0 | ---- | ---- | ---- |
| I32L (n=3) | 1605.5 ± 0.6 | ---- | 1605.3 ± 0.8 | 1606.0 | ---- |

<sup>a</sup> D.R.: dilution (molar) ratio. <sup>b</sup> S.R.: smoothing range in $\text{cm}^{-1}$ . <sup>c</sup>n: indicates number of repeats for the pure labeled peptide. <sup>d</sup>: these values are somewhat different from the ones shown in figures 4 and 5, since slightly different time ranges were averaged in each case. <sup>e</sup> n.d.: not determined (because the spectrum did not show a minimum for this smoothing range).

**Table 3.**
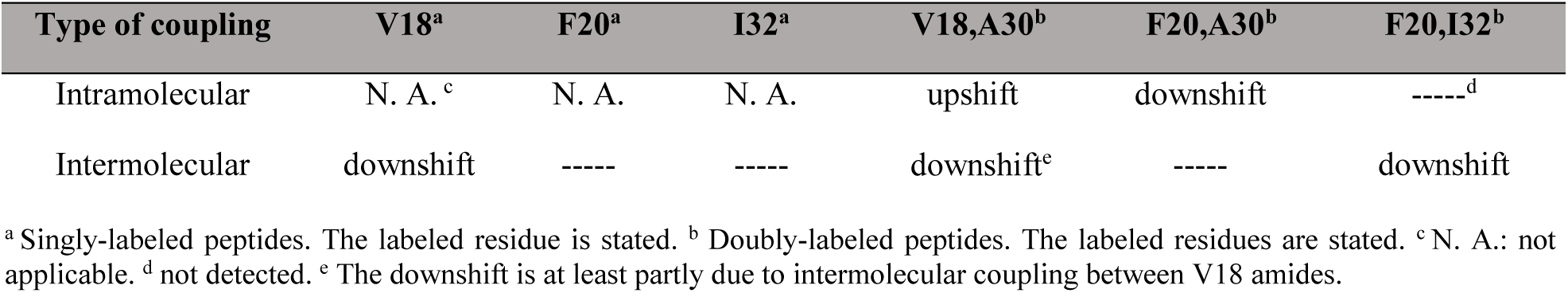
^13^C-band shifts due to intra- and intermolecular contacts in detergent-free oligomers after16-18 hours of co-aggregation at 37°C. The residue of the amide group’s carbonyl is given.

Isotope dilution experiments on V18L-Aβ42 revealed a lower ^13^C-band position for the pure peptide than for the 1:3 isotope-diluted peptide (1601.4 ± 0.2 cm^-1^ for the pure peptide, 1605.8 cm^-1^ after 1:3 isotope dilution, Table 2, smoothing range 9 cm^-1^). This downshift of 4.4 cm^-1^ upon isotope enrichment indicates intermolecular contacts between V18 amides in our detergent-free Aβ42Os. This shift indicates that V18 amide groups in neighboring peptide molecules are close to each other in the three-dimensional structure but does not imply that they need to reside in adjacent β-strands (see Supplementary Information).

For F20L-Aβ42, 1:1 isotope dilution did not produce a significant shift (Table 2), indicating the absence of strong intermolecular coupling between amide groups of F20. Isotope dilution experiments were not performed with A30-Aβ42 because of the small amplitude of the ^13^C-band.

For I32-Aβ42, isotope dilution produced a relatively small upshift of the ^13^C-band within the error limits (from 1606.1 ± 0.5 cm^-1^ to 1606.8 ± 0.3 cm^-1^ at 1:2 dilution, smoothing range 9, see Table 2). Thus, no intermolecular coupling was detected for the I32 amide group either.

Turning to doubly-labeled peptides, we first discuss shifts of the ^13^C-band upon isotope dilution and then compare the band position of the diluted sample to that of an average spectrum of the two respective singly-labeled peptides. A band shift upon dilution will reveal intermolecular coupling, while intramolecular coupling is detected when the band positions of the diluted sample and of the average spectrum are different. In the absence of intermolecular coupling, intramolecular coupling can also be detected by comparing the spectrum of the pure doubly-labeled peptide with the average spectrum of the singly-labeled peptides. Spectra of the pure doubly-labeled peptides are shown in Figure 2, compared to the average spectra of the respective singly-labeled peptides in Figure 4, and to isotope diluted spectra in Figure 5.

**Figure 4.**
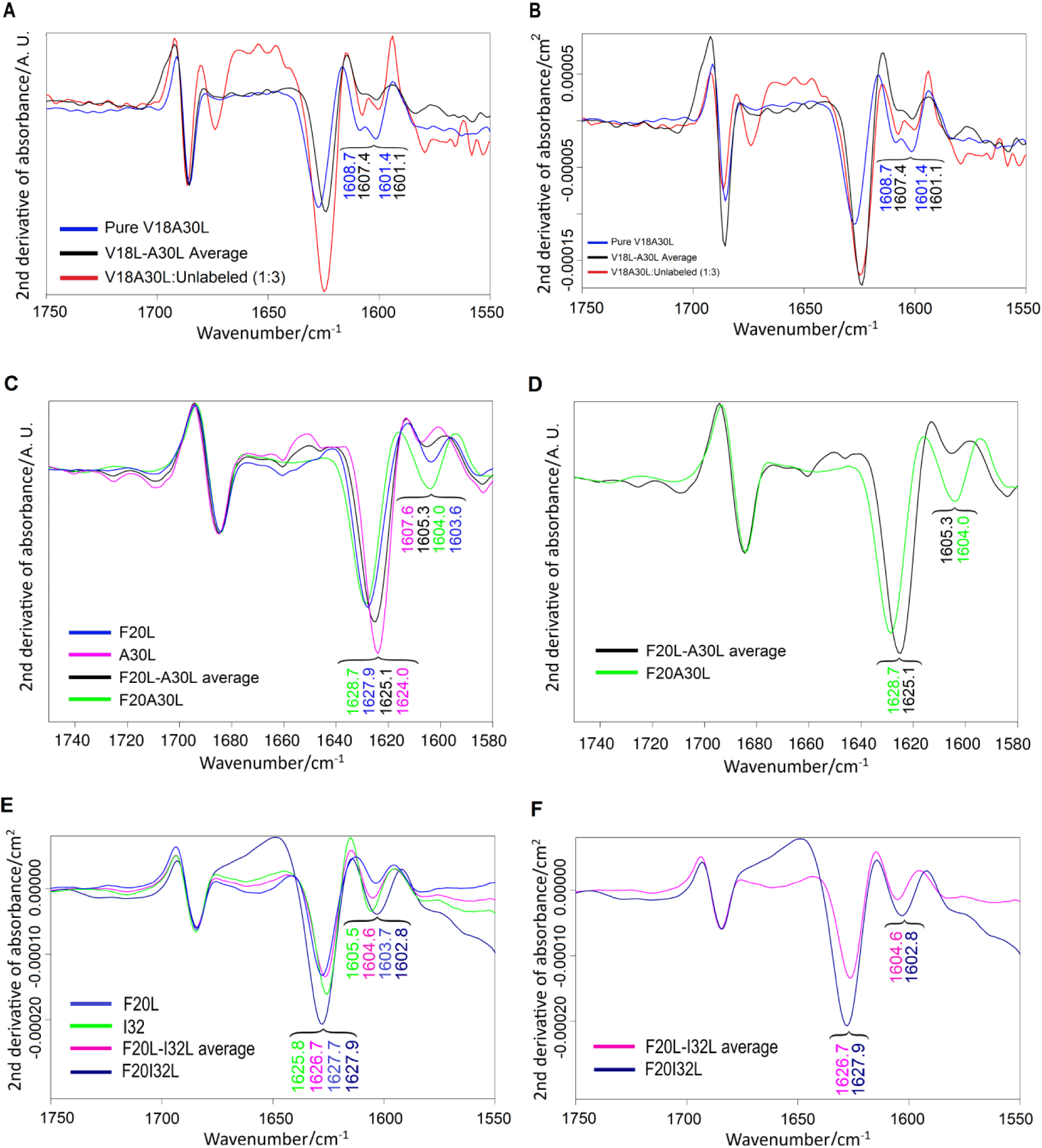
Second derivatives of IR absorbance spectra of the end-point aggregation state in the time-resolved experiments for: A-B) V18A30L-(blue) as well as the average of V18L- and A30L- (black) Aβ42. The red spectrum shows the 1:3 diluted peptide. Spectra in panel A were normalized on the high- wavenumber band, while those in panel B were not. Second derivatives were calculated with smoothing range 9. C-D) F20L- (blue), A30L- (purple), F20L- and A30L- average (black), and F20A30L- (green) Aβ42; normalized on the high-wavenumber band. E-F) F20L- (blue), I32L- (green), F20L- and I32L- average (purple), and F20A30L- (darkblue) Aβ42. Spectra in panels E-F were not normalized. Panels C and E depict all four spectra, while in panels D and F the spectra for single-position labeled peptides are deleted for a better comparison. Second derivatives in panels C-F were calculated with smoothing range 17.

**Figure 5.**
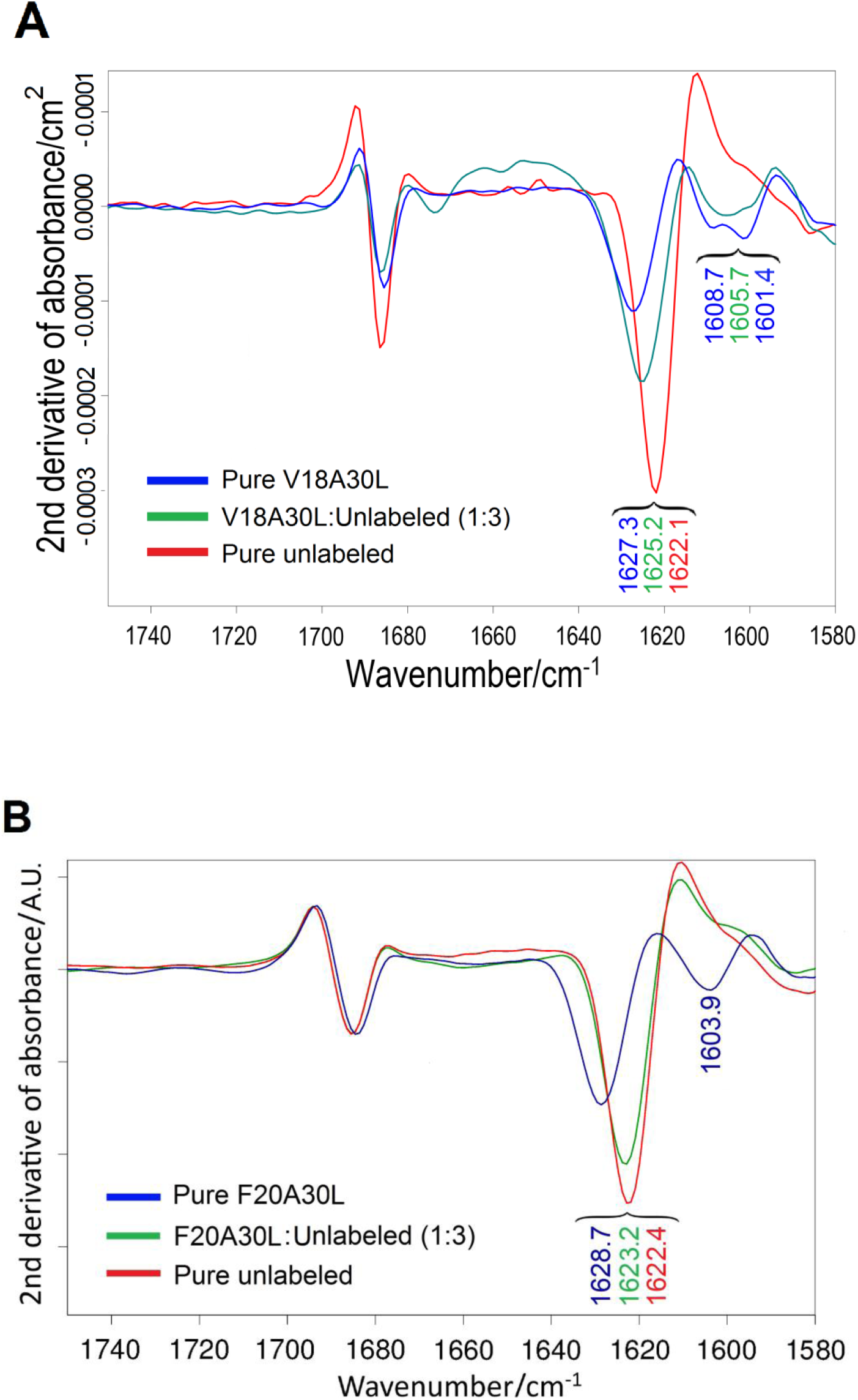
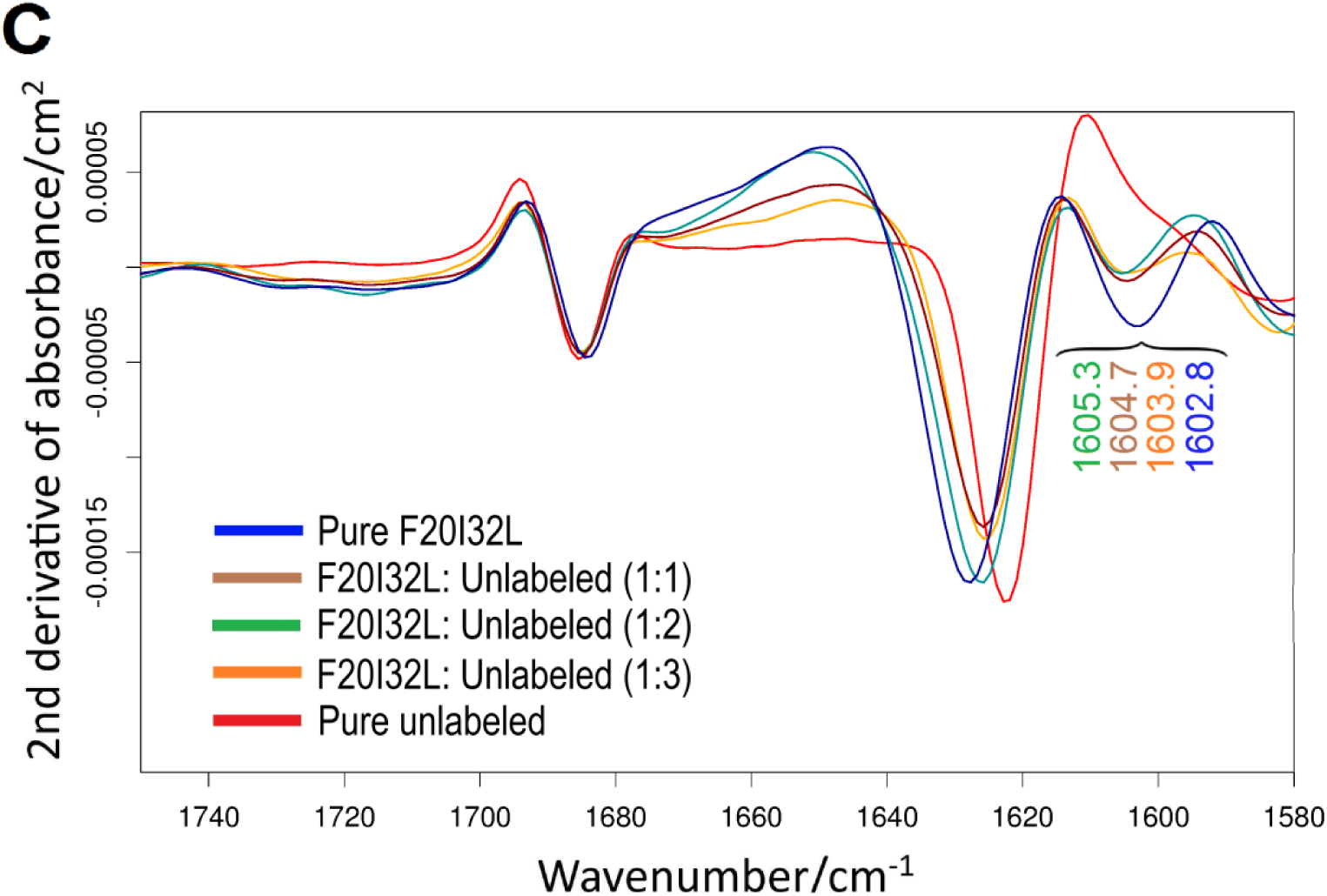
Second derivatives of IR absorbance spectra for the isotope dilution experiment with A) V18A30L-, B) F20A30L-, and C) F20I32L-Aβ42. The doubly-labeled peptide was mixed with unlabeled Aβ42 in the molar ratio 1:3 (A,B) and either 1:1, 1:2 or 1:3 (C). Each spectrum shows the respective end- point (after 16-18 hours of aggregation) of the time-resolved aggregation experiment for the pure labeled (blue), the 1:3 molar ratio mixture of the labeled and unlabeled peptide, and the unlabeled (red) Aβ42. The second derivative of IR spectra were calculated with smoothing range of 9 for spectra in panel A and 17 for those in panels B and C. Spectra in panel B were normalized on the high-wavenumber ^12^C-band.

For V18A30L-Aβ42, the ^13^C-band is broad and seems to be composed of two component bands (Figures 4A-B, once evaluated in second derivative spectra with smoothing range 9). The weight of the band is on the low-wavenumber side near 1602 cm^-1^ for the pure peptide but on the high-wavenumber side near 1608 cm^-1^ for the 1:3 diluted peptide (Figures 4A-B and 5A). Thus, the ^13^C-band shifts down upon isotope enrichment, which indicates intermolecular coupling. A downshift upon isotope enrichment was also observed for the singly-labeled V18L-Aβ42, which implies that at least part of the downshift for the doubly-labeled peptide is caused by coupling between the V18 amides in different molecules. In contrast to the isotope-diluted spectrum, the average spectrum of the singly-labeled peptides V18L- and A30L- Aβ42 has the weight of the ^13^C-band on the low-wavenumber side between1602 and 1601 cm^-1^. Thus, incorporating a second labeled amide group into the labeled peptides produces an upshift of the ^13^C-band due to intramolecular coupling.

For F20A30L-Aβ42, isotope dilutions of 1:1 and 1:3 were employed. The ^13^C-band develops after the temperature raise to 37°C and its position remains unchanged throughout the aggregation process in the second derivative spectra. Figure 5B depicts the second derivative spectra for the pure unlabeled Aβ42, pure F20A30L-Aβ42 and the 1:3 mixture of the two after 16-18 hours of aggregation. The 1:1 dilution gives a well-defined ^13^C-band position with less error than the 1:3 dilution, and both positions are unchanged with respect to that of the pure peptide within the error limits. Therefore, we can exclude considerable intermolecular coupling between the F20 and A30 amide groups of Aβ42.

However, F20A30L-Aβ42 exhibits intramolecular coupling between the labeled amide groups. As the ^13^C-band position is not affected by dilution, we can compare it directly to the calculated average spectrum of F20L- and A30L-Aβ42 (Figures 4C-D, smoothing range of 17). The average shows the ^13^C- band at 1605.3 cm^-1^, while it is resolved at 1604.0 cm^-1^ for F20A30L-Aβ42 (Figures 4C-D, green spectrum). Because of the absence of intermolecular coupling, this difference indicates a downshift of the ^13^C-band due to intramolecular coupling between the F20 and A30 amide groups, which entails relative spatial proximity of the two amide groups of the same molecule in our late detergent-free Aβ42Os.

For F20I32L-Aβ42, the pure peptide has a significantly lower ^13^C-band position than in the isotope diluted mixture (1602.8 ± 0.2, 1605.5, 1605.9 ± 1.7 and 1606.0 ± 0.1 for pure labeled, and 1:1, 1:2, and 1:3 diluted peptide, respectively, Table 2, smoothing range 9). These data clearly indicate a downshift of the ^13^C-band due to intermolecular coupling. Since we did not detect intermolecular coupling for the respective singly-labeled peptides, the coupling must be between F20 and I32 amide groups in different molecules of the peptide. The ^13^C-band position in the average spectrum of the singly-labeled peptide variants is similar to that of the diluted doubly-labeled peptide, ruling out strong intramolecular coupling between F20 and I32. The results are also depicted in Figure 5C.

#### FTIR spectroscopy of SDS-stabilized Aβ42Os

SDS-stabilized oligomers were prepared by incubating the labeled peptide variants in the presence of 0.2% and 0.05% of SDS in a high ionic strength (containing 140 mM NaCl) buffer for 24 hours, which yielded predominantly tetrameric small SDS-Aβ42Os and - mainly dodecameric - medium SDS-Aβ42Os, respectively.^63^ Figure 6 presents the second derivatives of IR absorption spectra for these oligomers. The high-wavenumber ^12^C-band was used to normalize the IR spectra for comparisons, as it is only little affected by ^13^C-labeling according to our calculations. As shown in Figure 6 and listed in Table 1, the spectral positions of the β-sheet main band for unlabeled Aβ42 are different for the two SDS-Aβ42Os and higher than for the detergent-free Aβ42Os according to the different oligomer sizes.^63^

**Figure 6.**
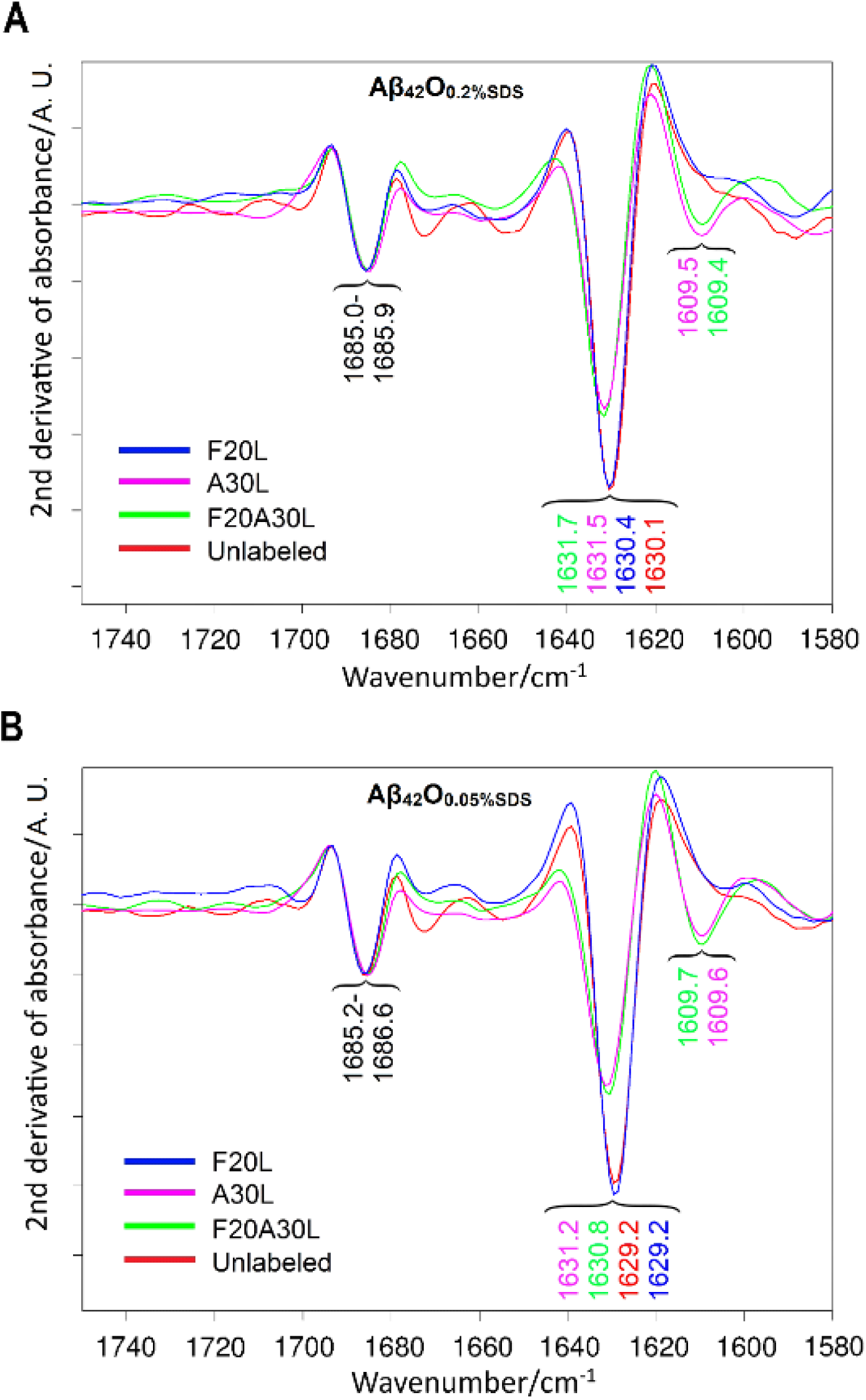
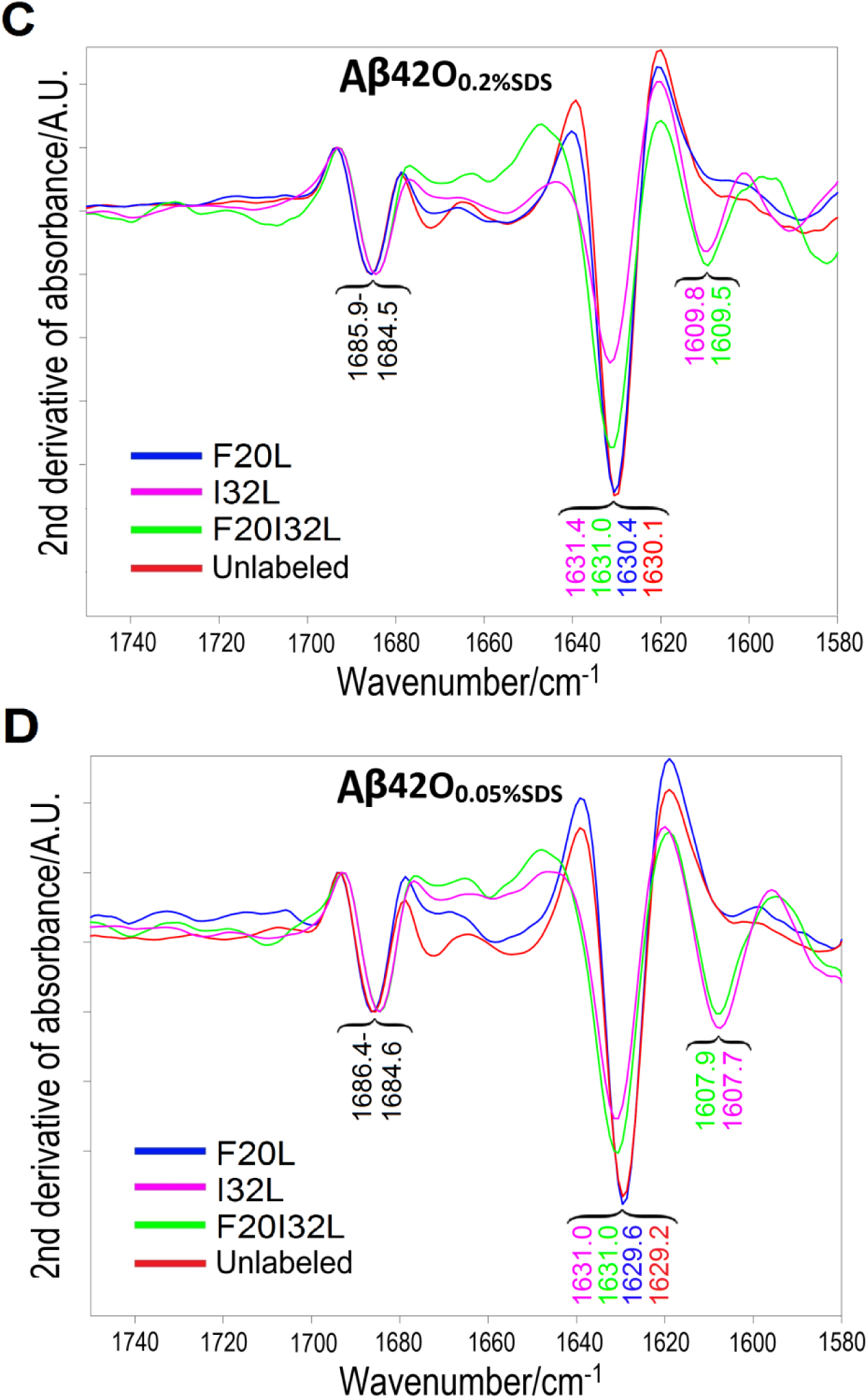
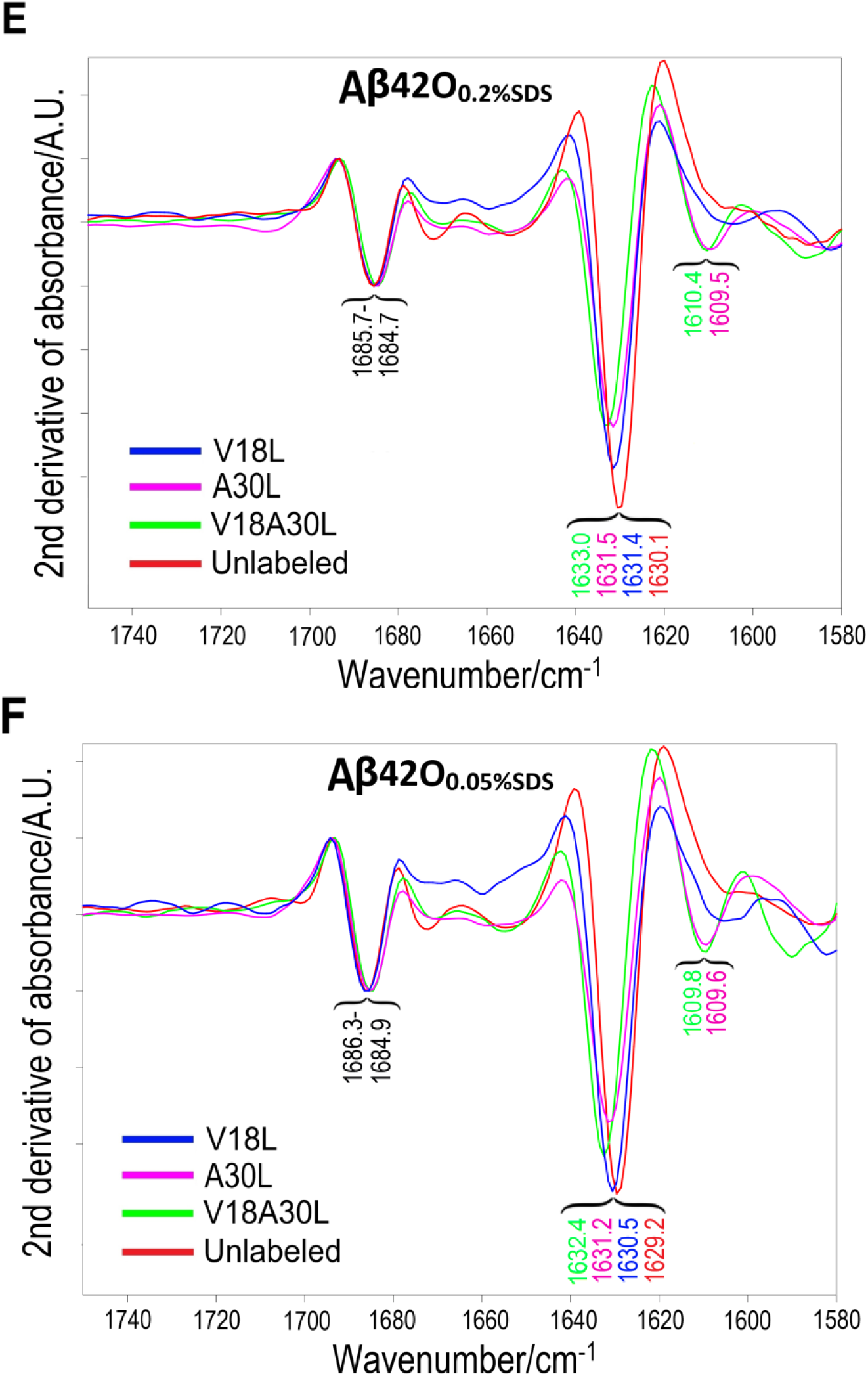
Second derivatives of IR absorbance spectra (calculated with smoothing range 17) for SDS-stabilized oligomers of the Aβ42 peptide labeled at residues: A-B) F20, A30 and F20A30; C-D) F20, I32 and F20I32; and E-F) V18, A30 and V18A30, as well as of the unlabeled peptide (in red). Panels A, C, E and B, D, F show the spectra for the 0.2% SDS-stabilized and 0.05% SDS-stabilized oligomers, respectively. The spectra were normalized on the high-wavenumber ^12^C-band.

No ^13^C-bands are detected for both types of V18L- and F20L- SDS-Aβ42Os (blue spectra) and the IR spectra approximately match those obtained for the unlabeled Aβ42Os (red spectra). However, we note a deeper indent near 1605 cm^-1^ in V18L and F20L spectra than in the spectrum of unlabeled Aβ42, which could indicate a weak ^13^C-band. In line with the absence of a clear ^13^C-band, the main ^12^C-band remains largely unaffected by F20 labeling, since neither its spectral position, nor its intensity changes. In case of V18 labeling, some moderate effects on the position and intensity of the main ^12^C-band are observed (Table 1 and Figures 6A-B and 6E-F). This indicates that the unlabeled V18 amide group participates in the normal mode(s) that generate the ^12^C β-sheet main band. When V18 is labeled, the coupling to the other β-sheet amide groups is disrupted, which explains the observed effects on the β-sheet main band. These observations suggest that V18 is close to amide groups within a β-sheet, while it may not be stably incorporated into it.

The IR spectra for SDS-Aβ42Os of A30L-, I32L-, V18A30L-, F20A30L- and F20I32L-Aβ42 are distinct from those for the unlabeled Aβ42Os in several aspects. Most remarkably is the clear appearance of the ^13^C-band in the range 1610.4-1606.9 cm^-1^. In addition, an upshift of about 0.5-2.5 cm^-1^ is observed for the β-sheet main band when compared to the unlabeled Aβ42. Lastly, the relative intensity of the β-sheet main band is significantly decreased relative to the high-wavenumber band for the SDS-Aβ42Os of the above-mentioned labeled variants compared to the same oligomers from the unlabeled peptide. Moreover, the near-perfect match of the IR spectra in case of same size SDS-Aβ42Os of A30L- and F20A30L-Aβ42 (Figure 6A-B) further confirms that there is very little contribution from the F20 residue to the IR spectral features in the amide I’ region. Similar, but less strong overlap, can be observed between the same-size SDS-Aβ42Os of either I32L and F20I32L- or A30L- and V18A30L-Aβ42 variants (Figure 6C-D and E-F, respectively). Interestingly, incorporation of V18L into A30L-Aβ42 leads to an upshift of the ^12^C β-sheet main band, similar to incorporation of V18L into unlabeled Aβ42, which strengthens the above interpretation that V18 is close to β-sheet amide groups in both SDS-Aβ42Os.

In addition to the transmission IR spectra described above, ATR-IR spectra were recorded for SDS- Aβ42Os of F20L-, A30L- and F20A30L-Aβ42 (Figures 7A-B) after drying them from D_2_O solution on the diamond ATR crystal. As observed in the spectra for F20L- SDS-Aβ42Os (blue spectra in Figure 7), a weak band is resolved at 1607.5 and 1606.4 cm^-1^ for small and medium oligomers, respectively, which is not evident in transmission IR spectra for the same samples (Figures 6A-B). This indicates that F20 resides in a β-sheet in the dry state of these oligomers. The absence of the ^13^C-band in solution may suggest a flexible structure of this peptide region which prevents a stable incorporation of the F20 residue into the β-sheet of SDS-Aβ42Os. Another explanation of the ATR-IR results is that drying induces the formation of larger oligomers. However, the ATR-IR spectra for the SDS-Aβ42Os of A30L-, F20A30L- Aβ42 and unlabeled Aβ42 resemble the corresponding solution spectra. In particular, the positions of the β-sheet main bands and of the ^13^C-bands are in the same spectral ranges as in solution, indicating that the extentof their β-sheets and thus their oligomer size are only little affected by drying. In addition, the main band position is still higher for the Aβ42Os prepared in 0.2% SDS, indicating that they are still smaller even in the dry state than those prepared at 0.05% SDS.

**Figure 7.**
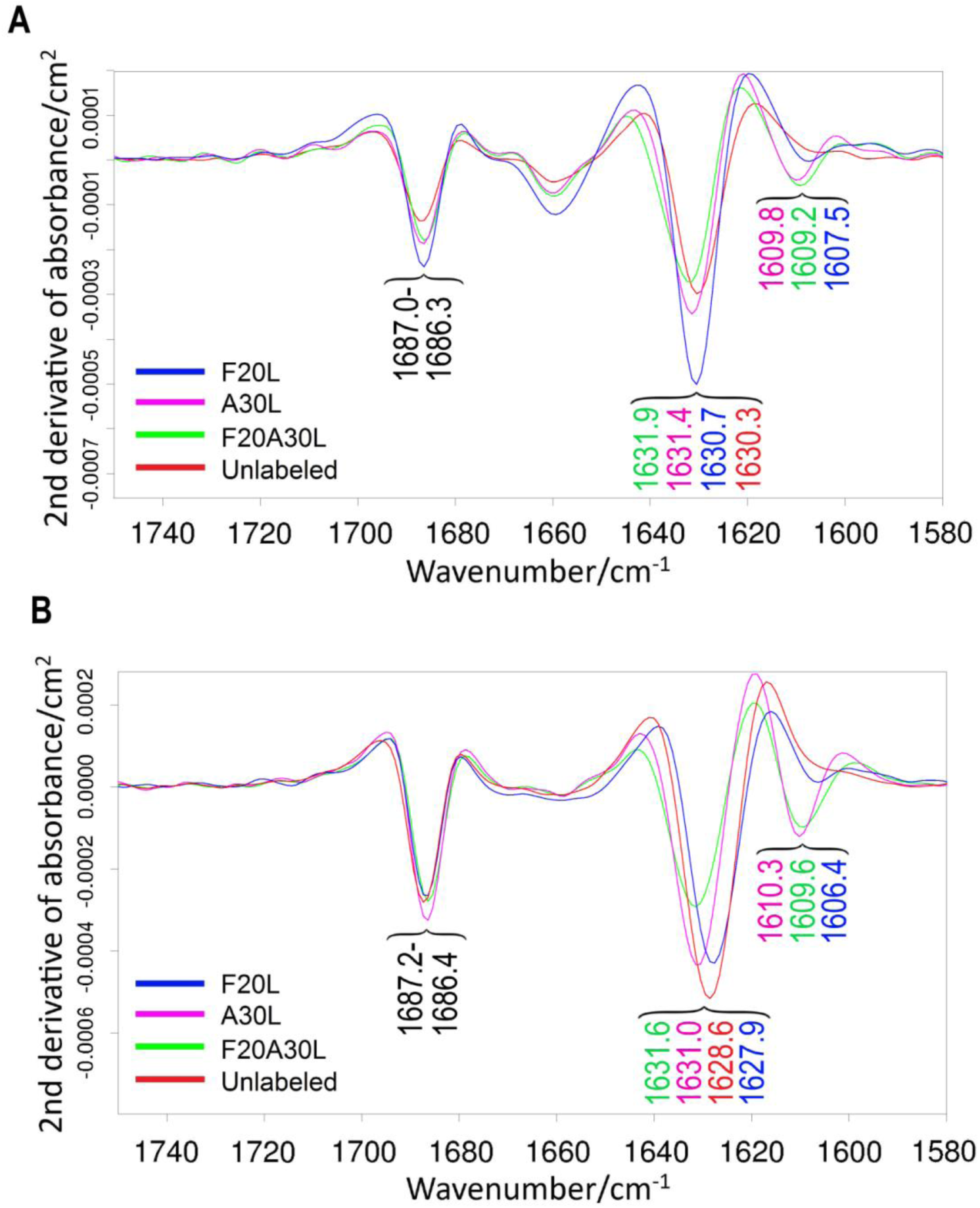
Second derivatives of ATR-IR absorbance spectra (calculated with smoothing range 13) for SDS-stabilized oligomers of the Aβ42 peptide labeled at residues F20 (blue), A30 (purple) and F20A30 (light green), as well as of the unlabeled (red) peptide. Panels A and B show the spectra for the 0.2% SDS-stabilized and 0.05% SDS-stabilized oligomers, respectively. None of the spectra was normalized.

### Comparison of SDS-stabilized and detergent-free Aβ42Os

When the above results are compared to those for the late detergent-free oligomers (Figures 2-3 and Table 1), stark differences are revealed. Most notable is that the ^13^C-band is detected for the detergent-free V18L- and F20L-Aβ_42_Os, but not for their SDS-stabilized oligomers in solution. This indicates that theV18 and F20 residues are not stably inserted in the β-sheet structures of SDS-Aβ42Os, which is in contrast to the case of detergent-free oligomers. On the other hand, a relatively intense ^13^C-band is observed in IR spectra for the SDS-stabilized oligomers of A30L-Aβ42, whereas in corresponding detergent-free oligomers the ^13^C-band is much weaker. One explanation for the stronger ^13^C-band in A30L SDS-Aβ42Os is that there are less amide groups in the A30-containing β-strand in the SDS- Aβ42Os than in the detergent-free Aβ42Os (see Supplementary Information). Moreover, the positions of the ^12^C- and ^13^C-bands of the labeled peptides are significantly upshifted for the SDS-stabilized oligomers as compared to the detergent-free oligomers (Table 1). Taking into consideration such disparities, it can be concluded that the conformation and molecular structures for the detergent-free Aβ42Os and the SDS-stabilized ones are remarkably different.

On the other hand, the early detergent-free Aβ42Os resemble the SDS-Aβ42Os as they do not show a ^13^C- band for F20L-Aβ42Os and only display a very weak band for V18L-Aβ42Os, but instead a relatively large ^13^C-band for A30L-Aβ42Os. This indicates similar structural features in the early detergent-free Aβ42Os and the SDS-Aβ42Os.

### Interconversion of detergent-free and SDS-Aβ42Os

To study the possibility of interconversion between our detergent-free and SDS-Aβ42Os, 0.2% SDS was added to a solution of detergent-free Aβ42Os of F20L-Aβ42. The results are shown in Figure 8 and summarized in Table 4. As evident in Figure 8, within one hour from the moment of SDS introduction into the oligomer solution, the ^12^C-band upshifted significantly (over 2.5 cm^-1^). Initially resolved at 1627.2 cm^-1^ in the absence of detergent, after about two hours from SDS addition, the main band stabilized at about 1630.5 cm^-1^, which is the same value within 0.1 cm^−1^ as for the directly-prepared small SDS-Aβ42Os. The shift from 1627.2 to 1630.5 cm^−1^ indicates reorganization of β-sheets and a decrease in the average size of the oligomers.^63^

**Figure 8.**
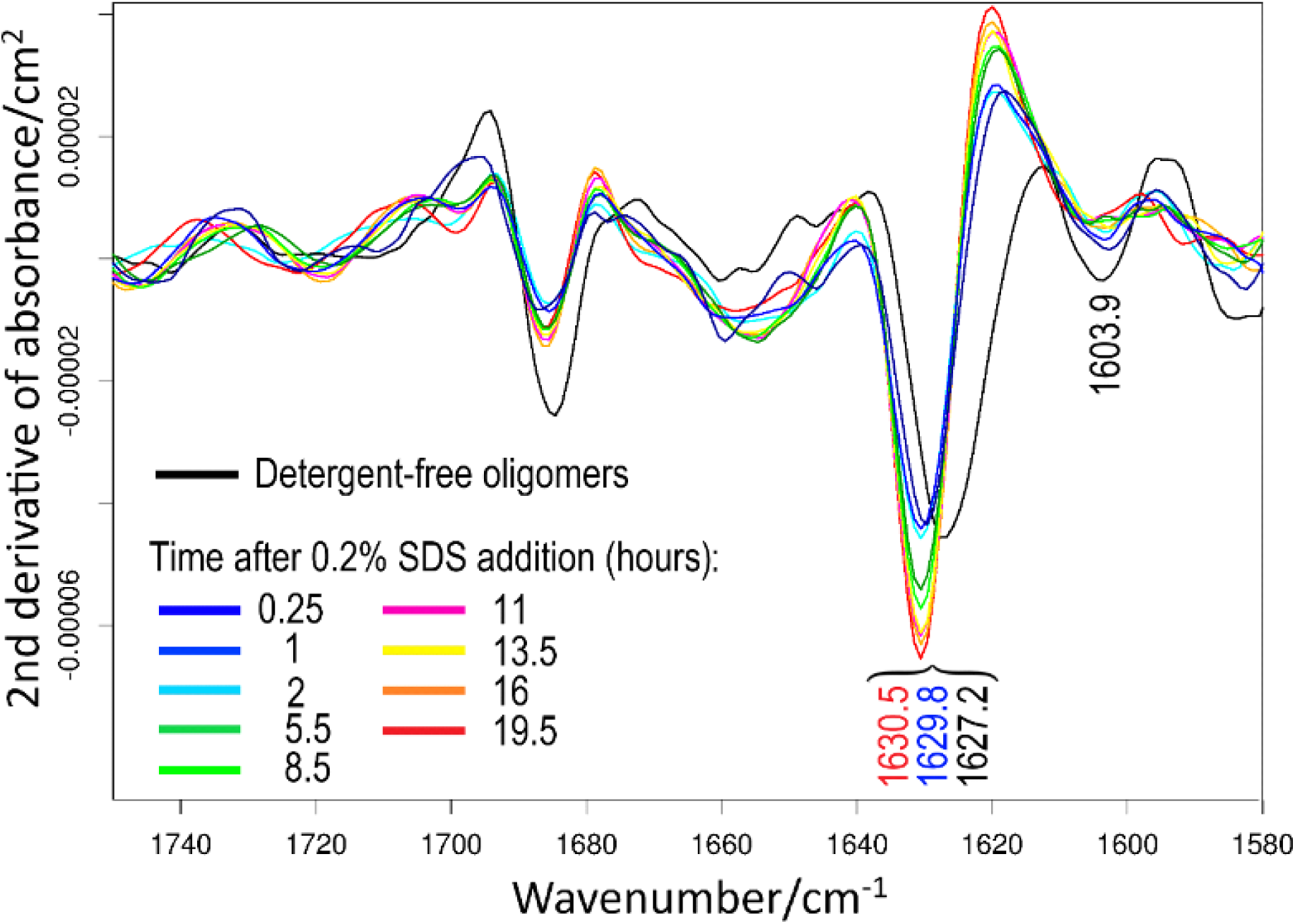
Second derivatives of transmission IR absorbance spectra (calculated with smoothing range 17) for F20L- Aβ42 oligomers before and after addition of 0.2% SDS. The black spectrum indicates the detergent-free oligomers, whereas the spectra for the oligomers at various time-points after SDS addition are shown in different colors.

**Table 4.** The IR band positions and ^12^C-band intensity ratios in the amide I’ region of the second derivative IR spectra for F20L-Aβ42Os before and after addition of 0.2% SDS. The values before and after SDS addition are averages for two independent experiments.

| Time from aggregation induction (h) | $^{13}\text{C}$-band ($\text{cm}^{-1}$) | Main $^{12}\text{C}$-band ($\text{cm}^{-1}$) | $^{12}\text{C}$-band intensity ratio<sup>a</sup> |
| --- | --- | --- | --- |
| 0 | 1603.9 | 1627.2 | 1.8 |
| 0.25 | --- | $1629.8 \pm 0.1$ | $6.3 \pm 1.6$ |
| 1 | --- | $1630.4 \pm 0.1$ | $6.7 \pm 3.5$ |
| 2-20 | --- | $1630.4 \pm 0.2$ | $5.9 \pm 1.7$ |

a) The $^{12}\text{C}$ -band intensity ratio was calculated by dividing the value of the second derivative at the position of the main $^{12}\text{C}$ -band by that of the high-wavenumber $^{12}\text{C}$ -band.

Interestingly, the intensity of the ^13^C-band reduced after SDS introduction and became almost undetectable after a few hours of incubation at 37°C. In addition, the ^12^C-band intensity ratio (main ^12^C- band normalized to high-wavenumber band) increased significantly (Figure 8, Table 4). Such observations indicate changes in F20’s microenvironment and can be interpreted as a reorganization of the F20 residue from within to outside of β-sheets. From the similar spectral properties, we conclude that the final oligomers in these experiments are the intended small SDS-Aβ42Os. The spectral changes monitored here are inversely related to those of the conversion from early to late detergent-free Aβ42Os.

## Discussion

### General architecture of different types of Aβ42Os

The isotope-edited FTIR studies in this work help to further scrutinize the molecular structure of Aβ42Os and to develop relevant molecular models. Our findings shed light on the organization of β-strands in the oligomeric assemblies and on potential intra- or intermolecular contacts between pairs of amino acid residues (V18 and A30, F20 and A30, F20 and I32) in the stretches of hydrophobic amino acids within the central and C-terminal regions of the Aβ42 peptide sequence.

F20L-Aβ42 does not generate a ^13^C-band in IR spectra for our SDS-Aβ42Os and for the early detergent-free oligomers. In contrast, a clear band is observed for the late detergent-free Aβ42Os. When these oligomers disassemble upon addition of SDS, also the F20L ^13^C-band disappears.

V18L-Aβ42 behaves somewhat similar to F20L variant. The band is not detected for SDS-Aβ42Os in solution and is also very small at the beginning of the time-resolved aggregation experiment. However, it significantly grows is intensity and downshifts by several cm^-1^ as late and large detergent-free Aβ42Os form.

In contrast, the ^13^C-band of A30L-Aβ42 is present and relatively strong for small oligomers, either stabilized by SDS or seen in the early phase of the kinetic experiments. Its signal decreases during aggregation relative to the high-wavenumber β-sheet band, while its band position shifts. Both observations indicate a structural change near A30 when the oligomers grow in size.

I32L-Aβ42 produces relatively intense ^13^C-bands in IR spectra for both detergent-free and SDS-stabilized oligomers. The band position is significantly upshifted in case of SDS-Aβ42Os.

These observations demonstrate that the structure of the small and medium SDS-Aβ42Os is remarkably different from that of the late detergent-free Aβ42Os. This is indicated by the different positions of the β-sheet main band, which reflect oligomer size^63^ and by the isotope effects observed upon labeling.

On the other hand, the similar isotope effects for the two types of SDS-Aβ42Os and for the early detergent-free Aβ42Os indicate similar molecular environments of V18, F20 and A30 under all three conditions, indicating that the results obtained in the presence of SDS are also relevant for the detergent-free case. In case of I32, the ^13^C-band is significantly downshifted in larger SDS-Aβ42Os.

The small SDS-stabilized oligomers (∼tetramers) may be the building blocks of the medium SDS-stabilized oligomers (∼dodecamers) because both types show similar isotope effects. This is partially consistent with the suggestion of Yu and colleagues that larger oligomers are formed via assembly of building blocks with the same structure as in smaller oligomers.^31^ In contrast, the isotope effects for large oligomers at the end of the kinetic experiment are different, indicating that the structure of their building blocks is different.

### Involvement of residues V18, F20, A30 and I32 in β-sheets

Our results indicate that all residues V18, F20, A30 and I32 reside in an antiparallel β-sheet in the late detergent-free Aβ42Os formed during kinetic aggregation experiments under near-physiological conditions (solvent polarity, pH, temperature). Our findings are in agreement with several models,^29,31,37,39^ but only in partial agreement with the model for detergent-free Aβ42 pentamers prepared at low temperature.^32^ In this model, the antiparallel β-sheet structure involves V18, F20 and I32, but A30 is just outside the central β-strand. It is noteworthy that the Aβ42Os reported by Ahmed and colleagues are also different from ours in other ways: their secondary structure is considerably disordered, while upon incubation at 37°C for a maximum of 1 hour, fibrillar structures prevail. By contrast, the β-conformation constitutes a major component of our oligomers and they are stable either at room temperature or at 37°C for at least several days.

For our SDS-Aβ42Os and the early detergent-free Aβ42Os, the results indicate that residues A30 and I32 are positioned within a β-sheet structure, whereas residues V18 and F20 are not. These findings are largely inconsistent with predictions from Ahmed’s model,^32^ and are only partly in agreement with the data reported by Yu and colleagues for dimeric constituent units of Aβ42Os stabilized by SDS.^31^ The Yu model predicts the basic structure as a dimer which consists of intramolecular antiparallel β-sheets by residues V17-D23 and K28-I33, whereas an intermolecular parallel in-register β-structure is formed between C-terminal residues 34 to 42 from each monomer. For both of their oligomers, the same study found fast H/D exchange for the amide protons in residues F19 and A21, but slow exchange for those in residues I31 and G33. This indicates unstable hydrogen bonds for the amide groups with their carbonyl in V18 and F20, while stable hydrogen bonds for those with the carbonyl in A30 and I32. This is in good agreement with our SDS-stabilized Aβ42Os spectra in solution where the V18 and F20 amides are not stably incorporated in a β-sheet, while A30 and I32 are. Nevertheless, V18 and F20 have a propensity to be part of a β-sheet, as shown by the effects of V18 labeling on the β-sheet main band and by the appearance of a ^13^C-band for F20L-Aβ42 in the dry state. In the published tetramer structure,^39^ V18 and F20 reside in the outer strands of the β-sheet, which might imply that their strand can easily detach from the core of the sheet.

### Arrangement of β-strands

Our results are in agreement with the antiparallel β-conformation in a putative hairpin structure. This is indicated by the distinct high-wavenumber β-sheet band in the oligomer spectra, observed and assigned previously^30,36^ and also by the isotope effects observed here, which rule out an in-register arrangement of V18, F20, A30 and I32 in a parallel β-sheet. This would generate larger shifts due to intermolecular coupling than observed here. An at least partially antiparallel β-conformation has also been concluded from previous studies some of which proposed a β-hairpin as the structural unit. Also an electron paramagnetic resonance spectroscopy study indicated an antiparallel β-sheet conformation in AβOs.^35^ An intramolecular hairpin-like structure was proposed for artificially-stabilized Aβ monomers,^29^ as well as several molecular AβO models including those for detergent-free oligomers,^32^ oligomers of engineered peptide with intramolecular disulfide bonds,^37,42^ detergent-stabilized small oligomers,^31,39^ detergent-depleted large oligomers,^95^ and oligomers formed within lipid bilayers in computational studies.^33^ Additionally, a previous study from our group concluded that each monomeric peptide unit contributes at least two strands to the β-sheet structure of homooligomers of Aβ40 and Aβ42,^59^ which is consistent with hairpin formation. Hairpin formation has been suggested as a key event in early steps of the peptide aggregation process.^96–98^ Such a structure closely resembles that of protomers for several pore-forming, naturally-occurring peptides and proteins, most notably in α-hemolysin from *Staphylococcus Aureus*,^69^ which has been shown to share structural similarities with AβOs.^70^

### Intra-and intermolecular contacts

Our experiments served also to establish whether or not the labeled residues are close in the three-dimensional structure. When two ^13^C-groups are close in space, their amide I vibrations couple. This results in a spectral change that makes it possible to identify such contacts between ^13^C-groups. The spectral change depends on distance and relative orientation of the coupling amide groups.^88–90^ In antiparallel β-sheets, the strongest downshifts are calculated when the other labeled carbonyl is in an adjacent β-strand and either in a hydrogen bonded amide group (HA) or in an amide group that follows the hydrogen bonded amide in the sequence (HA+1). In these cases downshifts up to -6 cm^-1^ are calculated.^67,88,99^ Further significant shifts are expected for HA-1 interactions. Interestingly, the sign of these shifts depend on whether the carbonyls are oriented toward or away from each other. The HA-1 interaction with a face to face orientation of the carbonyl groups is the only interaction for which an upshift is calculated. The shifts are expected to be smaller for distorted or flexible sheets.

The upshift due to intramolecular coupling between V18 and A30 indicates therefore an HA-1 interaction with the carbonyls pointing into the hairpin. This predicts a HA+1 interaction for F20 and A30 and indeed a downshift due to intramolecular coupling is observed when the carbonyls of these residues are labeled. The downshift is expected for such an interaction. However, it is less than calculated for an ideal, rigid antiparallel β-sheet, possibly because the sheet is distorted close to the loop region. The proposed interactions are the best interpretation for our data and are illustrated in a putative scheme in Figure 9. We also detected intermolecular couplings between V18 amide groups and between F20 and I32 indicating their proximity in the three-dimensional structure.

**Figure 9.**
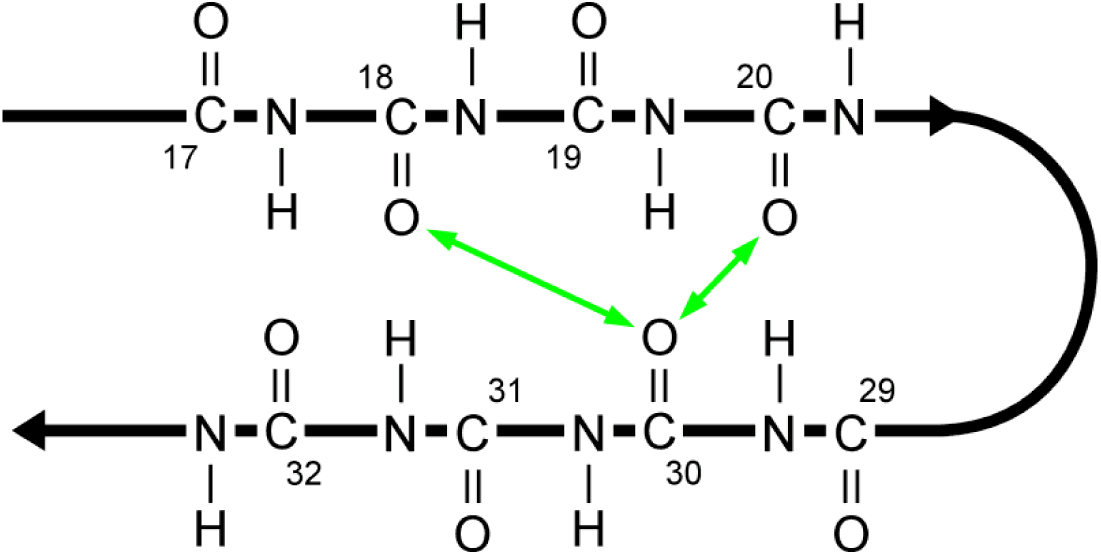
A putative model for the hairpin structure of Aβ42 molecules in detergent-free Aβ_42_Os after 16-18 hours of aggregation at 37°C. The numbers indicate the residue numbers of the carbonyl groups and the arrows the intramolecular couplings detected in the present work.

These results can be discussed in the light of previously suggested molecular AβO models which disagree regarding intramolecular contacts between residues from the central hydrophobic cluster and the C-terminal hydrophobic region. We apply the term *contact* in the following to HA and HA±1 interactions.

V18···A30 intramolecular contacts are not consistent with most published models which position V18 and A30 residues relatively far apart on adjacent strands.^29,31,32,37,39^ This is in contrast to our results, which indicate a HA-1 interaction.

F20···A30 contacts are ruled out in Hoyer’s hairpin model for Aβ40^29^ and in the model by Ahmed and colleagues.^32^ By contrast, the models by Yu *et al*.^31^, Lendel and colleagues^37^ and Ciudad *et al*.^39^ predict or are consistent with a strong intramolecular contact between the amide groups of F20 and A30. In these models, shifts of amino acid registry in the hairpin β-strands as compared to Hoyer’s model bring residues F20 and A30 in adjacent strands closer together and generate strong F20···A30 contacts. More specifically, in the models of Yu *et al*. and Ciudad *et al.*, the carbonyl groups of F20 and A30 belong to amide groups that are hydrogen bonded to each other via the NH group of A21 (HA interaction). Such an interaction is difficult to reconcile with our data for two reasons: (i) it is expected to generate a strong down-shift for F20A30L-Aβ42O due to intramolecular coupling, but only a small downshift of 1.3 cm^-1^ is observed and (ii) it is incompatible with the upshift due to intramolecular coupling observed for V18A30L-Aβ42O as Yu *et al.* predict this to be a HA-2 interaction, which is calculated to have a negligible influence on the ^13^C-band position.

Regarding intramolecular F20···I32 contacts, Hoyer’s hairpin model predicts a strong HA interaction between the carbonyls of these residues.^29^ Other models are also consistent with intramolecular F20···I32 contacts.^31,32,37^ Whereas we cannot definitely rule out this type of contact in our detergent-free Aβ42Os, isotope dilution studies point to intermolecular interactions as the major source of contact between F20 and I32 residues.

Our SDS-Aβ42Os were prepared according to a protocol^63,65^ similar to the one used by Yu and colleagues.^31^ Nevertheless, no contact was detected for our SDS-Aβ42Os, whereas a strong F20···A30 contact would be expected from Yu’s dimer model. An important reason for this discrepancy seems to be that our SDS-Aβ42Os in solution do not include F20 in their β-sheets. We conclude that none of our SDS-Aβ42Os adopts the hairpin structure with the registry and arrangement proposed by Yu and colleagues.^31^ The discrepancy between our results and those predicted by the model may be due to the nature of the Aβ42 peptide used and also the oligomerization conditions in each study. Yu and colleagues used recombinantly-expressed Aβ42 with an additional Met residue at the N-terminal, whereas in our study Aβ42 from a synthetic source was utilized for preparation of the oligomers. Moreover, the oligomer samples in Yu *et al* study were concentrated before the biophysical measurements unlike ours, whereas we prepared our oligomers in a D_2_O-based buffer, in contrast to PBS in H_2_O which was the buffer used by Yu and colleagues. D_2_O can influence the stability of proteins^100^ and is reported to affect the kinetics of Aβ aggregation process.^101^ Despite that, our oligomers prepared in D_2_O have the same size^63^ as the ones in the original report,^65^ which were prepared in H_2_O-based buffer and our data with singly-labeled peptides are in line with the H/D exchange data by Yu *et al*..

We turn now from intramolecular to possible intermolecular contacts, which are indicated in our work for V18-V18 and for F20-I32 in the late detergent-free Aβ42Os. None of these are suggested in models for dimers,^31^ pentamers,^32^ hexamers,^37^ and tetramers/octamers.^39^ An explanation is that our late detergent-free Aβ42Os are larger than the Aβ42Os in the cited studies and that the here-described conformational changes between early and late detergent-free Aβ42Os generate these contacts.

## Conclusions

This work studied AβOs with isotope-edited FTIR spectroscopy. Advantages of the chosen approach were that oligomers in a size range from 20 to 100 kDa could be studied, which is difficult to cover by NMR and cryogenic electron microscopy, and that time-resolved experiments in aqueous solution were possible. According to our knowledge, this is the first time that residue-specific structural information from the same method is compared for different types of AβOs.

The following scenario is concluded from our results. Small and medium oligomers, either formed in the early phase of kinetic experiments or stabilized by SDS, incorporate A30 in β-sheets. V18 and F20 are not part of a stable β-sheet but may reside in a flexible β-strand that is loosely attached to the β-sheet core. When the oligomers grow from ∼60 kDa to ∼100 kDa, a conformational change within the oligomer building blocks incorporates V18 and F20 into the β-sheets and may lengthen the β-strands. Vibrational coupling between V18 and A30 as well as between F20 and A30 within the monomeric building block can then be detected. When these oligomers shrink in size upon addition of SDS, also F20 detaches from the β-sheet. Accordingly, the structures of the building blocks is different for small and medium Aβ42Os on the one hand and large Aβ42Os on the other hand. In contrast, small SDS-stabilized Aβ42Os (∼tetramers) and medium SDS-stabilized Aβ42Os (∼dodecamers) show similar isotope effects and therefore the small Aβ42Os may be the building blocks of the medium Aβ42Os.

The findings from this study add to the available information on the internal structure of Aβ oligomers and help to develop a better understanding of structural features for the peptide’s most neurotoxic aggregates.

## Acknowledgements

We are indebted to Eeva-Liisa Karjalainen and Cesare M. Baronio for our program for calculating the amide I spectrum. We also thank Oleksandra Kurysheva for her assistance with experimental work. This work was supported by the Swedish Research Council (2021-04595), Hjärnfonden (FO2019-0127), Olle Engkvists stiftelse (211-0051), and Magn. Bergvalls stiftelse (2020-03624).

