## Supplementary Information for "Residue-level structural information from isotope-edited FTIR spectroscopy reveals distinct molecular structures of different amyloid-β42 oligomers"

### ISOTOPE EFFECTS IN SIMULATED AMIDE I SPECTRA

**Introduction** – We performed simulations of the amide I absorption of β-sheets as described in the Methods section of the main text. Our simulations used parameters that were optimized by comparison to density functional theory simulations.[1,2] The only adjustable parameter was the local wavenumber of an unperturbed amide group, which was chosen to be 1707 cm$^{-1}$ because it corresponds to the wavenumber of *trans*-*N*-methylacetamide in a $N_2$ matrix (1706-1707 cm$^{-1}$)[3,4] and is close to those in $H_2$ (1710 cm$^{-1}$)[5] and Ar (1708 cm$^{-1}$)[4] matrices. In spite of this limited possibility to adjust the parameters to the experimental results, the main absorption band of the unlabeled peptide is calculated close to our experimental results: 1625.5-1621.3 cm$^{-1}$ for antiparallel β-sheets of 8 or 24 strands and 6 or 10 residues. Relevant spectral properties are compiled in Tables S1 to S14 in Appendix II.

**Terminology** – As in the main text, we will use the term $^{13}$C-band for the strong low wavenumber normal modes that are dominated by the amide I vibration of the labeled amide groups although $^{12}$C-groups contribute as well to these modes. Also the term "main $^{12}$C-band" for the main absorption band in the spectrum is a simplification because in rare cases $^{13}$C-groups may contribute substantially to the strongly absorbing normal modes in this spectral region. Similarly we use "$^{12}$C-modes" and "$^{13}$C-modes" for normal modes with a strong contribution of the respective isotope, while noting that also the other isotope may contribute considerably. Strong and weak modes are normal modes that give rise to strong and weak absorption of infrared light, respectively. Amide groups are numbered according to the residue number of their carbonyl group. For example, amide group 5 is the amide group between residue 5 (containing the carbonyl group of amide group 5) and residue 6 (containing its amino group).

**Antiparallel β-sheet models with labels in every second strand** – The isotope effects for a single strand of an antiparallel β-sheet (10 residues, 9 complete amide groups) are compiled in Table S1. A clear $^{13}$C-band is seen for all label positions (groups 1-8) except for the C-terminal amide group (group 9), which causes only a shoulder on the low wavenumber side of the main $^{12}$C-band. This end-effect has also been observed in previous calculations.[6] The deviating behavior of group 9 can be explained by its higher local wavenumber, which is also seen in density functional theory calculations of a single strand.[2] This increases the wavenumber of the $^{13}$C-modes and separates them less from the main $^{12}$C-band wavenumber than required for generating a distinct $^{13}$C-band.

The main $^{12}$C-band is upshifted by 1 to 2 cm$^{-1}$ for central label positions (amide groups 3-7), but not or less for those at the strand ends (groups 1, 2, 8, and 9). The former can be explained by the large participation of the central amide groups

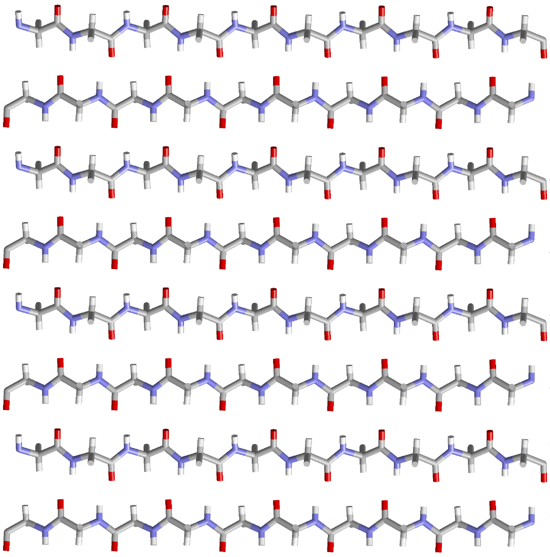


**Figure S1.** Antiparallel β-sheet with 8 strands and 10 residues per strand (9 amide groups). The residue numbers are indicated. Atoms of incomplete amide groups at the strand termini are ignored in the calculations.

in the strongest amide I mode of the unlabeled peptide. Labeling one of the central amide groups abolishes its contribution to the strongest $^{12}$C-modes and largely localizes the strongest $^{12}$C-mode on the preceding and/or the following amide groups. This strong impact of labeling on the nature of the strongest $^{12}$C normal mode explains the different $^{12}$C-band positions of unlabeled and centrally labeled strands.

The intensity of the $^{13}$C-band is larger than expected from the $^{13}$C-group abundance in the structure and close to 25% of the main $^{12}$C-band intensity in the central part of the strand (groups 2-8). The enhancement is due to the out-of-phase contributions of the unlabeled nearest sequence neighbors to the $^{13}$C-normal mode.

We then studied β-sheets consisting of 8 or 24 strands with 6 or 10 residues (5 or 9 complete amide groups) each. The results are compiled in Tables S2 to S7. The in-register 8-stranded sheet is shown in Figure S1 and is identical to the first 8 strands of the 24-stranded β-sheet. These sheets model a tetramer and a dodecamer of amyloid-β (Aβ) hairpins where each Aβ molecule contributes two adjacent strands to the sheet. We considered a single, labeled amide group per Aβ hairpin, which gives one labeled amide group in every second strand of the sheet. The results for the 8- and 24-stranded sheets with 10 residues (Table S3 and S6) are similar to those obtained for the single strand (Table S1). Only labeling of the terminal amide groups does not produce a clear $^{13}$C-band as noted before.[6] For all label positions that are not at the strand ends, a clear $^{13}$C-band is calculated and the intensity of the $^{12}$C-band reduced

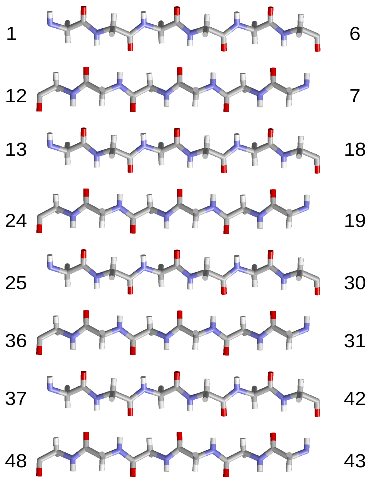


**Figure S2.** Antiparallel β-sheet with 8 strands and 6 residues per strand (5 amide groups). The residue numbers are indicated.

while that of the $^{13}$C-band is enlarged relative to the $^{13}$C-group abundance.

For label positions in the central portions of the strands, the main $^{12}$C-band is upshifted. It is interesting, that the $^{12}$C-band band shift does not seem to be correlated with the intensity reduction of the same band. For example, the intensity reduction is considerable even for label positions close to the strand termini (amide groups 2 and 8) for which the $^{12}$C-band shift is very small.

The high wavenumber band of β-sheets shifts down upon labeling as observed previously in experiments.[7] This is particularly pronounced for the sheet with 6 residues per strand (Table S2). Its intensity however, is little affected by labeling: on average it is slightly reduced to 98% with a standard deviation of 2.6 percentage points. The lowest value was 93% and the highest 103% (Tables S2, S3, and S5).

The only isotope effect that has a strong dependency on the number of strands is the difference between the $^{13}$C-band wavenumber $\tilde{\nu}(^{13}\mathrm{C})$ and the main $^{12}$C-band wavenumber $\tilde{\nu}(^{12}\mathrm{C_{UL}})$ of the unlabeled model – $\Delta\tilde{\nu}(^{13}\mathrm{C}-^{12}\mathrm{C_{UL}}) = \tilde{\nu}(^{13}\mathrm{C}) - \tilde{\nu}(^{12}\mathrm{C_{UL}})$ – which reduces with increasing number of strands. This is due to two effects:

(i) Vibrational coupling between amide I vibrations in neighboring strands leads to a stronger downshift of the main $^{12}$C-band with more strands.

(ii) Vibrational coupling of the $^{13}$C-amide groups with other $^{13}$C- or with $^{12}$C-groups generates a downshift of the $^{13}$C-band that is much smaller than the downshift of the $^{12}$C-band.

Because of these effects, the spectral positions of the main $^{12}$C-band and of the $^{13}$C-band are closer for sheets with more strands. This is exemplified in more detail in Appendix I.

Also the absolute positions of the $^{13}$C-band are different for different sheets. They decrease for labels in the core of the strands (amide groups 3-7) from ~1635 cm$^{-1}$ for the single strand to 1600 cm$^{-1}$ for the 8-stranded sheet and to 1598 cm$^{-1}$ for the 24-stranded sheet due to lower local wavenumbers in the sheets compared to the single strand and stronger coupling induced downshifts when the number of strands is increased. The latter explains the small difference between the 8-stranded and the 24-stranded sheet because the averaged local wavenumbers of amide groups 3-7 in each strand were identical for both sheets.

For other spectral parameters, there is very little difference between the single strand and the 8-strand results regarding effects of isotopes in the central portion of the strands (corresponding to residues 3 to 7). This holds for the intensity of the $^{13}$C-band relative to the main $^{12}$C-band in the spectrum of the labeled sheet, $I(^{13}\mathrm{C})\,/\,I(^{12}\mathrm{C})$, for the intensity of the main $^{12}$C-band of the labeled structure relative to that of the unlabeled model, $I(^{12}\mathrm{C_L})\,/\,I(^{12}\mathrm{C_{UL}})$, as well as for the shift of the main $^{12}$C-band due to labeling $\Delta\tilde{\nu}(^{12}\mathrm{C_L}-^{12}\mathrm{C_{UL}}) = \tilde{\nu}(^{12}\mathrm{C_L}) - \tilde{\nu}(^{12}\mathrm{C_{UL}})$. The effects are slightly stronger for the 24-stranded sheet but because of the considerable variation of the effects of different label positions, this influence is likely of little diagnostic value. While the $^{12}$C-band shifts up for centrally placed labels it shifts slightly down for labels at the edge of a sheet. This effect is seen clearest for the 8-stranded sheet.

In contrast to the number of strands, a smaller number of residues per strand strengthens the isotope effects on these spectral properties dramatically. Table S2 lists results obtained for an 8-stranded sheet with 6 residues (5 amide groups) and Figure S2 shows its structure. When the number of amide groups per strand is reduced from 9 to 5 in the 8-stranded β-sheet, the intensity of the $^{13}$C-band relative to the main $^{12}$C-band $I(^{13}\mathrm{C})\,/\,I(^{12}\mathrm{C})$ is strongly enhanced (on average from 23% to 58%, see Tables S2 and S3) and the shift of the main $^{12}$C-band upon labeling the central position (residue 3 in every second strand) is nearly three times as large (~9 cm$^{-1}$) as the largest shift noted for the sheets with 9 amide groups (~3 cm$^{-1}$). It is notable that the $^{13}$C-band can be nearly as strong (~70%) as the $^{12}$C-band in the narrower sheet, much more than expected from the abundance of $^{13}$C-amide groups.

The stronger isotope effects in the narrower sheet with 5 amide groups are conceivable, because the labeled groups are relatively more abundant here than in the wider sheet with 9 amide groups. Ignoring the terminal amide groups, the abundance of labeled groups $a(^{13}\mathrm{C})$ in the narrow sheet is 17% (groups 2-4 in each strand, 4 labeled groups out of a total of 24 amide groups) and in the wider sheet 7% (groups 2-8, 4 labeled groups out of 56 groups). The $^{13}$C-band of the narrow sheet is enhanced relative to the $^{13}$C-group abundance by a factor of 3.4 ($I(^{13}\mathrm{C})\ I^{-1}(^{12}\mathrm{C})\ a^{-1}(^{13}\mathrm{C}) = 58/17$) and by the similar factor of 3.3 (= 23/7) for the wider sheet. When the number of strands of the wider sheet is increased to 24 strands, the relative abundance of labeled groups remains the same, but the average relative $^{13}$C-band intensity becomes somewhat higher (29%) giving a relative intensity enhancement of 4.1. The similar average values for different sheets indicate that the relative abundance of the labeled groups is a main factor for the relative $^{13}$C-band intensity $I(^{13}\mathrm{C})\,/\,I(^{12}\mathrm{C})$.

In the narrow sheet, $I(^{13}\mathrm{C})/I(^{12}\mathrm{C})$ varies strongly for different label positions (50% - 71%) whereas it is more uniform for the wider sheet. The variation is largely due to a variation of $I(^{12}\mathrm{C})$, whereas $I(^{13}\mathrm{C})$ is relatively constant.

The effect of sheet width was reproduced in a sheet of 4 amide groups, which was generated from an out of register arrangement of 8 strands with 10 residues as shown in Figure S3. The last 4 amide groups in each strand formed an antiparallel β-sheet with complete sheet-type hydrogen bonding. This sheet produced isotope effects that were very similar to those of the 8-stranded sheet with 5 amide groups but the maximal shifts of the $^{13}$C- and $^{12}$C-band relative to the $^{12}$C-band of the unlabeled sheet were slightly larger as compiled in Table S4.

The purpose of this model was to test whether non-hydrogen bonded groups with a local backbone conformation as in a β-sheet would also generate a distinct $^{13}$C-band. Accordingly, the first amide groups extended from the sheet and were not hydrogen bonded. Groups 1-3 are prime examples for illustrating the effects of lacking hydrogen bonds while keeping the same backbone conformation. This results in local wavenumbers that are 30 cm$^{-1}$ higher than those of amide

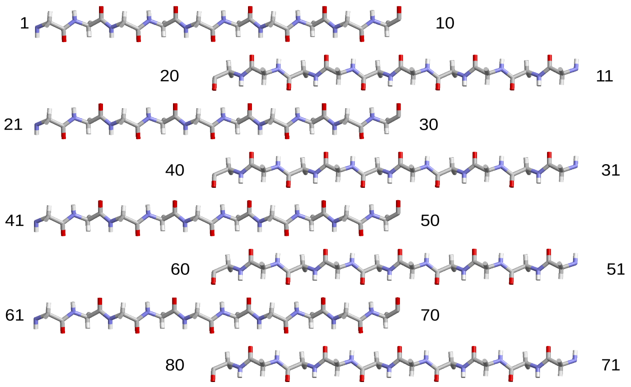


**Figure S3.** Out of register arrangement of 8 strands with 10 residues of which the C-terminal four amide groups form an antiparallel β-sheet. The residue numbers are indicated.

groups 6-8 in the sheet. The consequence of this are weak $^{13}$C-modes with wavenumbers that are higher than the main $^{12}$C-band wavenumber. Accordingly, no low wavenumber $^{13}$C-band is seen in the spectrum for labels close to the N-termini of the strands. Because the $^{13}$C-modes were on the high wavenumber side of the main $^{12}$C-band, the main $^{12}$C-band was slightly upshifted.

Groups 4 and 5 in each strand seemed to experience the electric field of the adjacent β-sheet because their local wavenumbers were higher or lower, respectively, than those of amide groups 1-3. The local wavenumber of the middle amide groups was intermediate between those of the N-terminal groups and those in the sheet. Upon labeling these groups in every second strand (at 5, 25, 45, 65 or 15, 35, 55, 75), most $^{13}$C-modes were on the low wavenumber side of the main $^{12}$C-band, but they were weak and therefore not resolved.

The C-terminal amide group in each strand showed unusual isotope effects. This group was part of the β-sheet in all strands. When it was labeled in every second strand (at position 9, 29, 49, 69 or 19, 39, 59, 79), the main $^{12}$C-band shifted down by ~6 cm$^{-1}$. This is due to a strong, but unresolved $^{13}$C-mode on the low wavenumber side of the strongest $^{12}$C-mode. Because the local wavenumbers of the terminal amide groups are higher than those of amide groups within the sheet section, the isotope shift is not sufficient to produce a well-separated $^{13}$C-band when the C-terminal amide groups are labeled.

**Antiparallel β-sheet models with labels in every third strand –** Some of the models for Aβ42 oligomers assume a 3-stranded β-sheet section for individual Aβ molecules[8,9] and such a sheet is in line with our previous infrared study.[10] Therefore we tested the effects of labeling a single amide group in every third strand using an antiparallel β-sheet with 24 strands and 10 residues (Table S7). Compared to the same sheet with an isotope label in every second strand, the isotope effects on the $^{13}$C- and $^{12}$C-band intensities were similar in spite of the fewer labels. The isotope shift of the $^{13}$C-band, $\Delta\tilde{\nu}(^{13}\text{C}-^{12}\text{C}_{\text{UL}})$, was smaller and the upshift of the $^{12}$C-band, $\Delta\tilde{\nu}(^{12}\text{C}-^{12}\text{C}_{\text{UL}})$, larger. An interesting difference was that the observation of a distinct $^{13}$C-band was more restricted to labels in the core of the β-sheet: when every third strand was labeled, only labels in positions 3-7 generated a distinct $^{13}$C-band, in contrast to positions 2-8 when every second strand was labeled. Instead, label positions 2 and 8 generated a clear shoulder when every third strand is labeled. The reason that this shoulder does not show up as a distinct band is a combination of lower intensity of the $^{13}$C-modes, higher intensity of the strongest $^{12}$C-mode and slightly less

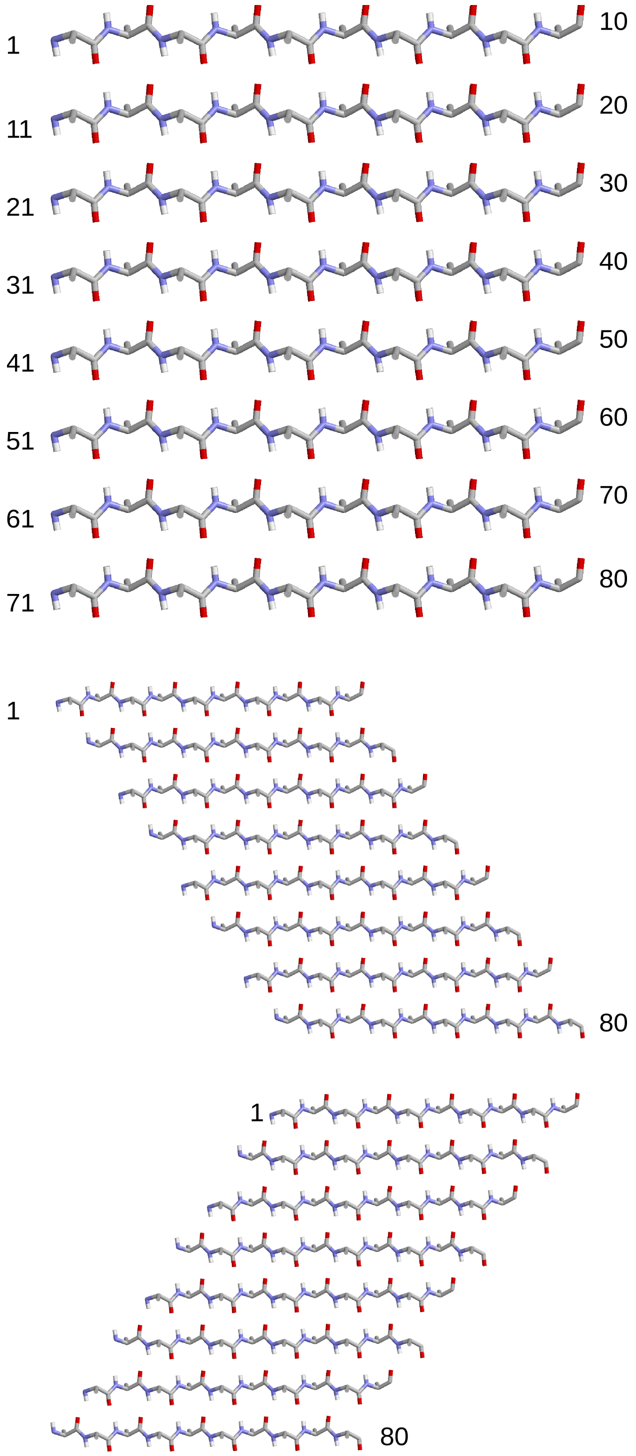


**Figure S4.** Parallel β-sheet with 8 strands and 10 residues per strand (9 amide groups). The residue numbers are indicated. In register arrangement (top) and two out-of-register arrangements where subsequent strands are shifted to the C-terminus (middle) or the N-terminus (bottom) of the preceding strand.

wavenumber separation between the strongest $^{13}$C- and $^{12}$C-modes.

**Parallel β-sheet models** – Parallel β-sheet sections have been suggested for Aβ oligomers.[11] Thus, we explored also the effects of labeling a parallel β-sheet of 8 strands and 10 residues. In-register and out-of-register versions of this sheet are shown in Figure S4. The following discussion compares the in-register parallel β-sheet results with those obtained with the 8-stranded antiparallel β-sheet, where every second strand contained a label based on a hairpin structure of individual Aβ molecules. In both cases, the upshift of the $^{12}$C-band and the intensities of the $^{13}$C- and $^{12}$C-bands were similar (compare Tables S3 and S8) but the downshift of the $^{13}$C-band was much larger for the parallel β-sheet. This is due to the strong vibrational coupling between the labeled groups because they reside in-register in every strand. When this coupling is reduced by placing a label only in every second strand, the $^{13}$C-band downshifts became similar to those of the antiparallel β-sheet (see Tables S3 and S9). The other isotope effects did not change significantly due to isotope dilution.

Different results were obtained when each strand in the parallel sheet was labeled but when the labeled groups were out-of-register, either by shifting their positions within the sheet (*e.g.* labels at residues 1,12,23,34,45,56,67,78, results in Table S10), or by offsetting adjacent strands as shown in Figure S4 and compiled in Tables 11 and 12. Out-of-register labeling considerably increased the $^{12}$C-band shifts $\Delta\tilde{\nu}(^{12}\mathrm{C_L}-^{12}\mathrm{C_{UL}})$, the intensity of the $^{13}$C-band relative to that of the $^{12}$C-band $I(^{13}\mathrm{C})\,/\,I(^{12}\mathrm{C})$, and decreased the relative intensity of the $^{12}$C-band, $I(^{12}\mathrm{C_L})\,/\,I(^{12}\mathrm{C_{UL}})$.

**Vibrational coupling and isotope dilution effects** – As mentioned in the previous section, the $^{13}$C-band of our parallel β-sheet model had a higher wavenumber when only every second strand contained one label. This upshift was ~9 cm$^{-1}$ and corresponds approximately to the effect of a 1:1 dilution of labeled with unlabeled peptides. In this diluted stage, the isotope effects for the parallel β-sheet were similar to those of the corresponding antiparallel β-sheet, which also had a label in every second strand (Tables S3 and S9).

Further dilution was studied for a 9-stranded β-sheet with 6 residues (see Table S5), the structure of which corresponded to that shown in Figure S2 with an extra strand at the C-terminus. To facilitate the interpretation, we labeled only groups in the inner strands, *i.e.* those that were hydrogen bonded to both the amide oxygen and hydrogen atom, so that the local wavenumbers differed by less than 4 cm$^{-1}$. With a label in every second strand, the normal mode wavenumber was 8.2 cm$^{-1}$ lower than the local wavenumber of the $^{13}$C-amide groups. This downshift is due to vibrational coupling of the $^{13}$C-group with other $^{12}$C- and $^{13}$C-groups.

When the labeled peptides are strongly diluted with unlabeled peptides in an isotope-dilution experiment, there will be only a single labeled group in an oligomer. To model this case, we labeled one of the three central amide groups in strand 4, which generated a downshift of the normal mode wavenumber of 4.9 cm$^{-1}$ from the local wavenumber of the $^{13}$C-amide group. Thus coupling of a single $^{13}$C-group with only $^{12}$C-groups accounts for a ~5 cm$^{-1}$ downshift. This downshift is ~3 cm$^{-1}$ less than when a group in every second strand is labeled. Thus, isotope dilution from a label in every second strand to a single label per sheet will upshift the $^{13}$C-band by ~3 cm$^{-1}$.

In all multi-label cases considered in this section, the labeled amide groups were part of a hydrogen bonded chain of amide groups, typical for an in-register arrangement. For such an arrangement, the discussed results indicate an upshift of the $^{13}$C-band by close to 10 cm$^{-1}$ upon 1:1 isotope dilution from one label in every strand to one label in every second strand, and a further ~3 cm$^{-1}$ shift upon further dilution. The band shifts caused by isotope-dilution arise because coupling between amide groups with the same isotope is more effective.

**Isotope effects in the α-hemolysin β-barrel** – The β-barrel structure of α-hemolysin (pdb code 7AHL)[12] shares structural

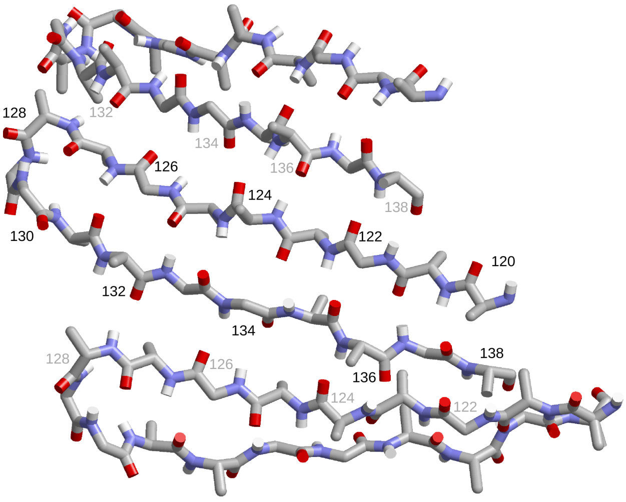


**Figure S5.** Three β-hairpins of the α-hemolysin β-barrel (pdb code 7AHL). Residue numbers of the middle hairpin are indicated in black and of the other hairpins in gray.

similarities with Aβ oligomers.[13] Therefore, we explored isotope effects for such a structure. It consists of seven β-hairpins that form a distorted antiparallel β-sheet of 14 strands. Figure S5 shows three of these hairpins and Table S13 compiles the results. The local wavenumbers vary along the hairpin and between the same amide groups in different hairpins. In general, amide groups in the N-terminal section (amide groups containing C=O of residues 121-124, and 126) have low local wavenumbers (~1660 cm$^{-1}$), indicative of strong, favorable electrostatic interactions like hydrogen bonding, followed by groups in the C-terminal section (residues 133-136) with somewhat higher local wavenumbers (~1670 cm$^{-1}$), and groups in the loop region (residues 127-131) as well as at the termini (residues 120 and 137) with the highest local wavenumbers (~1691 cm$^{-1}$) because of weak electrostatic interactions. Of these, the amide groups of residues 127 and 129 experience intermediate interactions in some hairpins and weak interactions in others, leading to a 40 cm$^{-1}$ spread in their local wavenumbers. Two amide groups do not follow the general pattern: amide group 125, which has a considerably higher local wavenumber (~1677 cm$^{-1}$) than other groups in the N-terminal section, and group 132, which in most hairpins experiences strong to very strong interactions but only intermediate interactions in one hairpin. This results in a 37 cm$^{-1}$ variation of its local wavenumbers.

The local wavenumber variations are clearly reflected in the isotope effects. Distinct $^{13}$C-bands are observed when amide groups are labeled that have low local wavenumbers already when they are unlabeled (residues 121-124, 126, 132, 134 and 135). The $^{13}$C-band wavenumber correlates with the local wavenumber of the group before labeling, being lower for lower local wavenumbers. Thus the electrostatic interactions of an individual amide group – reflected by its local $^{12}$C-wavenumber – is an important influence on the $^{13}$C-band position when this group is labeled.

Many other label positions that do not produce a distinct $^{13}$C-band show a clear shoulder (amide groups 125, 127, 133, 136, 137) or increase the intensity (groups 120, 129, and 131) on the low wavenumber side of the main $^{12}$C-band. The exceptions are amide groups 128 and 130 in the loop, for which there is no or very little intensity increase. Their local $^{13}$C-wavenumbers are high due to weak electrostatic interactions in all hairpins so that the $^{13}$C-normal modes have a higher wavenumber than the main $^{12}$C-band.

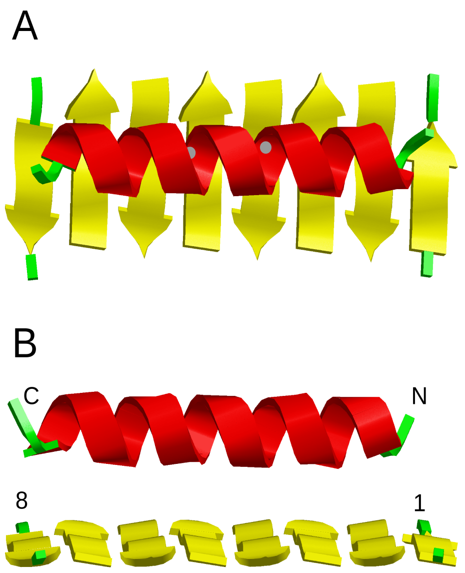


**Figure S6.** Top (A) and side (B) view of the helix-sheet model. The positions of the carbonyl carbon atom of helix residues 9 and 13 are indicated by gray circles in panel A. N- and C-terminus of the helix and strands 1 and 8 of the sheet are indicated in panel B.

Interestingly, the intensity of the $^{13}$C-band seems to increase with decreasing local wavenumber variation of the labeled groups. Accordingly, amide group 135 has the highest band intensity, followed by groups 121-124, 126 and 134 with medium intensity, and by group 132 with low intensity.

The average isotope effects are within the range observed for the antiparallel β-sheets of 8 and 24 strands with 6 or 10 residues. For most amide groups of α-hemolysin, a large isotope shift $\Delta\tilde{\nu}(^{13}C{-}^{12}C_{UL})$ correlates with a weak $^{13}$C-band and a strong $^{12}$C-band. This can be explained by less coupling between $^{13}$C- and $^{12}$C-vibrations when their wavenumbers are further apart. The same effect can be seen when the parallel β-sheet results are compared. In-register labeling leads to a strong downshift $\Delta\tilde{\nu}(^{13}C{-}^{12}C_{UL})$ and small intensity effects, whereas out-of-register labeling causes the opposite. However, this is not the only effect, since there is no clear correlation when all data are analyzed. A further effect on the band intensities is for example the relative abundance of labeled amide groups as discussed above.

**A helix-sheet model with single isotope labels** – The calculations of the out-of-register sheet shown in Fig. S3 established that a $^{13}$C-band below 1620 cm$^{-1}$ is only observed when the labeled group is hydrogen bonded. Additionally, the α-hemolysin calculations showed that even weak hydrogen bonding as in the loops of the β-hairpins is not sufficient to produce such a band. Both observations indicate that labeled residues outside β-sheets will not produce a distinct $^{13}$C-band. We wanted to challenge this conclusion by studying the effects of labeling a hydrogen bonded group outside but close to a β-sheet. As a model we chose an α-helix on top of the 8-stranded antiparallel β-sheet with 6 residues as shown in Fig. S6. The number of amide groups in the helix was 21. The orientation of the helix was chosen such that the transition dipole moments of the amide groups in both substructures were largely parallel or antiparallel and thus had optimum conditions to couple. Nevertheless, the large majority of the normal modes of the unlabeled model where localized on either the sheet or the helix. In 40 of the 61 normal modes, the contribution of one of the substructures to the vibrational energy was less than 10% with an average of 3.3%. Only to 6 normal modes both substructures contributed between 20% and 80%. Nevertheless, the spectrum of the complete model is different from the added helix and sheet spectrum, indicating long-range vibrational coupling in line with previous observations that helix packing[14] and β-sheet stacking[15–19] influences the amide I spectrum.

As intended, the local wavenumbers of the central helix residues were in the same range as those in the central part of the β-sheet: amide groups 1 and 3 in each strand had a lower wavenumber, amide groups 2 and 4 a similar wavenumber (within 1 cm$^{-1}$), and amide group 5 a higher wavenumber than the central helix amide groups.

We placed a single label in the central helix at residue numbers 9 or 13, because the respective amide groups were hydrogen bonded at both the oxygen and the hydrogen atom, and were close to the β-sheet (~7.0 Å distance between closest C=O groups). For comparison, single labels were also placed in the β-sheet at residue numbers 19 to 23 in strand 4 and at position 27 in strand 5. In all cases, a normal mode – highly localized on the labeled amide group – was obtained with a lower wavenumber than the main $^{12}$C-band of the β-sheet. However, a distinct $^{13}$C-band is only observed for labels in one of the central three amide groups in a strand (Table S14). The second derivative spectra of the labeled helices are hardly distinguishable from the unlabeled spectra in the low wavenumber range, whereas all labeled β-sheets produce clear changes, even those without a distinct $^{13}$C-band in the absorption spectrum.

These differences between helix and sheet labels are caused by the low intensity of the $^{13}$C-mode when the helix is labeled, which is similar to that of an isolated amide I oscillator[20] but five times less than when the central amide groups in β-strands are labeled. The $^{13}$C-helix modes are more localized on the labeled amide group than most $^{13}$C-sheet modes and this lower participation of $^{12}$C-groups in the $^{13}$C-mode correlates with its lower intensity. The exception is the $^{13}$C-mode of labeled sheet residue 19, which is slightly more localized on the labeled group (energy contribution 96%) than when the helix is labeled (95%). Nevertheless it has more than twice the intensity. This points to a role of the specific coupling with $^{12}$C-groups in β-sheets for the high intensity of the $^{13}$C-band. The interstrand coupling seems to be particularly decisive for the high intensity, as single labels in a single strand produce a $^{13}$C-mode with similar intensity as single labels in a helix.

Furthermore, helix labeling does not change the position and intensity of the main β-sheet band in contrast to central strand labeling, which is a further manifestation of the lack of spectral signature of the helix labels.

### DISCUSSION

Previous work regarding $^{13}$C-edited infrared spectroscopy of β-sheets focused on labels in every strand of the sheets.[6,7,20–33] In contrast, our work on the antiparallel β-sheets studied labels in every second strand in order to simulate a single label in a β-hairpin structure of individual peptide molecules. Nevertheless, our calculations are in agreement with previous experimental and computational work: our calculated $^{13}$C-band positions are in the same spectral range as previously found for in-register parallel β-sheets and for antiparallel β-sheets.[7,20–22,24,25,27–33] Also observed in experiments and calculation were the strong intensity of the $^{13}$C-band,[6,7,20,21,25,27,29–33] its low wavenumber when the labels are in-register,[6,7,21,25,28,32,33] its upshift upon isotope dilution,[6,7,24,28–30,32] as well as the shifts of the main[6,7,21–23,25,28–32] and of the high wavenumber[7] $^{12}$C-bands upon incorporation of labels in the structure. The concomitant intensity decrease of the main $^{12}$C-band has been seen before in calculations,[6,7,22] but has remained less discussed in the literature. Our work contributes a comprehensive analysis of such spectral properties for different label positions and highlights the role of hydrogen bonding for the observation of a $^{13}$C-band.

### CONCLUSIONS

A prerequisite for observing a $^{13}$C-shoulder or $^{13}$C-band on the low wavenumber side of the main $^{12}$C-band of β-sheets is a low *local* wavenumber of the labeled group. Otherwise, the $^{13}$C-band gets too close to the main $^{12}$C-band, so that the absorption of the $^{13}$C-modes is not resolved. A low local wavenumber can be achieved by hydrogen bonding (or more general by favorable electrostatic interactions)[34,35] and by the influence of the preceding and following amide group. The latter effect depends on the local conformation and generates a downshift for extended conformations but an upshift for helices.[36–39] Thus, an extended conformation and hydrogen bonding, in particular to the carbonyl group, promote the appearance of a distinct $^{13}$C-band. In the absence of hydrogen bonding, the $^{13}$C-mode wavenumbers are even higher than the main $^{12}$C-band wavenumber.

These effects can be seen in our α-hemolysin calculations, where labels in disordered sheet regions and in the loop of the hairpin fail to produce a distinct $^{13}$C-band (Table S13). They are also evident for amide groups that extend from the β-sheet without hydrogen bonding (Table S4) and for the terminal amide groups in the antiparallel β-sheet models (Tables S2, S3, and S6), which have a higher local wavenumber because they lack a sequence neighbor and thus generate only a low wavenumber shoulder but no distinct $^{13}$C-band. In some of these cases a $^{13}$C-band in the second derivative is observed that is upshifted relative to that caused by more central label positions.

When a labeled amide group experiences electrostatic interactions comparable to those in β-sheets, also the wavenumber of the $^{13}$C-mode is similar. However, its intensity is much lower when it is not in a sheet, as shown here for a helix-sheet model. An important reason for the high intensity of the $^{13}$C-band in β-sheets seems to be the coupling with $^{12}$C-groups in nearby strands.

The position of the $^{13}$C-band depends very little on the number of residues in a β-strand and little on the number of strands as long as this number is 8 or larger (Tables S2, S3, and S6). In such extended sheets it reflects mainly the local wavenumber of the labeled amide group. This can be clearly seen in our α-hemolysin results (Table S13) which show a 20 cm$^{-1}$ spread of the $^{13}$C-band position for different label positions that can be correlated with the local wavenumbers of the labeled groups.

The main $^{12}$C-band of β-sheets experiences an upshift of a few cm$^{-1}$ when residues in the core of the β-strands are labeled because the participation of the labeled groups to the $^{12}$C-modes is much reduced. The upshift might vary considerably for different amide groups in a narrow sheet (4-5 amide groups per β-strand) and can reach 10 cm$^{-1}$ (Tables S2 and S4) but is more modest for wider sheets (9 amide groups per strand). It is strongest in central strand regions. In general, $^{13}$C-groups contribute little to the intensity of the main $^{12}$C-band but in some cases, their contribution may be substantial.

The intensity of the $^{13}$C-band seems to decrease with increasing local wavenumber variation of the labeled groups. (see our hemolysin results). On the other hand it does not depend much on the label position in a given regular structure. However, its relative intensity compared that of the main $^{12}$C-band of the same spectrum can vary considerably due to a variation of the $^{12}$C-band intensity. This relative intensity can amount to more than 70% in our calculations (8-stranded sheets with 4 and 5 amide groups per strand, Tables S2 and S4), which is much more than expected from the relative abundance of the $^{13}$C-groups. The relative intensity is larger for narrower sheets, partly because of the higher relative abundance of the $^{13}$C-groups.

The intensity of the main $^{12}$C-band for a labeled structure is reduced relative to that of the completely unlabeled structure when the labels reside within antiparallel β-sheets. The reduction can be drastic, down to ~60%, as observed for sheets with 4 or 5 amide groups per strand (Tables S2 and S4). The reduction is strongest for peripheral label positions that are 1-3 amide groups away from the strand ends depending on the width of the sheet (Tables S2, S3, and S6). It is less in the central parts of the β-strands and closer to the strand ends. Nevertheless, the reduction one amide group away from the ends was always considerable and this is remarkable because the respective $^{12}$C-band shift was very small for the wider sheets (9 amide groups per strand, Tables S3 and S6). Thus these two $^{12}$C-band properties are not correlated. This is also evident for central label positions, which caused larger $^{12}$C-band shifts but smaller $^{12}$C-band reductions than peripheral positions.

The results for parallel β-sheets are similar to those for the antiparallel sheets (Tables S9 to S12) with the exception of in-register label positions in every strand (Table S8) which lead to a strong downshift of the $^{13}$C-band. Such label positions were not of interest for the antiparallel sheets, because they were intended to model a β-hairpin structure.

### AUTHOR INFORMATION


#### Corresponding Author

**Andreas Barth** – *Department of Biochemistry and Biophysics, Stockholm University, Stockholm, Sweden*; orcid.org/0000-0001-5784-7673;

#### Authors

**Faraz Vosough** – *Department of Biochemistry and Biophysics, Stockholm University, Stockholm, Sweden*; orcid.org/0000-0001-5856-3226

**Simon M. Carstensen** – *Department of Biochemistry and Biophysics, Stockholm University, Stockholm, Sweden*

**Cesare M. Baronio** – *Department of Biochemistry and Biophysics, Stockholm University, Stockholm, Sweden*; orcid.org/0000-0003-1399-748X

(22) Setnicka, V.; Huang, R.; Thomas, C. L.; Etienne, M. a; Kubelka, J.; Hammer, R. P.; Keiderling, T. a. IR Study of Cross-Strand Coupling in a β-Hairpin Peptide Using

Isotopic Labels. *J. Am. Chem. Soc.* **2005**, *127* (14), 4992–4993. https://doi.org/10.1021/ja043007f.

### APPENDIX I

This appendix explains the influence of the number of strands on the downshift of the $^{13}$C-band relative to the main $^{12}$C-band of the unlabeled model. Considered are a single strand with a single labeled amide group and sheets with 8 and 24 strands in which every second strand contains a label at the same position. The considered downshift is expressed as

$$\Delta\tilde{\nu}(^{13}\mathrm{C}-^{12}\mathrm{C}_{\mathrm{UL}}) = \tilde{\nu}(^{13}\mathrm{C}) - \tilde{\nu}(^{12}\mathrm{C}_{\mathrm{UL}}),$$

where $\tilde{\nu}(^{13}\mathrm{C})$ is the wavenumber of the $^{13}$C-band and $\tilde{\nu}(^{12}\mathrm{C}_{\mathrm{UL}})$ the wavenumber of the main $^{12}$C-band of the unlabeled model. $\Delta\tilde{\nu}(^{13}\mathrm{C}-^{12}\mathrm{C}_{\mathrm{UL}})$ decreases from −38 cm$^{-1}$ in the single strand to −26 and −23 cm$^{-1}$ for the 8- and 24-stranded sheets, respectively (Tables S1, S3, and S6 in Appendix II). This downshift is the result of two contributions: (i) the local wavenumbers $\tilde{\nu}_0$ of the participating amide groups and (ii) the wavenumber shifts due to vibrational coupling $\Delta\tilde{\nu}_{\mathrm{VC}}$. These contributions are quantified below at the examples of a single strand and of an 8-stranded sheet, both with 10 residues per strand.

Ignoring local wavenumber variations, simple expressions for $\tilde{\nu}(^{13}\mathrm{C})$ and of $\tilde{\nu}(^{12}\mathrm{C}_{\mathrm{UL}})$ can be obtained

$$\tilde{\nu}(^{13}\mathrm{C}) = \tilde{\nu}_0(^{13}\mathrm{C}) + \Delta\tilde{\nu}_{\mathrm{VC}}(^{13}\mathrm{C})$$

$$\tilde{\nu}(^{12}\mathrm{C}_{\mathrm{UL}}) = \tilde{\nu}_0(^{12}\mathrm{C}) + \Delta\tilde{\nu}_{\mathrm{VC}}(^{12}\mathrm{C})$$

where the carbon isotope indicates the $^{13}$C- or the main $^{12}$C-band. This gives for the isotope shift of the $^{13}$C-band

$$\Delta\tilde{\nu}(^{13}\mathrm{C}-^{12}\mathrm{C}_{\mathrm{UL}}) = \tilde{\nu}_0(^{13}\mathrm{C}) + \Delta\tilde{\nu}_{\mathrm{VC}}(^{13}\mathrm{C}) - \tilde{\nu}_0(^{12}\mathrm{C}) - \Delta\tilde{\nu}_{\mathrm{VC}}(^{12}\mathrm{C}).$$

Isotope labeling affects the local wavenumbers $\tilde{\nu}_0$ in the same way in the single strand and in the β-sheets. However, its effect on the coupling-induced wavenumber shifts $\Delta\tilde{\nu}_{\mathrm{VC}}$ is different as outlined in the following.

$^{13}$C-labeling leads to a downshift of the local wavenumber of the labeled group $\tilde{\nu}_0(^{13}\mathrm{C})$ by ~44 cm$^{-1}$ in our calculations:

$$\tilde{\nu}_0(^{13}\mathrm{C}) = \tilde{\nu}_0(^{12}\mathrm{C}) - 44 \text{ cm}^{-1}$$

where $\tilde{\nu}_0(^{12}\mathrm{C})$ is the local wavenumber of the same amide group when it is unlabeled. This gives

$$\Delta\tilde{\nu}(^{13}\mathrm{C}-^{12}\mathrm{C}_{\mathrm{UL}}) = -44 \text{ cm}^{-1} + \Delta\tilde{\nu}_{\mathrm{VC}}(^{13}\mathrm{C}) - \Delta\tilde{\nu}_{\mathrm{VC}}(^{12}\mathrm{C})$$

The local wavenumbers of unlabeled amide groups $\tilde{\nu}_0(^{12}\mathrm{C})$ in the center of the single strand are ~1681 cm$^{-1}$ and of the 8-stranded β-sheet ~1652 cm$^{-1}$. These amide groups couple in the strongest $^{12}$C-modes, which makes the wavenumber of the $^{12}$C-band $\tilde{\nu}(^{12}\mathrm{C}_{\mathrm{UL}})$ lower than the local wavenumber. For the single strand, $\tilde{\nu}(^{12}\mathrm{C}_{\mathrm{UL}}) \approx 1673$ cm$^{-1}$ (Table S1) giving

$$\Delta\tilde{\nu}_{\mathrm{VC}}(^{12}\mathrm{C}) \approx -8 \text{ cm}^{-1}$$

and for the 8-stranded sheet, $\tilde{\nu}(^{12}\mathrm{C}_{\mathrm{UL}}) \approx 1626$ cm$^{-1}$ (Table S3) giving

$$\Delta\tilde{\nu}_{\mathrm{VC}}(^{12}\mathrm{C}) \approx -26 \text{ cm}^{-1}.$$

Labeling generates local wavenumbers $\tilde{\nu}_0(^{13}\mathrm{C})$ of ~1636 and ~1608 cm$^{-1}$ for the single strand and the 8-stranded sheet, respectively. As for the $^{12}$C-modes, vibrational coupling lowers the wavenumber of the $^{13}$C-modes: for the single strand, $\tilde{\nu}(^{13}\mathrm{C}) \approx 1635$ cm$^{-1}$ (Table S1) giving

$$\Delta\tilde{\nu}_{\mathrm{VC}}(^{13}\mathrm{C}) \approx -1 \text{ cm}^{-1}$$

and for the 8-stranded sheet $\tilde{\nu}(^{13}\mathrm{C}) \approx 1600$ cm$^{-1}$ giving

$$\Delta\tilde{\nu}_{\mathrm{VC}}(^{13}\mathrm{C}) \approx -8 \text{ cm}^{-1}.$$

The downshift due to coupling is much smaller than for the strongest $^{12}$C-modes because the $^{12}$C-groups couple more effectively with other $^{12}$C-groups than with $^{13}$C-groups. The downshift is more reduced for the 8-stranded sheet than for the single strand. Using the above approximate numbers for $\Delta\tilde{\nu}_{\mathrm{VC}}$ we get for the isotope shifts

$$\Delta\tilde{\nu}(^{13}\mathrm{C}-^{12}\mathrm{C}_{\mathrm{UL}}, \text{single strand}) = (-44 -1 + 8) \text{ cm}^{-1} = -37 \text{ cm}^{-1}$$

$$\Delta\tilde{\nu}(^{13}\mathrm{C}-^{12}\mathrm{C}_{\mathrm{UL}}, \text{sheet}) = (-44 -8 +26) \text{ cm}^{-1} = -26 \text{ cm}^{-1}$$

Thus, the stronger reduction of the vibrational coupling effect on the $^{13}$C-modes of the 8-stranded sheet than on those of the single strand explains the larger isotope shifts of the latter. The reduction is even stronger for the 24-stranded sheet.

### APPENDIX II

**Table S1**. Spectral parameters for an isotope-labeled **single** strand of an **antiparallel** β-sheet with **10** residues (9 amide groups).

| Label positions | $\tilde{\nu}(^{13}C)$ | $\tilde{\nu}(^{12}C)$ | $\Delta\tilde{\nu}(^{13}C-^{12}C_{UL})$ | $\Delta\tilde{\nu}(^{12}C_L-^{12}C_{UL})$ | $I(^{13}C)$ | $I(^{12}C)$ | $I(^{13}C)/I(^{12}C)$ | $I(^{12}C_L)/I(^{12}C_{UL})$ |
|---|---|---|---|---|---|---|---|---|
| Unlabeled | 1650.0 | 1672.9 | | | 0.0 | 2.3 | | |
| 1 | 1640.5 | 1672.9 | -32.4 | 0.0 | 0.4 | 2.0 | 18 | 89 |
| 2 | 1634.2 | 1673.2 | -38.7 | 0.3 | 0.4 | 1.7 | 26 | 75 |
| 3 | 1635.6 | 1674.7 | -37.3 | 1.8 | 0.4 | 1.8 | 25 | 78 |
| 4 | 1635.1 | 1674.2 | -37.8 | 1.3 | 0.4 | 1.8 | 24 | 81 |
| 5 | 1635.3 | 1674.1 | -37.6 | 1.2 | 0.4 | 1.9 | 24 | 82 |
| 6 | 1635.2 | 1674.6 | -37.7 | 1.7 | 0.4 | 1.8 | 24 | 81 |
| 7 | 1635.1 | 1674.3 | -37.8 | 1.4 | 0.4 | 1.7 | 26 | 75 |
| 8 | 1635.4 | 1672.9 | -37.5 | 0.0 | 0.4 | 1.8 | 23 | 81 |
| 9 | 1650.0 | 1672.7 | | -0.2 | 0.3 | 2.1 | | 93 |
| Average IE (3-7) | | | -37.6 | 1.5 | | | 25 | 79 |

The residue numbers of the labeled carbonyl groups are indicated in the left column. Missing numbers in the table indicate that no distinct $^{13}$C-band was observed.
Average IE: average isotope effect for amide groups in the core of the sheet. The bracket indicates the amide groups in each strand that were considered to be in the core.
$\tilde{\nu}(^{13}C)$: wavenumber of the $^{13}$C-band in cm$^{-1}$ (black) evaluated from the calculated spectrum or a convenient wavenumber (gray) used to evaluate the intensity on the low wavenumber side of the $^{12}$C-band.
$\tilde{\nu}(^{12}C)$: wavenumber of the main $^{12}$C-band in cm$^{-1}$ evaluated from the calculated spectrum
$\Delta\tilde{\nu}(^{13}C-^{12}C_{UL})$: wavenumber difference of the $^{13}$C-band to the main $^{12}$C-band of the unlabeled peptide in cm$^{-1}$. Missing numbers in the table indicate that no distinct $^{13}$C-band was observed.
$\Delta\tilde{\nu}(^{12}C_L-^{12}C_{UL})$: wavenumber shift of the main $^{12}$C-band due to labeling in cm$^{-1}$ (wavenumber of the main $^{12}$C-band in the presence of one or more labeled groups minus $^{12}$C-band wavenumber in the absence of a labeled group)
$I(^{13}C)$: intensity at the maximum of the $^{13}$C-band (black) or intensity at the specified wavenumber on the low wavenumber side of the $^{12}$C-band (gray). The gray numbers are meant to give an indication of the $^{13}$C-absorption in cases where no distinct $^{13}$C-band was observed. The intensity integral of the spectrum of a normal mode is equal to its squared dipole derivative in D Å$^{-1}$ u$^{-1/2}$.
$I(^{12}C)$: intensity at the maximum of the $^{12}$C-band
$I(^{13}C)/I(^{12}C)$: maximum intensity of the $^{13}$C-band divided by the $^{12}$C-band intensity of the same labeled model in %
$I(^{12}C_L)/I(^{12}C_{UL})$: maximum intensity of the main $^{12}$C-band in the presence of a labeled group divided by the $^{12}$C-band intensity in the absence of a labeled group in %

**Table S2**. Spectral parameters for an isotope-labeled **8**-stranded **antiparallel** β-sheet with **6** residues per strand (5 amide groups) as shown in Figure S2. Labels were placed in every **second** strand.

| Label positions | $\tilde{\nu}(^{13}C)$ | $\tilde{\nu}(^{12}C)$ | $\tilde{\nu}(^{12}C,hw)$ | $\Delta\tilde{\nu}(^{13}C-^{12}C_{UL})$ | $\Delta\tilde{\nu}(^{12}C_L-^{12}C_{UL})$ | $I(^{13}C)$ | $I(^{12}C)$ | $I(^{12}C,hw)$ | $I(^{13}C)/I(^{12}C)$ | $I(^{12}C_L)/I(^{12}C_{UL})$ | $I(^{12}C,hw_L)/I(^{12}C,hw_{UL})$ |
|---|---|---|---|---|---|---|---|---|---|---|---|
| Unlabeled | 1610.0 | 1626.5 | 1676.8 | | | 0.4 | 11.8 | **2.0** | | | |
| 1, 13, 25, 37 | 1610.0 | 1626.3 | 1672.0 | | -0.2 | 2.6 | 11.4 | 2.1 | | 96 | |
| 2, 14, 26, 38 | 1600.2 | 1628.4 | 1673.2 | -26.3 | 1.9 | 4.2 | 7.5 | 2.0 | 56 | 63 | 103 |
| 3, 15, 27, 39 | 1600.4 | 1635.0 | 1671.6 | -26.1 | 8.5 | 4.5 | 8.3 | 2.2 | 54 | 70 | 113 |
| 4, 16, 28, 40 | 1600.7 | 1630.1 | 1673.0 | -25.8 | 3.6 | 4.5 | 6.3 | 2.0 | 71 | 54 | 99 |
| 5, 17, 29, 41 | 1610.0 | 1625.8 | 1677.0 | | -0.7 | 3.5 | 9.4 | 1.8 | | 79 | 93 |
| 7, 19, 31, 43 | 1610.0 | 1626.5 | 1675.5 | | 0.0 | 2.9 | 10.8 | 1.9 | | 91 | 99 |
| 8, 20, 32, 44 | 1600.0 | 1627.7 | 1672.0 | -26.5 | 1.2 | 3.9 | 7.8 | 2.1 | 50 | 66 | 108 |
| 9, 21, 33, 45 | 1600.1 | 1635.5 | 1671.3 | -26.4 | 9.0 | 4.7 | 9.0 | 2.2 | 53 | 76 | 112 |
| 10, 22, 34, 46 | 1601.3 | 1629.1 | 1673.3 | -25.2 | 2.6 | 4.3 | 7.0 | 2.0 | 61 | 59 | 103 |
| 11, 23, 35, 47 | 1610.0 | 1625.5 | 1677.3 | | -1.0 | 3.9 | 9.7 | 1.9 | | 82 | 95 |
| Average IE (2-4) | | | | -26.1 | 4.5 | | | | 58 | 65 | 106 |

$\tilde{\nu}(^{12}C,hw)$: Wavenumber of the high wavenumber band (black) or wavenumber close to a high wavenumber shoulder (gray), both evaluated from the spectrum
$I(^{12}C,hw)$: Intensity of the high wavenumber band (black) or intensity at the specified wavenumber close to the high wavenumber shoulder (gray).
$I(^{12}C,hw_L)/I(^{12}C,hw_{UL})$: Intensity of the high wavenumber band in the presence of a labeled group divided by the band intensity in the absence of a labeled group in %. Missing numbers in the table indicate that no distinct high wavenumber band was observed.
See Table S1 for further explanation.

**Table S3**. Spectral parameters for an isotope-labeled **8**-stranded **antiparallel** β-sheet with **10** residues per strand (9 amide groups) as shown in Figure S1. Labels were placed in every **second** strand.

| Label positions | $\tilde{\nu}(^{13}C)$ | $\tilde{\nu}(^{12}C)$ | $\tilde{\nu}(^{12}C,hw)$ | $\Delta\tilde{\nu}(^{13}C-^{12}C_{UL})$ | $\Delta\tilde{\nu}(^{12}C_L-^{12}C_{UL})$ | $I(^{13}C)$ | $I(^{12}C)$ | $I(^{12}C,hw)$ | $I(^{13}C)/I(^{12}C)$ | $I(^{12}C_L)/I(^{12}C_{UL})$ | $I(^{12}C,hw_L)/I(^{12}C,hw_{UL})$ |
|---|---|---|---|---|---|---|---|---|---|---|---|
| Unlabeled | 1610.0 | 1626.1 | 1678.7 | | | 1.2 | 24.2 | 3.5 | | | |
| 1, 21, 41, 61 | 1610.0 | 1625.8 | 1677.5 | | -0.3 | 3.2 | 23.9 | 3.4 | | 99 | 97 |
| 2, 22, 42, 62 | 1600.4 | 1625.8 | 1677.7 | -25.7 | -0.3 | 4.0 | 19.6 | 3.4 | 20 | 81 | 97 |
| 3, 23, 43, 63 | 1600.2 | 1626.6 | 1677.0 | -25.9 | 0.5 | 4.2 | 17.7 | 3.4 | 24 | 73 | 98 |
| 4, 24, 44, 64 | 1600.1 | 1628.0 | 1677.6 | -26.0 | 1.9 | 4.5 | 19.1 | 3.5 | 24 | 79 | 101 |
| 5, 25, 45, 65 | 1600.3 | 1627.3 | 1677.2 | -25.8 | 1.2 | 4.3 | 20.4 | 3.5 | 21 | 84 | 99 |
| 6, 26, 46, 66 | 1599.9 | 1627.9 | 1677.5 | -26.2 | 1.8 | 4.5 | 19.6 | 3.5 | 23 | 81 | 100 |
| 7, 27, 47, 67 | 1600.5 | 1627.6 | 1676.5 | -25.6 | 1.5 | 4.3 | 18.4 | 3.4 | 23 | 76 | 98 |
| 8, 28, 48, 68 | 1600.7 | 1625.8 | 1677.9 | -25.4 | -0.3 | 4.3 | 18.0 | 3.3 | 24 | 75 | 94 |
| 9, 29, 49, 69 | 1610.0 | 1625.2 | 1678.9 | | -0.9 | 4.0 | 22.3 | 3.4 | | 92 | 96 |
| 11, 31, 51, 71 | 1610.0 | 1625.8 | 1678.2 | | -0.3 | 3.6 | 23.3 | 3.4 | | 96 | 97 |
| 12, 32, 52, 72 | 1600.0 | 1625.5 | 1677.0 | -26.1 | -0.6 | 3.7 | 20.1 | 3.3 | 18 | 83 | 95 |
| 13, 33, 53, 73 | 1600.3 | 1627.2 | 1677.3 | -25.8 | 1.1 | 4.5 | 17.1 | 3.5 | 26 | 71 | 99 |
| 14, 34, 54, 74 | 1600.1 | 1627.6 | 1677.1 | -26.0 | 1.5 | 4.2 | 19.5 | 3.5 | 22 | 81 | 100 |
| 15, 35, 55, 75 | 1600.1 | 1627.7 | 1677.7 | -26.0 | 1.6 | 4.5 | 20.0 | 3.5 | 23 | 83 | 101 |
| 16, 36, 56, 76 | 1600.3 | 1627.5 | 1677.1 | -25.8 | 1.4 | 4.3 | 20.0 | 3.4 | 21 | 83 | 98 |
| 17, 37, 57, 77 | 1599.9 | 1628.2 | 1677.2 | -26.2 | 2.1 | 4.5 | 17.6 | 3.5 | 26 | 73 | 99 |
| 18, 38, 58, 78 | 1601.6 | 1625.8 | 1677.5 | -24.5 | -0.3 | 4.0 | 19.0 | 3.3 | 21 | 79 | 95 |
| 19, 39, 59, 79 | 1610.0 | 1625.1 | 1679.0 | | -1.0 | 4.4 | 22.7 | 3.4 | | 94 | 97 |
| Average IE (2-8) | | | | -25.8 | 0.9 | | | | 23 | 79 | 98 |
| Average IE (3-7) | | | | -25.9 | 1.5 | | | | 23 | 78 | 99 |

See Tables S1 and S2 for further explanation.

**Table S4**. Spectral parameters for an isotope-labeled **antiparallel** β-sheet with **8 out-of-register** strands and **10** residues/strand of which 5 residues/strand (4 amide groups) are in the central part of the structure as shown in Figure S3. Labels were placed in every **second** strand.

| Label positions | $\tilde{\nu}(^{13}C)$ | $\tilde{\nu}(^{12}C)$ | $\Delta\tilde{\nu}(^{13}C-^{12}C_{UL})$ | $\Delta\tilde{\nu}(^{12}C_L-^{12}C_{UL})$ | $I(^{13}C)$ | $I(^{12}C)$ | $I(^{13}C)/I(^{12}C)$ | $I(^{12}C_L)/I(^{12}C_{UL})$ |
|---|---|---|---|---|---|---|---|---|
| Unlabeled | 1610.0 | 1627.9 | | | 0.2 | 10.0 | | |
| 1, 21, 41, 61 | 1610.0 | 1628.4 | | 0.5 | 0.2 | 10.4 | | 104 |
| 2, 22, 42, 62 | 1610.0 | 1628.7 | | 0.8 | 0.2 | 11.3 | | 113 |
| 3, 23, 43, 63 | 1610.0 | 1628.8 | | 0.9 | 0.2 | 11.1 | | 111 |
| 4, 24, 44, 64 | 1610.0 | 1628.4 | | 0.5 | 0.2 | 10.5 | | 105 |
| 5, 25, 45, 65 | 1610.0 | 1627.0 | | -0.9 | 0.7 | 11.0 | | 111 |
| 6, 26, 46, 66 | 1602.7 | 1627.8 | -25.2 | -0.1 | 2.9 | 8.2 | 36 | 82 |
| 7, 27, 47, 67 | 1600.2 | 1637.5 | -27.7 | 9.6 | 4.1 | 5.9 | 71 | 59 |
| 8, 28, 48, 68 | 1600.7 | 1638.4 | -27.2 | 10.5 | 4.0 | 7.2 | 56 | 72 |
| 9, 29, 49, 69 | 1610.0 | 1621.8 | | -6.1 | 2.8 | 7.5 | | 76 |
| 11, 31, 51, 71 | 1610.0 | 1628.4 | | 0.5 | 0.2 | 10.4 | | 104 |
| 12, 32, 52, 72 | 1610.0 | 1628.7 | | 0.8 | 0.2 | 11.3 | | 113 |
| 13, 33, 53, 73 | 1610.0 | 1628.8 | | 0.9 | 0.2 | 11.1 | | 111 |
| 14, 34, 54, 74 | 1610.0 | 1628.4 | | 0.5 | 0.2 | 10.5 | | 105 |
| 15, 35, 55, 75 | 1610.0 | 1627.0 | | -0.9 | 0.7 | 11.0 | | 111 |
| 16, 36, 56, 76 | 1602.7 | 1627.8 | -25.2 | -0.1 | 2.9 | 8.2 | 36 | 82 |
| 17, 37, 57, 77 | 1600.1 | 1637.5 | -27.8 | 9.6 | 4.1 | 5.9 | 71 | 59 |
| 18, 38, 58, 78 | 1600.7 | 1638.4 | -27.2 | 10.5 | 4.0 | 7.2 | 56 | 72 |
| 19, 39, 59, 79 | 1610.0 | 1621.8 | | -6.1 | 2.8 | 7.5 | | 76 |
| Average IE (6-8) | | | -26.7 | 6.7 | | | 54 | 71 |

Residues 6-10, 16-20, 26-30 are inside the β-sheet while the others are outside. See Table S1 for further explanation.

**Table S5**. Spectral effects of isotope-dilution in a **9**-stranded **antiparallel** β-sheet with **6** residues per strand (5 amide groups). The sheet contained either a **single label**, or labels in every **second** inner strand.

| Label positions | $\tilde{\nu}(^{13}C)$ | $\tilde{\nu}(^{12}C)$ | $\tilde{\nu}(^{12}C,hw)$ | $\Delta\tilde{\nu}(^{13}C-^{12}C_{UL})$ | $\Delta\tilde{\nu}(^{12}C_L-^{12}C_{UL})$ | $I(^{13}C)$ | $I(^{12}C)$ | $I(^{12}C,hw)$ | $I(^{13}C)/I(^{12}C)$ | $I(^{12}C_L)/I(^{12}C_{UL})$ | $I(^{12}C,hw_L)/I(^{12}C,hw_{UL})$ |
|---|---|---|---|---|---|---|---|---|---|---|---|
| Unlabeled | 1610.0 | 1625.8 | 1676.9 | | | 0.7 | 13.5 | 2.2 | | | |
| 20 | 1603.1 | 1627.0 | 1675.8 | -22.7 | 1.2 | 1.9 | 11.4 | 2.2 | 17 | 84 | 100 |
| 21 | 1602.8 | 1629.4 | 1675.6 | -23.0 | 3.6 | 2.3 | 12.1 | 2.2 | 19 | 89 | 100 |
| 22 | 1604.2 | 1628.0 | 1675.9 | -21.6 | 2.2 | 2.2 | 11.2 | 2.2 | 19 | 83 | 99 |
| 25 | 1610.0 | 1625.9 | 1676.4 | -15.8 | 0.1 | 1.5 | 13.0 | 2.2 | | 96 | 100 |
| 26 | 1603.0 | 1627.3 | 1675.9 | -22.8 | 1.5 | 1.9 | 11.4 | 2.2 | 17 | 84 | 99 |
| 27 | 1603.0 | 1629.6 | 1675.6 | -22.8 | 3.8 | 2.3 | 12.4 | 2.3 | 19 | 91 | 101 |
| 28 | 1603.9 | 1628.2 | 1675.8 | -21.9 | 2.4 | 2.2 | 11.4 | 2.2 | 19 | 84 | 99 |
| 29 | 1610.0 | 1625.6 | 1676.8 | -15.8 | -0.2 | 1.9 | 12.3 | 2.2 | | 91 | 98 |
| 7, 19, 31, 43 | 1610.0 | 1625.9 | 1673.6 | -15.8 | 0.1 | 3.5 | 12.1 | 2.3 | | 89 | 103 |
| 8, 20, 32, 44 | 1599.6 | 1627.9 | 1673.7 | -26.2 | 2.1 | 4.9 | 8.4 | 2.2 | 58 | 62 | 96 |
| 9, 21, 33, 45 | 1599.7 | 1635.1 | 1672.3 | -26.1 | 9.3 | 5.6 | 10.1 | 2.4 | 55 | 75 | 105 |
| 10, 22, 34, 46 | 1600.5 | 1629.5 | 1671.9 | -25.3 | 3.7 | 5.3 | 7.3 | 2.2 | 73 | 54 | 99 |
| 11, 23, 35, 47 | 1610.0 | 1624.8 | 1677.0 | -15.8 | -1.0 | 4.9 | 10.6 | 2.1 | | 78 | 94 |
| Average IE (20-22, 26-28)[1] | | | | -22.5 | 2.5 | | | | 18 | 86 | 100 |
| Average IE (8-10)[2] | | | | -25.9 | 5.0 | | | | 62 | 64 | 100 |

See Tables S1 and S2 for further explanation.

[1] Average isotope effect for single labels in the sheet. The numbers in the bracket indicate the residue numbers of the labeled groups that were considered for averaging.

[2] Average isotope effect for multiple labels in the sheet. The numbers in the bracket indicate the numbers of the first labeled residue that were considered for averaging. For example, 8 stands for residues 8, 20, 32, and 44.

**Table S6**. Spectral parameters for an isotope-labeled **24**-stranded **antiparallel** β-sheet with **10** residues per strand (9 amide groups) of which the first 8 strands are shown in Figure S1. Labels were placed in every **second** strand.

| Label positions | $\tilde{\nu}(^{13}C)$ | $\tilde{\nu}(^{12}C)$ | $\tilde{\nu}(^{12}C,hw)$ | $\Delta\tilde{\nu}(^{13}C-^{12}C_{UL})$ | $\Delta\tilde{\nu}(^{12}C_L-^{12}C_{UL})$ | $I(^{13}C)$ | $I(^{12}C)$ | $I(^{12}C,hw)$ | $I(^{13}C)/I(^{12}C)$ | $I(^{12}C_L)/I(^{12}C_{UL})$ | $I(^{12}C,hw_L)/I(^{12}C,hw_{UL})$ |
|---|---|---|---|---|---|---|---|---|---|---|---|
| Unlabeled | 1605.0 | 1621.3 | 1678.3 | | | 3.9 | 78.8 | 11.4 | | | |
| 1, 21, ...221 | 1605.0 | 1621.1 | 1677.6 | | -0.2 | 13.6 | 74.5 | 11.1 | | 95 | 97 |
| 2, 22, ...222 | 1598.8 | 1621.3 | 1677.2 | -22.5 | 0.0 | 15.6 | 62.4 | 10.8 | 25 | 79 | 95 |
| 3, 23, ...223 | 1598.4 | 1622.3 | 1676.9 | -22.9 | 1.0 | 17.6 | 54.2 | 11.2 | 33 | 69 | 98 |
| 4, 24, ...224 | 1598.2 | 1624.2 | 1677.1 | -23.1 | 2.9 | 18.5 | 60.3 | 11.5 | 31 | 77 | 101 |
| 5, 25, ...225 | 1598.2 | 1624.0 | 1677.1 | -23.1 | 2.7 | 18.4 | 65.3 | 11.4 | 28 | 83 | 100 |
| 6, 26, ...226 | 1598.2 | 1624.2 | 1677.1 | -23.1 | 2.9 | 18.6 | 62.5 | 11.3 | 30 | 79 | 99 |
| 7, 27, ...227 | 1598.3 | 1623.5 | 1676.7 | -23.0 | 2.2 | 17.9 | 55.4 | 11.1 | 32 | 70 | 97 |
| 8, 28, ...228 | 1599.4 | 1621.3 | 1677.4 | -21.9 | 0.0 | 16.9 | 58.5 | 10.6 | 29 | 74 | 93 |
| 9, 29, ...229 | 1605.0 | 1619.9 | 1678.3 | | -1.4 | 11.6 | 74.2 | 10.8 | | 94 | 95 |
| 11, 31, ...231 | 1605.0 | 1621.0 | 1677.8 | | -0.3 | 13.8 | 74.1 | 11.1 | | 94 | 97 |
| 12, 32, ...232 | 1598.6 | 1621.3 | 1677.1 | -22.7 | 0.0 | 15.3 | 63.1 | 10.7 | 24 | 80 | 94 |
| 13, 33, ...233 | 1598.6 | 1621.7 | 1677.0 | -22.7 | 0.4 | 16.7 | 58.6 | 11.0 | 28 | 74 | 96 |
| 14, 34, ...234 | 1598.2 | 1624.1 | 1677.0 | -23.1 | 2.8 | 18.2 | 61.2 | 11.4 | 30 | 78 | 100 |
| 15, 35, ...235 | 1598.2 | 1624.0 | 1677.2 | -23.1 | 2.7 | 18.7 | 64.5 | 11.4 | 29 | 82 | 100 |
| 16, 36, ...236 | 1598.3 | 1624.1 | 1677.1 | -23.0 | 2.8 | 18.4 | 63.3 | 11.2 | 29 | 80 | 98 |
| 17, 37, ...237 | 1598.2 | 1623.5 | 1676.8 | -23.1 | 2.2 | 18.1 | 54.3 | 11.1 | 33 | 69 | 98 |
| 18, 38, ...238 | 1599.6 | 1621.4 | 1677.3 | -21.7 | 0.1 | 16.6 | 59.2 | 10.6 | 28 | 75 | 93 |
| 19, 39, ...239 | 1605.0 | 1619.9 | 1678.3 | | -1.4 | 11.9 | 74.4 | 10.9 | | 94 | 95 |
| Average IE (2-8) | | | | -22.8 | 1.6 | | | | 29 | 76 | 97 |
| Average IE (3-7) | | | | -23.0 | 2.3 | | | | 30 | 76 | 99 |

The first two and the last labeled residues are indicated in the left column: for example ”2, 22, ...222” means that residues 2, 22, 42, 62, 82, 102, 122, 142, 162, 182, 202 and 222 were labeled. See Tables S1 and S2 for further explanation.

**Table S7**. Spectral parameters for an isotope-labeled **24**-stranded **antiparallel** β-sheet with **10** residues per strand (9 amide groups) of which the first 8 strands are shown in Figure S1. In contrast to the other antiparallel β-sheets, only every **third** strand contained a label.

| Label positions | $\tilde{\nu}(^{13}C)$ | $\tilde{\nu}(^{12}C)$ | $\Delta\tilde{\nu}(^{13}C-^{12}C_{UL})$ | $\Delta\tilde{\nu}(^{12}C_L-^{12}C_{UL})$ | $I(^{13}C)$ | $I(^{12}C)$ | $I(^{13}C)/I(^{12}C)$ | $I(^{12}C_L)/I(^{12}C_{UL})$ |
|---|---|---|---|---|---|---|---|---|
| Unlabeled | 1605.0 | 1621.3 | | | 3.9 | 78.8 | | |
| 1, 31, ...211 | 1605.0 | 1621.1 | | -0.2 | 10.4 | 75.7 | | 96 |
| 2, 32, ...212 | 1605.0 | 1621.7 | | 0.4 | 16.8 | 60.6 | | 77 |
| 3, 33, ...213 | 1603.2 | 1624.3 | -18.1 | 3.0 | 18.8 | 59.4 | 32 | 75 |
| 4, 24, ...214 | 1602.2 | 1624.6 | -19.1 | 3.3 | 18.2 | 65.9 | 28 | 84 |
| 5, 35, ...215 | 1600.9 | 1623.7 | -20.4 | 2.4 | 15.6 | 68.1 | 23 | 86 |
| 6, 36, ...216 | 1602.0 | 1624.4 | -19.3 | 3.1 | 17.8 | 66.3 | 27 | 84 |
| 7, 37, ...217 | 1602.7 | 1624.7 | -18.6 | 3.4 | 18.8 | 62.8 | 30 | 80 |
| 8, 38, ...218 | 1605.0 | 1622.4 | | 1.1 | 19.3 | 56.7 | | 72 |
| 9, 39, ...219 | 1605.0 | 1619.9 | | -1.4 | 7.9 | 76.2 | | 97 |
| 11, 41, ...221 | 1605.0 | 1621.0 | | -0.3 | 11.1 | 74.9 | | 95 |
| 12, 42, ...222 | 1605.0 | 1621.8 | | 0.5 | 17.4 | 60.1 | | 76 |
| 13, 43, ...223 | 1603.2 | 1624.7 | -18.1 | 3.4 | 20.1 | 58.4 | 34 | 74 |
| 14, 44, ...224 | 1602.0 | 1624.6 | -19.3 | 3.3 | 18.9 | 65.7 | 29 | 83 |
| 15, 45, ...225 | 1600.9 | 1623.8 | -20.4 | 2.5 | 16.5 | 67.0 | 25 | 85 |
| 16, 46, ...226 | 1601.8 | 1624.4 | -19.5 | 3.1 | 18.4 | 66.0 | 28 | 84 |
| 17, 47, ...227 | 1602.8 | 1625.0 | -18.5 | 3.7 | 20.0 | 61.8 | 32 | 78 |
| 18, 48, ...228 | 1605.0 | 1622.4 | | 1.1 | 19.7 | 56.5 | | 72 |
| 19, 49, ...229 | 1605.0 | 1619.8 | | -1.5 | 8.1 | 76.7 | | 97 |
| 21, 51, ...231 | 1605.0 | 1621.0 | | -0.3 | 10.6 | 75.3 | | 96 |
| 22, 52, ...232 | 1605.0 | 1621.7 | | 0.4 | 16.4 | 61.7 | | 78 |
| 23, 53, ...233 | 1603.4 | 1624.5 | -17.9 | 3.2 | 19.3 | 58.6 | 33 | 74 |
| 24, 54, ...234 | 1602.0 | 1624.4 | -19.3 | 3.1 | 17.8 | 66.6 | 27 | 85 |
| 25, 55, ...235 | 1601.0 | 1623.8 | -20.3 | 2.5 | 16.0 | 67.2 | 24 | 85 |
| 26, 56, ...236 | 1601.8 | 1624.2 | -19.5 | 2.9 | 17.3 | 67.1 | 26 | 85 |
| 27, 57, ...237 | 1603.0 | 1624.9 | -18.3 | 3.6 | 19.3 | 61.9 | 31 | 79 |
| 28, 58, ...238 | 1605.0 | 1622.4 | | 1.1 | 18.5 | 58.3 | | 74 |
| 29, 59, ...239 | 1605.0 | 1619.9 | | -1.4 | 7.9 | 77.1 | | 98 |
| Average IE (3-7) | | | -19.1 | 3.1 | | | 29 | 81 |

The first two and the last labeled residues are indicated on the top of the columns: for example ”2, 32, ...212” means that residues 2, 32, 62, 92, 122, 152, 182, and 212 were labeled. See Table S1 for further explanation.

**Table S8**. Spectral parameters for an isotope-labeled **parallel** β-sheet with **8** strands and **10** residues (9 amide groups) per strand as shown in the top panel of Figure S4. **All** strands contained a labeled amide group.

| Label positions | $\tilde{\nu}(^{13}C)$ | $\tilde{\nu}(^{12}C)$ | $\Delta\tilde{\nu}(^{13}C-^{12}C_{UL})$ | $\Delta\tilde{\nu}(^{12}C_L-^{12}C_{UL})$ | $I(^{13}C)$ | $I(^{12}C)$ | $I(^{13}C)/I(^{12}C)$ | $I(^{12}C_L)/I(^{12}C_{UL})$ |
|---|---|---|---|---|---|---|---|---|
| Unlabeled | 1610.0 | 1633.1 | | | 0.0 | 26.2 | | |
| 1, 11, ...71 | 1595.5 | 1633.1 | -37.6 | 0.0 | 4.0 | 22.2 | 18 | 85 |
| 2, 12, ...72 | 1596.2 | 1633.6 | -36.9 | 0.5 | 4.7 | 19.4 | 24 | 74 |
| 3, 13, ...73 | 1596.9 | 1634.3 | -36.2 | 1.2 | 4.8 | 20.8 | 23 | 79 |
| 4, 14, ...74 | 1596.0 | 1633.9 | -37.1 | 0.8 | 4.6 | 21.2 | 22 | 81 |
| 5, 15, ...75 | 1596.4 | 1634.0 | -36.7 | 0.9 | 4.7 | 21.5 | 22 | 82 |
| 6, 16, ...76 | 1596.0 | 1634.4 | -37.1 | 1.3 | 4.5 | 20.9 | 22 | 80 |
| 7, 17, ...77 | 1596.3 | 1633.9 | -36.8 | 0.8 | 4.7 | 19.3 | 24 | 74 |
| 8, 18, ...78 | 1596.7 | 1632.7 | -36.4 | -0.4 | 4.2 | 21.6 | 19 | 82 |
| 9, 19, ...79 | 1610.0 | 1632.7 | | -0.4 | 2.4 | 25.2 | | 96 |
| Average IE (3-7) | | | -36.8 | 1.0 | | | 23 | 79 |

The first two and the last labeled residues are indicated in the left column: for example ”1, 11, ...71” means that residues 1, 11, 21, 31, 41, 51, 61, and 71 were labeled. See Table S1 for further explanation.

**Table S9**. Spectral parameters for an isotope-labeled **parallel** β-sheet with **8** strands and **10** residues (9 amide groups) per strand as shown in the top panel of Figure S4. Every **second** strand contained a labeled amide group.

| Label positions | $\tilde{\nu}(^{13}C)$ | $\tilde{\nu}(^{12}C)$ | $\Delta\tilde{\nu}(^{13}C-^{12}C_{UL})$ | $\Delta\tilde{\nu}(^{12}C_L-^{12}C_{UL})$ | $I(^{13}C)$ | $I(^{12}C)$ | $I(^{13}C)/I(^{12}C)$ | $I(^{12}C_L)/I(^{12}C_{UL})$ |
|---|---|---|---|---|---|---|---|---|
| Unlabeled | | 1633.1 | | | | 26.2 | | |
| 3, 23, 43, 63 | 1605.8 | 1634.5 | -27.3 | 1.4 | 4.2 | 21.1 | 20 | 81 |
| 5, 25, 45, 65 | 1605.2 | 1634.1 | -27.9 | 1.0 | 4.0 | 21.8 | 18 | 83 |
| 7, 27, 47, 67 | 1605.0 | 1634.2 | -28.1 | 1.1 | 3.9 | 19.8 | 20 | 76 |
| 13, 33, 53, 73 | 1605.8 | 1634.1 | -27.3 | 1.0 | 3.8 | 21.7 | 18 | 83 |
| 15, 35, 55, 75 | 1605.5 | 1633.9 | -27.6 | 0.8 | 3.7 | 22.2 | 17 | 85 |
| 17,37,57,77 | 1605.5 | 1633.9 | -27.6 | 0.8 | 3.7 | 20.3 | 18 | 78 |
| Average IE (3-7) | | | -27.6 | 1.0 | | | 19 | 81 |

See Table S1 for further explanation.

**Table S10**. Spectral parameters for an isotope-labeled **parallel** β-sheet with **8** in-register strands and **10** residues (9 amide groups) per strand as shown in the top panel of Figure S4. **All** strands contained a labeled amide group that were **out-of-register**.

| Label positions | $\tilde{\nu}(^{13}C)$ | $\tilde{\nu}(^{12}C)$ | $\Delta\tilde{\nu}(^{13}C-^{12}C_{UL})$ | $\Delta\tilde{\nu}(^{12}C_L-^{12}C_{UL})$ | $I(^{13}C)$ | $I(^{12}C)$ | $I(^{13}C)/I(^{12}C)$ | $I(^{12}C_L)/I(^{12}C_{UL})$ |
|---|---|---|---|---|---|---|---|---|
| Unlabeled | 1610.0 | 1633.1 | | | 0.0 | 26.2 | | |
| 1,12,23,34,45,56,67,78 | 1604.8 | 1636.0 | -28.3 | 2.9 | 7.6 | 17.7 | 43 | 67 |
| 2,13,24,35,46,57,68,79 | 1605.3 | 1635.6 | -27.8 | 2.5 | 6.8 | 18.0 | 38 | 69 |
| 8,17,26,35,44,53,62,71 | 1605.2 | 1635.8 | -27.9 | 2.7 | 7.1 | 18.3 | 39 | 70 |
| 9,18,27,36,45,54,63,72 | 1604.8 | 1636.0 | -28.3 | 2.9 | 7.1 | 18.3 | 39 | 70 |
| Average IE (1-9) | | | -28.1 | 2.8 | | | 40 | 69 |

See Table S1 for further explanation.

**Table S11**. Spectral parameters for an isotope-labeled **parallel** β-sheet with **8** strands and **10** residues (9 amide groups) per strand as shown in the middle panel of Figure S4. **All** strands contained a labeled amide group that were **out-of-register** because each following strand is shifted by one residue towards the C-terminus of the previous strand.

| Label positions | $\tilde{\nu}(^{13}C)$ | $\tilde{\nu}(^{12}C)$ | $\Delta\tilde{\nu}(^{13}C-^{12}C_{UL})$ | $\Delta\tilde{\nu}(^{12}C_L-^{12}C_{UL})$ | $I(^{13}C)$ | $I(^{12}C)$ | $I(^{13}C)/I(^{12}C)$ | $I(^{12}C_L)/I(^{12}C_{UL})$ |
|---|---|---|---|---|---|---|---|---|
| Unlabeled | 1615.0 | 1634.4 | | | 0.4 | 23.1 | | |
| 1, 11, ...71 | 1615.0 | 1634.1 | | -0.3 | 5.2 | 19.8 | | 86 |
| 2, 12, ...72 | 1602.9 | 1634.7 | -31.5 | 0.3 | 5.3 | 16.7 | 32 | 72 |
| 3, 13, ...73 | 1604.4 | 1636.5 | -30.0 | 2.1 | 6.7 | 13.8 | 48 | 60 |
| 4, 14, ...74 | 1605.1 | 1637.6 | -29.3 | 3.2 | 6.6 | 15.9 | 42 | 69 |
| 5, 15, ...75 | 1604.7 | 1638.3 | -29.7 | 3.9 | 7.0 | 16.4 | 43 | 71 |
| 6, 16, ...76 | 1605.2 | 1637.5 | -29.2 | 3.1 | 6.4 | 15.0 | 43 | 65 |
| 7, 17, ...77 | 1604.7 | 1635.7 | -29.7 | 1.3 | 6.4 | 14.3 | 45 | 62 |
| 8, 18, ...78 | 1605.0 | 1634.5 | -29.4 | 0.1 | 4.8 | 17.2 | 28 | 75 |
| 9, 19, ...79 | 1615.0 | 1633.6 | | -0.8 | 2.9 | 21.4 | | 93 |
| Average IE (3-7) | | | -29.6 | 2.7 | | | 44 | 65 |

The first two and the last labeled residues are indicated in the left column: for example "1, 11, ...71" means that residues 1, 11, 21, 31, 41, 51, 61, and 71 were labeled. See Table S1 for further explanation.

**Table S12**. Spectral parameters for an isotope-labeled **parallel** β-sheet with **8** strands and **10** residues (9 amide groups) per strand as shown in the bottom panel of Figure S4. **All** strands contained a labeled amide group that were **out-of-register** because each following strand is shifted by one residue towards the N-terminus of the previous strand.

| Label positions | $\tilde{\nu}(^{13}C)$ | $\tilde{\nu}(^{12}C)$ | $\Delta\tilde{\nu}(^{13}C-^{12}C_{UL})$ | $\Delta\tilde{\nu}(^{12}C_L-^{12}C_{UL})$ | $I(^{13}C)$ | $I(^{12}C)$ | $I(^{13}C)/I(^{12}C)$ | $I(^{12}C_L)/I(^{12}C_{UL})$ |
|---|---|---|---|---|---|---|---|---|
| Unlabeled | 1615.0 | 1634.2 | | | 0.4 | 22.8 | | |
| 1, 11, ...71 | 1615.0 | 1634.1 | | -0.1 | 5.6 | 19.2 | | 84 |
| 2, 12, ...72 | 1602.6 | 1634.8 | -31.6 | 0.6 | 5.0 | 16.3 | 30 | 72 |
| 3, 13, ...73 | 1604.6 | 1636.3 | -29.6 | 2.1 | 6.3 | 13.8 | 45 | 60 |
| 4, 14, ...74 | 1605.1 | 1637.4 | -29.1 | 3.2 | 6.4 | 15.9 | 41 | 70 |
| 5, 15, ...75 | 1604.6 | 1638.1 | -29.6 | 3.9 | 6.7 | 16.1 | 42 | 70 |
| 6, 16, ...76 | 1605.2 | 1637.3 | -29.0 | 3.1 | 6.3 | 14.8 | 42 | 65 |
| 7, 17, ...77 | 1604.4 | 1635.7 | -29.8 | 1.5 | 6.0 | 14.0 | 43 | 61 |
| 8, 18, ...78 | 1604.6 | 1634.3 | -29.6 | 0.1 | 4.5 | 17.1 | 26 | 75 |
| 9, 19, ...79 | 1615.0 | 1633.8 | | -0.4 | 3.3 | 21.2 | | 93 |
| Average IE (3-7) | | | -29.4 | 2.8 | | | 43 | 65 |

The first two and the last labeled residues are indicated in the left column: for example "1, 11, ...71" means that residues 1, 11, 21, 31, 41, 51, 61, and 71 were labeled. See Table S1 for further explanation.

**Table S13**. Spectral parameters for the isotope-labeled β-barrel section of **α-hemolysin** (pdb code 7AHL)[12] consisting of 7 hairpins with 19 residues each as shown in Figure S5. Every β-hairpin (every **second** strand) contained one label.

| Label positions | $\tilde{\nu}(^{13}C)$ | $\tilde{\nu}(^{12}C)$ | $\Delta\tilde{\nu}(^{13}C-^{12}C_{UL})$ | $\Delta\tilde{\nu}(^{12}C_L-^{12}C_{UL})$ | $I(^{13}C)$ | $I(^{12}C)$ | $I(^{13}C)/I(^{12}C)$ | $I(^{12}C_L)/I(^{12}C_{UL})$ |
|---|---|---|---|---|---|---|---|---|
| Unlabeled | 1625.0 | 1643.1 | | | 1.9 | 21.9 | | |
| 120.0 | 1625.0 | 1641.0 | | -2.1 | 3.4 | 20.6 | | 94 |
| 121 | 1608.9 | 1644.6 | -34.2 | 1.5 | 4.8 | 17.0 | 28 | 78 |
| 122 | 1615.4 | 1648.4 | -27.7 | 5.3 | 5.9 | 15.9 | 37 | 72 |
| 123 | 1609.5 | 1647.7 | -33.6 | 4.6 | 5.1 | 17.8 | 29 | 81 |
| 124 | 1613.8 | 1644.8 | -29.3 | 1.7 | 5.3 | 18.3 | 29 | 83 |
| 125 | 1625.0 | 1643.9 | | 0.8 | 5.4 | 17.3 | | 79 |
| 126 | 1617.6 | 1643.1 | -25.5 | 0.0 | 3.9 | 18.9 | 21 | 86 |
| 127 | 1625.0 | 1643.2 | | 0.1 | 3.9 | 20.7 | | 95 |
| 128 | 1625.0 | 1643.1 | | 0.0 | 1.9 | 22.0 | | 100 |
| 129 | 1625.0 | 1642.9 | | -0.2 | 3.1 | 21.9 | | 100 |
| 130 | 1625.0 | 1643.0 | | -0.1 | 2.0 | 23.1 | | 105 |
| 131 | 1625.0 | 1642.5 | | -0.6 | 3.3 | 21.9 | | 100 |
| 132 | 1602.8 | 1644.0 | -40.3 | 0.9 | 2.9 | 18.8 | 15 | 86 |
| 133 | 1625.0 | 1643.6 | | 0.5 | 4.9 | 19.5 | | 89 |
| 134 | 1619.3 | 1645.8 | -23.8 | 2.7 | 4.7 | 19.0 | 25 | 87 |
| 135 | 1622.7 | 1646.7 | -20.4 | 3.6 | 8.1 | 16.2 | 50 | 74 |
| 136 | 1625.0 | 1645.8 | | 2.7 | 5.9 | 17.7 | | 81 |
| 137 | 1625.0 | 1644.7 | | 1.6 | 7.7 | 16.5 | | 75 |
| Average IE (121-124, 134-135) | | | -28.2 | 2.7 | | | 33 | 79 |

The loops between the β-strands are included. The same residue in each monomer was labeled. See Table S1 for further explanation.

**Table S14**. Spectral parameters for an isotope-labeled **helix-sheet** model as shown in Fig. S7. A **single** label was placed either in the helix or the sheet. Chain A is the helix and chain B the antiparallel β-sheet.

| Label positions | $\tilde{\nu}(^{13}C)$ | $\tilde{\nu}(^{12}C)$ | $\Delta\tilde{\nu}(^{13}C-^{12}C_{UL})$ | $\Delta\tilde{\nu}(^{12}C_L-^{12}C_{UL})$ | $I(^{13}C)$ | $I(^{12}C)$ | $I(^{13}C)/I(^{12}C)$ | $I(^{12}C_L)/I(^{12}C_{UL})$ |
|---|---|---|---|---|---|---|---|---|
| Unlabeled | 1610.0 | 1625.9 | | | 0.5 | 10.0 | | |
| A9 | 1610.0 | 1625.8 | | -0.1 | 0.9 | 10.1 | | 101 |
| A13 | 1610.0 | 1625.9 | | 0.0 | 0.9 | 10.2 | | 102 |
| B19 | 1610.0 | 1625.8 | | -0.1 | 1.1 | 9.6 | | 96 |
| B20 | 1607.2 | 1627.3 | -18.7 | 1.4 | 2.0 | 8.1 | 24 | 81 |
| B21 | 1599.1 | 1631.6 | -26.8 | 5.7 | 1.9 | 9.9 | 19 | 99 |
| B22 | 1607.2 | 1629.2 | -18.7 | 3.3 | 2.3 | 8.0 | 28 | 80 |
| B23 | 1610.0 | 1625.6 | | -0.3 | 1.8 | 9.1 | | 91 |
| B27 | 1600.0 | 1631.9 | -25.9 | 6.0 | 2.0 | 9.7 | 20 | 98 |
| Average IE (B20-B22) | | | -21.4 | 3.5 | | | 24 | 87 |

See Table S1 for further explanation.
